# Embedded transport accelerates interaction-limited biosensing

**DOI:** 10.64898/2026.06.18.733113

**Authors:** Andrea Dsouza, Yuling Yang, Thomas S. Davies, Julia Brettschneider, David M. Haddleton, Rachel A. Hand, Natasha Ratnaraja, Meera Unnikrishnan, Chrystala Constantinidou, Jérôme Charmet

**Affiliations:** Division of Biomedical Sciences, Warwick Medical School, The University of Warwick, Coventry, CV4 7AL, United Kingdom; Department of Statistics, The University of Warwick, Coventry, CV4 7AL, United Kingdom; Department of Chemistry, The University of Warwick, CV4 7AL, United Kingdom; University Hospitals Coventry & Warwickshire, Coventry, CV2 2DX; Bioinformatics Research Technology Platform, The University of Warwick, CV4 7AL, United Kingdom; School of Engineering - HE-Arc Ingénierie, HES-SO University of Applied Sciences Western Switzerland, 2000 Neuchâtel, Switzerland; School of Biomedical and Precision Engineering, University of Bern, Güterstrasse 24/26, 3008 Bern, Switzerland

## Abstract

Phenotypic biosensors that measure bacterial viability and antimicrobial susceptibility are essential for rapid infectious disease diagnostics, yet their speed is fundamentally limited by the rate at which bacteria encounter reporter molecules, a transport bottleneck that has been typically addressed by complex microfluidic solutions. Here we show that this bottleneck can be overcome by engineering transport directly into the sensing material. A multifunctional ionic hydrogel matrix, co-encapsulating bacterial growth medium and the redox reporter resazurin, exploits swelling-driven convective transport to dramatically accelerate bacteria-reporter interactions without any change to assay chemistry. By systematically tailoring the hydrogel crosslinking density and optimizing the encapsulated nutrient-osmotic microenvironment, we maximize metabolic signal generation to achieve a 12-to-48-fold reduction in detection time relative to solution-phase and conventional hydrogel assays. Deployed in a standard 96-well format for urinary tract infection (UTI) diagnosis of 48 clinical samples, the platform rapidly detects infection in 15 minutes to 2 hours, achieving 95% sensitivity and 100% specificity for bacterial detection, and 100% sensitivity and 98% specificity for antimicrobial susceptibility profiling, compared to time-consuming gold-standard urine culture-based methods. Results are readable both quantitatively on a plate reader and visually as a colorimetric assay, enabling point-of-care deployment without additional instrumentation. Thus, embedding transport enhancement within the sensing matrix, represents a general and scalable design principle for accelerating interaction-limited biosensing, which has excellent scope for rapid diagnostic development.

## Introduction

Biosensors that measure bacterial viability and antimicrobial susceptibility (AST) are vital tools in combating the global threat of antimicrobial resistance (AMR), yet their speed is fundamentally limited by a critical mass transport bottleneck^1–6^. Phenotypic testing remains essential to capture the metabolic states and non-genetic resistance mechanisms that sequencing cannot predict ^7,8^. Consequently, overcoming this mass transport limitation is a critical parameter that must be prioritized in biosensor design. In the static or minimally mixed formats typical of clinical microbiology, such as resazurin metabolic assays, chromogenic identification media, and broth microdilution panels, these interactions rely almost entirely on diffusion^9^. Even though the encapsulated reporter molecules vastly outnumber the target bacteria, these assays frequently fail to generate a rapid signal. This delayed timescale stems not from a lack of chemical sensitivity, but rather from transport limitations that hinders physical interactions between bacteria and reporters. This reliance on diffusion-limited transport stretches detection timescales to hours or days, a delay epitomized by urinary tract infections (UTIs)^10–12^, which are among the most common bacterial infections worldwide. As current gold-standard urine cultures take up to 48 hours to deliver results, clinicians are left with no option but to prescribe empirical, broad-spectrum antibiotics to manage acute patient symptoms, creating a significant diagnostic gap which promotes the development of resistance^13,14^. Developing new sensitive technologies that overcome transport bottlenecks is therefore paramount to accelerating diagnostics and enabling targeted, timely clinical decision-making.

The standard engineering response to this transport bottleneck is to impose flow from outside the assay via microfluidic channels, active mixing and forced convection^13,15,16^. These approaches work, but they decouple transport control from the sensing chemistry, add hardware complexity, and may limit deployment at the point of care^10–12^. A more elegant solution is to embed transport enhancement within the sensing environment itself, enabling the material to perform the role of microfluidics without moving parts or external infrastructure. In this work, we present an autonomous transport-enhanced diagnostic platform based on an ionic hydrogel matrix. Utilizing a 2-acrylamido-2-methylpropane sulfonic acid sodium salt (NaAMPS) hydrogel co-encapsulating growth medium and the redox reporter resazurin, we leverage the material’s intrinsic swelling dynamics to drive convective transport, accelerating bacteria-reporter interactions without altering the underlying assay chemistry. We systematically investigate the interplay between hydrogel network density and encapsulated nutrient microenvironment to establish the fundamental design rules for minimizing detection time. Finally, we validate this platform using clinical urine samples in a standard 96-well format for rapid urinary tract infection (UTI) screening and antimicrobial susceptibility testing (AST). Capable of generating both quantitative and visual colorimetric readouts, this approach establishes hydrogel-mediated transport as a new paradigm in for biosensor design, shifting transport control from external infrastructure directly into the material itself.

## Results

### Hydrogel embedding enables transport-mediated acceleration of bacterial detection

Phenotypic bacterial detection is fundamentally constrained by the rate of bacterial growth^18,19^; however, in reporter-based assays, transport-limited interactions between bacteria and reporter molecules can impose an additional bottleneck. To test this, we compared resazurin-based assays performed in conventional liquid culture and multifunctional, swelling NaAMPS hydrogel matrix (Fig. 1A). As illustrated in Fig. 1A, solution-phase assays rely predominantly on diffusion to mediate bacteria-resazurin interactions, whereas the dynamic NaAMPS hydrogel matrix enhances transport through swelling-driven redistribution within a confined network.

**Figure 1.**
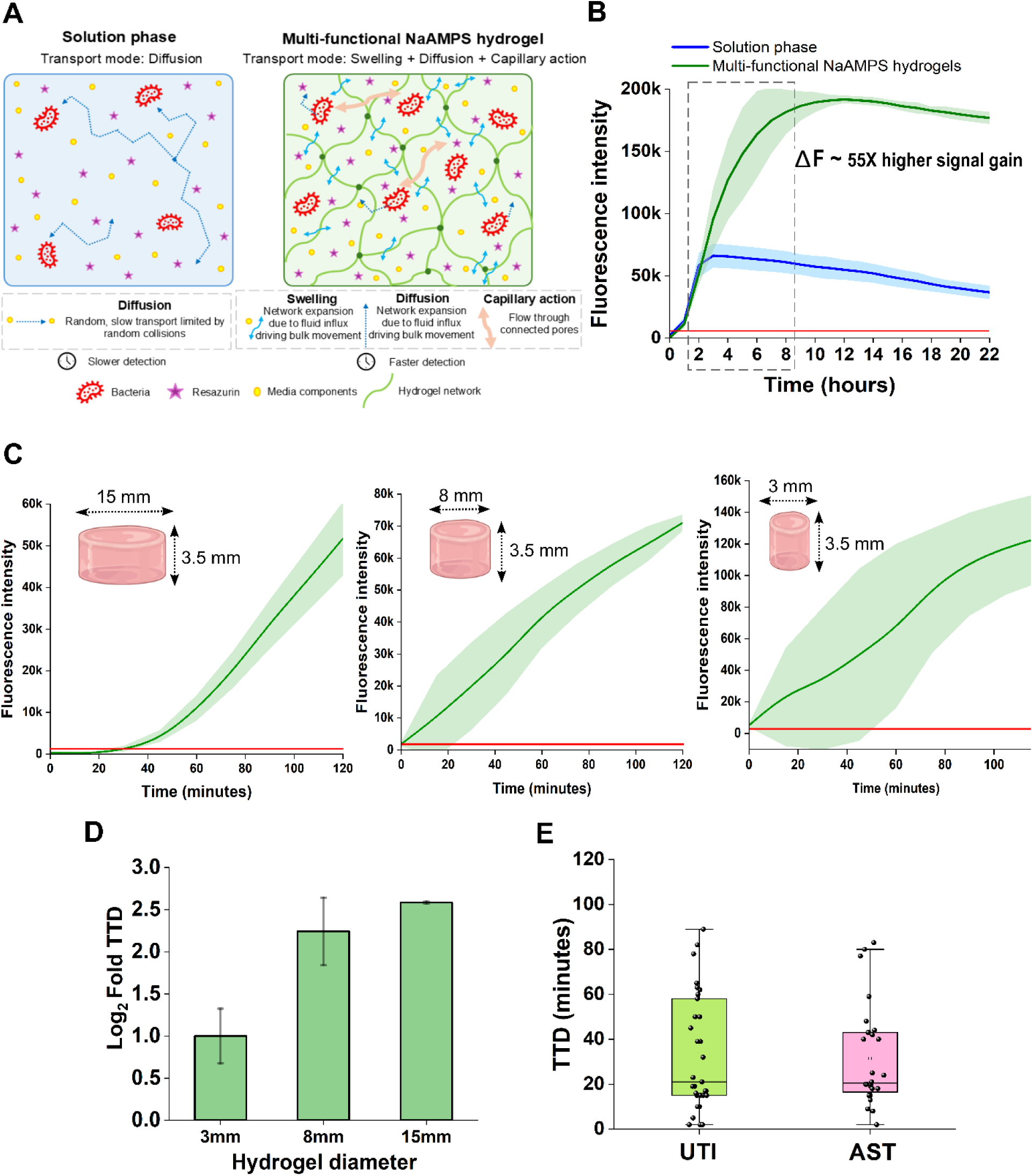
Hydrogel embedding enhances bacteria-reporter interactions and enables transport-mediated acceleration of detection. A) Schematic of transport mechanisms in conventional solution phase and multifunctional NaAMPS hydrogel systems. In solution, bacteria-resazurin encounters are governed by diffusion, resulting in slow and spatially heterogenous interactions. In the hydrogel, swelling-driven solvent flow, diffusion, and capillary action collectively enhance transport within a confined polymer network, promoting more efficient redistribution of bacteria and reporter molecules. B) Comparison of bacterial detection kinetics in solution and hydrogel-embedded assays. Solution phase assays rapidly plateau at low intensity, whereas hydrogel systems show sustained signal accumulation. C) Detection kinetics in NaAMPS hydrogels of varying diameters (15, 8, and 3 mm). Reducing gel size leads to faster fluorescence accumulation and earlier detection, consistent with transport length-scale effects. D) Log_2_ fold-change in TTD as a function of hydrogel diameter. Detection time decreases monotonically with decreasing size, demonstrating that transport length scale governs assay kinetics. E) TTD across 48 clinical urine samples for UTI detection and AST. Each point represents an individual sample; box plots indicate median and interquartile range. Hydrogel-based assays achieve rapid detection across a broad range of clinical conditions.

Both the solution and dynamic hydrogel system exhibit a similar onset of fluorescence (Fig. 1B), indicating comparable initial bacteria-resazurin encounter kinetics and detection thresholds. However, their kinetic trajectories diverge immediately thereafter. In solution and non-swelling hydrogels, fluorescence increases rapidly but plateaus at lower intensity, consistent with limited replenishment of substrate within the local microenvironment and transport-limited redistribution. In contrast, the NaAMPS hydrogel supports sustained signal accumulation, achieving approximately 55-fold higher signal gain (Δ*F*). To mathematically characterize these distinct kinetic regimes, we fitted the trajectories to a classical Gompertz growth model (Supplementary Fig. S1). While the solution-phase assays completely deviate from sigmoidal growth behaviour due to rapid local substrate depletion, yielding poor fits (*R*^2^ < 0) and a specific signal accumulation rate (*µ*) of zero, the NaAMPS hydrogel curves demonstrate a robust sigmoidal fit (*R*^2^ > 0.98) with a well-defined, reproducible accumulation rate (*µ* = 0.699 ± 0.277). This mathematical divergence highlights that the hydrogel matrix fundamentally alters the reaction kinetics, transitioning the system away from a transport-limited plateau.

To evaluate the physical constraints driving this divergence, we analysed the transport kinetics within the initial linear regime (*t* = 1.5 to 3.5 hours) using a classical two-dimensional (2D) diffusion framework governed by 2*Dt*. Scaling the fluorescence trajectory against a square-root-of-time dependency (*ΔF* ∝ √*t*), yielded an apparent transport rate inside the hydrogel network (*D*_app_) that outpaces the solution phase-diffusion by ∼4.45-fold (*D*_gel_ ≈ 4.45 *D*_sol_). Because structural polymer networks typically restrict passive molecular mobility through steric hinderance and tortuosity^20–22^, this rapid transport scaling cannot be reconciled with classical passive diffusion alone. Instead, we hypothesize that the dynamic, swelling-driven expansion and intrinsic capillary forces of the NaAMPS hydrogel introduce an active, matrix-mediated transport component that bypasses the passive 2*Dt* diffusion bottleneck. While direct visualization of internal advective flow profiles remains challenging, this transport-enhanced model is strongly supported by the macro-scale geometric dependence of the system. The absence of early saturation in the NaAMPS hydrogel indicates prolonged maintenance of bacteria-resazurin interaction conditions beyond the initial detection event, pointing to a matrix-mediated modulation of transport that prevents the rapid local substrate depletion and equilibration typical of purely diffusion-limited regimes.

To determine whether this behaviour is governed by transport length scale, we varied hydrogel geometry by fabricating discs of defined diameters (15, 8, and 3 mm). Reducing gel size led to progressively faster kinetics, with smaller constructs exhibiting steeper trajectories and earlier detection (Fig 1C). Detection time decreased monotonically with decreasing diameter (Fig. 1D), following a log_2_ scaling with system size. This dependence is consistent with a transport-limited regime in which encounter rates are set by the distance over which bacteria and reporter must traverse. Thus, the sustained kinetics observed in bulk hydrogels (Fig. 1B) can be converted into faster detection by reducing transport length scales.

We next evaluated performance in clinical samples by measuring time to detection (TTD) across 48 urine specimens for urinary tract infections (UTI) and antimicrobial susceptibility testing (AST). Hydrogel-embedded assays yielded rapid detection across a broad distribution of samples, with TTD spanning ∼2 to 90 minutes for both applications (Fig. 1E). Despite biological variability, median detection times remained tightly clustered, indicating robust transport-enhanced bacteria-resazurin interactions across heterogeneous clinical conditions. These timescales represent a substantial compression of conventional assay durations and demonstrate that embedding transport functionality within the sensing matrix enables rapid phenotypic readout in clinically relevant settings. Therefore, these results identify sustained reporter-cell interaction, rather than initial encounter rate, as the dominant transport limitation in conventional assays.

### Hydrogel composition and structure govern the functional sensing environment

Having established that hydrogel-mediated transport enhances bacteria-reporter interactions, we next sought to determine how material composition and network structure define this functional sensing environment. We hypothesized that hydrogel performance is governed by a combination of swelling capacity, charge density, and network architecture, which together regulate both transport dynamics and bacterial accessibility to reporter molecules. We first characterized the swelling behaviour of NaAMPS hydrogels as a function of time and environmental conditions. NaAMPS exhibited rapid and extensive swelling, reaching >600% volumetric expansion within 120 minutes (Fig. 2A) and continuing up to >2000% over 6 hours (Supplementary Fig S2 (i)), accompanied by significant macro-scale geometric expansion (Supplementary Fig S2 (ii)). This swelling behaviour was strongly dependent on pH, with maximum expansion observed under neutral conditions (4 cm diameter) compared to basic (3 cm) and acidic (1.5 cm) conditions (Supplementary Fig S2 (iii)). This is consistent with ionization of sulfonate groups and the resulting increase in osmotic driving force under neutral to basic conditions. (Fig. 2B). Given that clinical urine specimens span a wide physiological spectrum (pH 4.5 to 8.0), characterizing this sensitivity is vital for diagnostic robustness in UTI applications. While extreme acidity partially restricts maximum swelling, the matrix maintains a highly functional baseline hydration. Crucially, because the hydrogel is pre-encapsulated with Mueller Hinton Broth 2 (MHB-II), a standardized medium natively formulated at pH 7.3 ± 0.2, the network possesses an intrinsic buffering capacity. Upon sample introduction, this pre-loaded environment normalizes pH deviations, ensuring the NaAMPS matrix operates near its optimal swelling window to yield reproducible transport and bacterial detection kinetics regardless of the pH value. These results confirm that NaAMPS generates substantial solvent influx, providing a physical basis for enhanced convective transport within the matrix.

**Figure 2.**
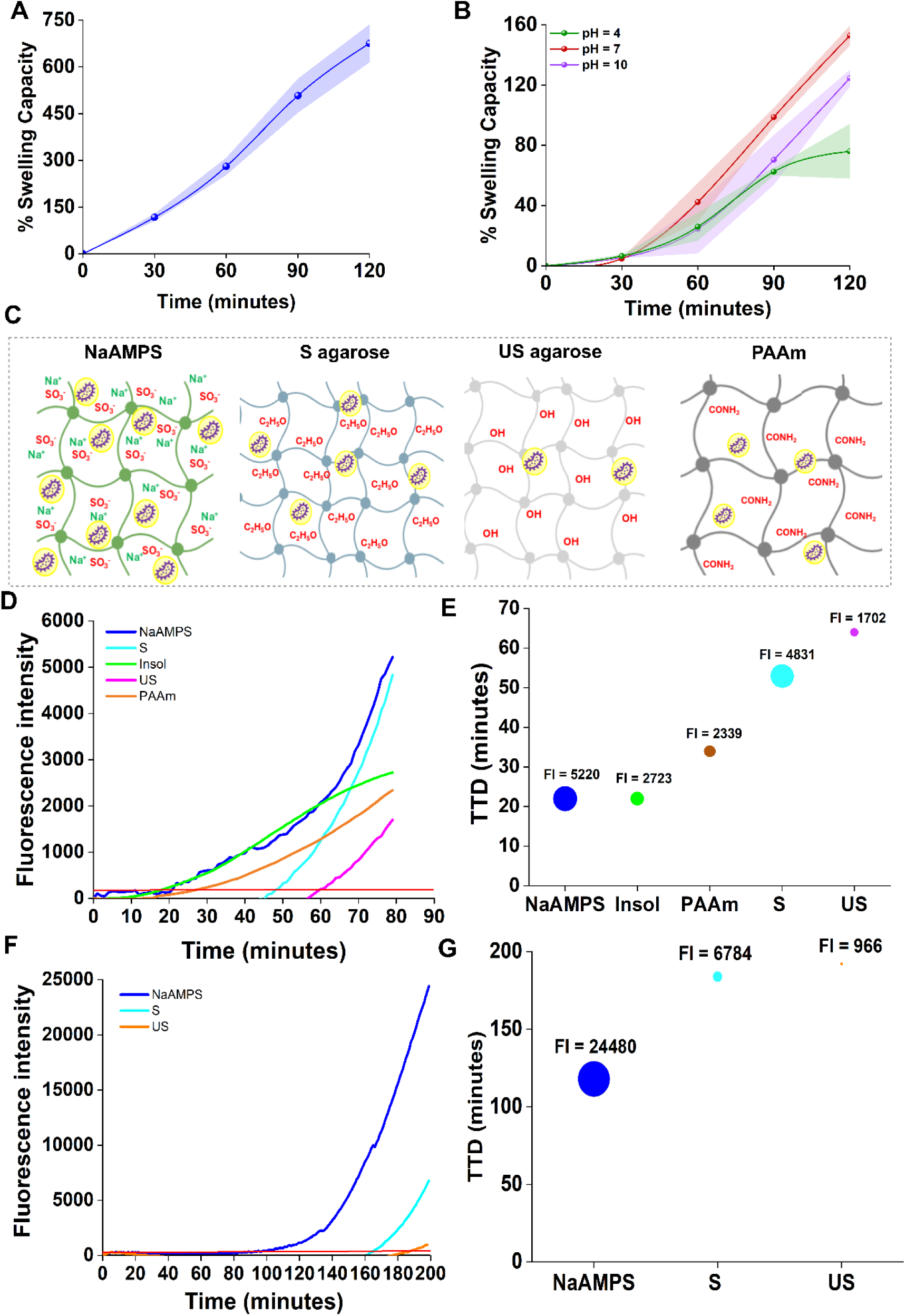
Physicochemical characterization and transport-dependent sensing performance of hydrogel matrices. A) Swelling kinetics of NaAMPS hydrogel matrix (n=3) over 120 minutes, demonstrating rapid volumetric expansion exceeding 600%. B) Influence of environmental pH on NaAMPS swelling capacity; maximum expansion occurs at neutral to basic pH (pH 7-10), driven by the ionization of sulfonate groups. C) Schematic representations of the chemical architecture and proposed bacterial interaction modes for ionic (NaAMPS), hydrophilic S agarose, neutral US agarose, and PAAm hydrogels. (D to G) Comparative bacterial detection kinetics across different material platforms. Fluorescence intensity over time for 10^7^ CFU/mL (D), 10^5^ CFU/mL (F) and corresponding TTD metrics (E) and (G). (E-G) reveal that NaAMPS significantly outperforms non-swelling or moderately swelling matrices, establishing a link between material-driven convective transport and accelerated sensing. Bubble size in (E) and (G) represents total fluorescence intensity (FI) at the end point.

To systematically probe the role of material properties, we compared NaAMPS to a panel of hydrogels spanning distinct swelling capacities and network structures, including hydroxyethylated agarose (S), non-hydroxyethylated agarose (US), and polyacrylamide (PAAm) (Fig. 2C). These materials provide a controlled framework to decouple the contributions of swelling, hydration, and structural accessibility. At a high bacterial concentration (10^7^ CFU/mL *E. coli*), detection kinetics varied markedly across the difference materials (Fig. 2D,E). To quantify these differences, we evaluated the Time-to-Detection (TTD) and the corresponding maximum Fluorescence intensity (FI). The speed of detection followed the order of NaAMPS > Insol > PAAm > S > US. NaAMPS hydrogels exhibited a significantly reduced TTD (∼22 minutes) paired with a high FI = 5220, demonstrating accelerated signal generation. This kinetic trend reflects the efficiency with which bacteria-resazurin interactions are established and sustained within each matrix. NaAMPS achieved the most rapid detection, consistent with its structural capacity to support multimodal mass transport including swelling-driven solvent influx, passive diffusion, and capillary flow. Together, this promotes the rapid, continuous redistribution of reporter molecules toward metabolically active bacteria. In contrast, materials lacking significant swelling constrained bacteria-resazurin interactions. Within the agarose cohorts, S agarose hydrogels outperformed US agarose hydrogels^23^, confirming that increased network hydration and structural accessibility enhance the frequency and uniformity of these metabolic interactions. At a lower bacterial concentration (10^5^ CFU/mL), the performance gap between the materials became even more pronounced (Fig 2F, G). NaAMPS maintained robust and rapid sensing capabilities (TTD ∼ 118 minutes, FI = 24480). Conversely, S agarose lagged significantly (TTD ∼ 185 minutes), and US agarose yielded an extremely weak, delayed signal.

Notably, PAAm exhibited a distinct divergence in performance. Despite possessing moderate swelling capabilities, its detection kinetics were considerably slower at high concentrations, and it failed to produce a viable signal within the tested timeframe at lower bacterial concentrations. This divergence suggests that volumetric swelling alone is insufficient to sustain optimal bacteria-resazurin interactions; rather, transport efficiency within the hydrogel is heavily dictated by polymer charge density. PAAm lacks the highly dense, fixed negative sulfonate groups present in the NaAMPS backbone that drive strong electrostatic swelling pressure and govern matrix-ion interactions^17,24^. Consequently, the PAAm network structure likely imposes greater diffusional resistance, restricting the effective mobility and spatial redistribution of bacteria or reporter molecules within the matrix. These findings highlight that tuning the ionic content and charge profiles of the hydrogel matrix, rather than optimizing swelling capacity alone is essential to engineering an optimised functional sensing environment.

To define how network architecture shapes the sensing environment, we varied the crosslinking density of NaAMPS hydrogels by tuning PEGDA concentration and visualised bacterial growth using GFP-expressing *E. coli* (Fig. 3A). Low crosslinking density (0.1 mM), hydrogels resulted in structurally unstable networks and heterogeneous bacterial infiltration, whereas high densities (≥0.5 mM) imposed excessive physical confinement, forcing confluent, overlapping bacterial populations (Supplementary Fig. S3). Optimizing the architecture at an intermediate crosslinking density (0.2 mM) yielded physically stable networks that maintained localized, discrete colony domains. This structural balance is critical; intermediate confinement preserves both nutrient/reporter accessibility and spatial organization, enabling controlled sustained bacteria-resazurin interactions.

**Figure 3.**
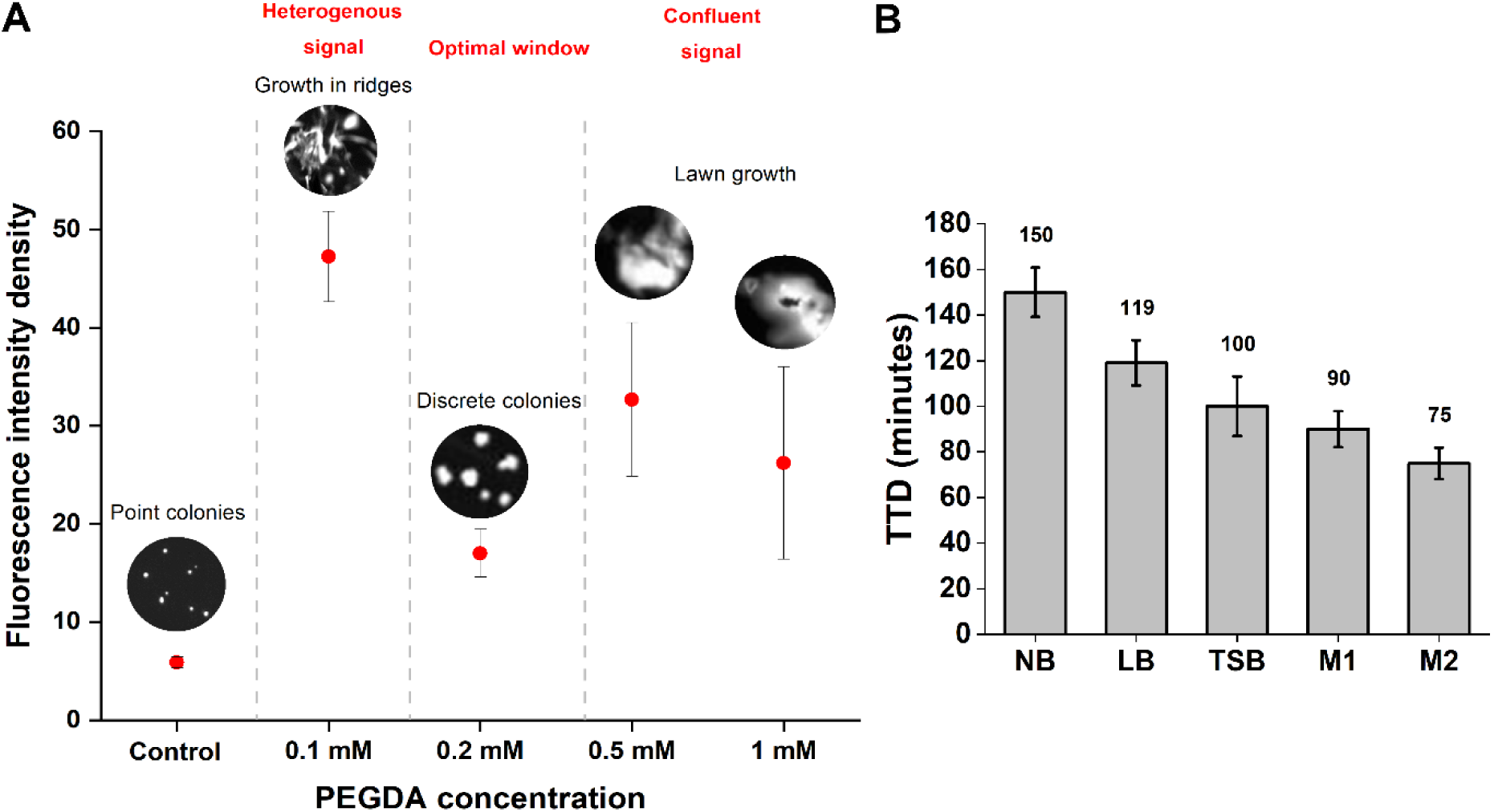
Optimization of network architecture and chemical microenvironment for enhanced sensing. A) Effect of crosslinking density (PEGDA concentration) on the spatial organization of bacterial growth and signal homogeneity. Representative fluorescence images of GFP-expressing *E. coli* show a transition from heterogeneous growth at low density (0.1 mM) to discrete, stable colonies within an optimal window (0.2 mM), and finally confluent, transport-restricted lawn growth at higher densities (0.5-1.0 mM). B) Modulation of sensing kinetics by the encapsulated chemical environment. TTD values for 5 different growth media (NB, LB, TSB, M1, and M2) demonstrate that M2 medium provides the fastest detection (∼75 min), highlighting the synergy between matrix architecture and local nutrient/ionic composition in facilitating bacteria-reporter encounters.

We next examined how the chemical environment modulates hydrogel performance by encapsulating various growth media within the optimized matric (Fig. 3B-C). Detection kinetics varied substantially with media composition; an M2 (Mueller Hinton Broth – 2) formulation achieved the fastest time-to-detection (∼75 min), driven by a minimized lag phase (t_lag_ = 0.954 ± 0.024 hours) and the highest specific signal accumulation rate (µ = 0.915 ± 0.076 h^-1^) (Fig. 3C, Supplementary Fig. S4, Table S1). This accelerated profile may be modulated by divalent ions within the M2 media that network architecture and environmental composition jointly define the functional sensing environment by regulating the spatial distribution and persistence of bacteria-reporter interactions.

Together, these results demonstrate that hydrogel performance arises from the coupling of transport, network structure, and environmental composition. Efficient bacterial detection requires not only swelling-driven transport, but also a network architecture that maintains spatial accessibility and a chemical microenvironment that supports sustained bacteria-resazurin interactions. By tuning these parameters, the hydrogel can be engineered to regulate both the spatial organisation of bacterial activity and persistence of reporter-cell interactions. This establishes the sensing matrix itself as a controllable microenvironment, in which transport and interaction dynamics can be systematically optimised to enhance phenotypic detection.

### Performance validation in synthetic urine and antibiotic susceptibility testing

To evaluate the diagnostic performance of the hydrogel sensing platform under clinically relevant conditions, we first assessed its ability to detect uropathogenic bacteria and quantify sensitivity in synthetic urine. As shown schematically in Fig. 4A, the hydrogel matrix co-encapsulates growth nutrients and resazurin, enabling metabolic activity to be transduced into a visible colour change and a corresponding increase in fluorescence intensity upon bacterial growth. This dual-readout configuration allows for both high-throughput quantitative tracking via a plate reader and immediate, instrument-free binary classification by the naked eye. An encapsulated resazurin concentration of 1 µg/mL was determined to be optimal for the diagnostic formulation, prioritizing the fastest onset kinetics while maintaining an entirely unambiguous signal window. Systematic evaluation across a range of reporter concentrations (1 to 5 µg/mL; Supplementary Fig. S5) revealed a complex interplay between early analytical clarity, kinetic velocity, and terminal signal capacity. While an elevated concentration of 5 µg/mL resulted in an anomalous, non-sigmoidal bell-shaped trajectory with late-stage signal decay, the analytically viable range (1 to 4 µg/mL) displayed an inverse relationship between resazurin density and the specific signal accumulation rate (µ). Specifically, µ decreased from 0.885 ± 0.092 at 1 µg/mL to 0.557 ± 0.029 at 4 µg/mL, despite non-linear variations in the early signal-to-noise ratio (SNR) at 2 hours (Supplementary Table S2 and S3). Consequently, the 1 µg/mL configuration was selected for all downstream validations to ensure rapid diagnostic turnaround and a reliable baselines SNR (86.58) without the metabolic deceleration typical of higher reporter loads. In the absence of infection, no colour or fluorescence change is observed, confirming a clear on-off response suitable for point-of-care screening.

**Figure 4.**
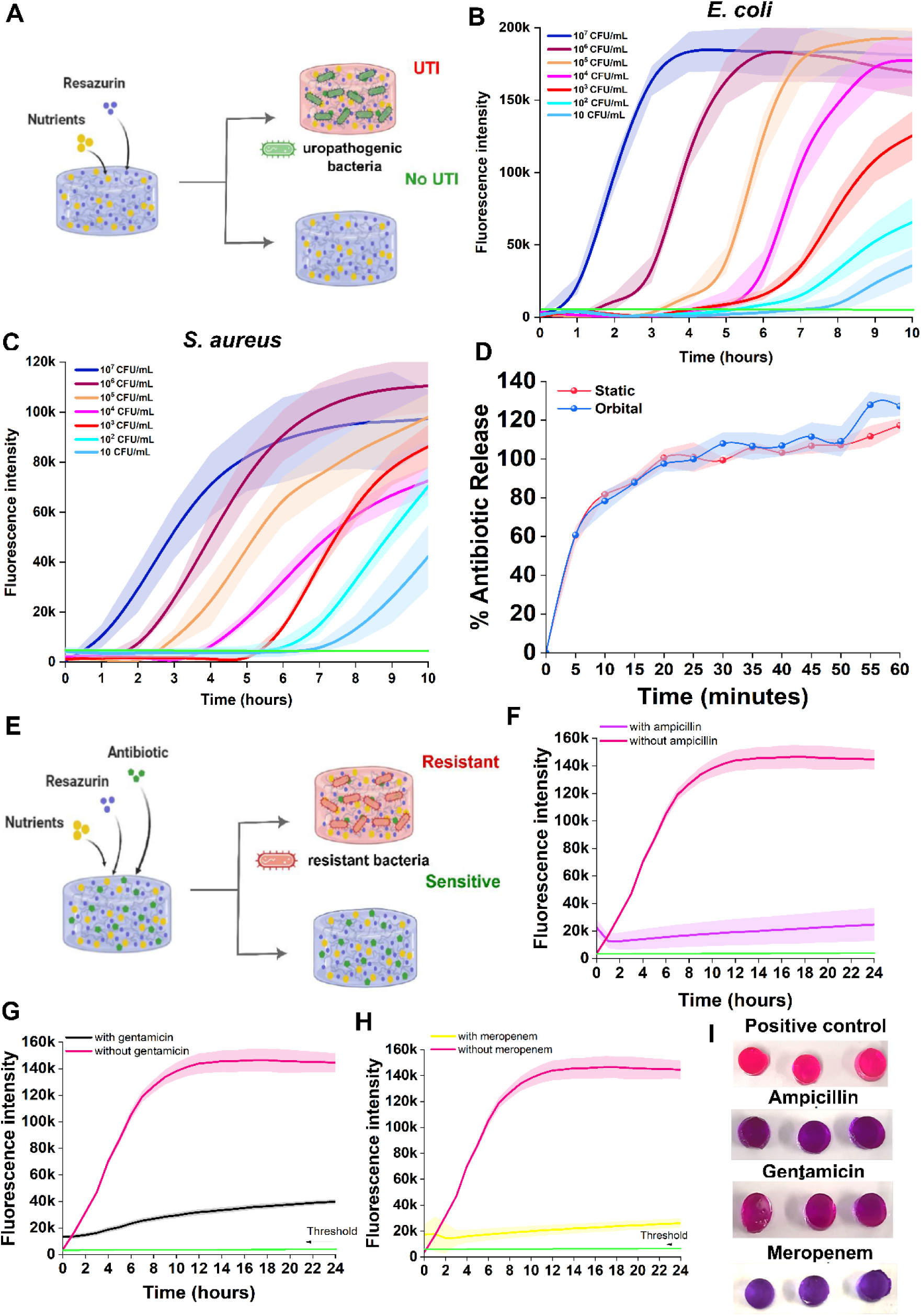
Hydrogel-based bacterial detection and integrated AST. A) Schematic of the hydrogel sensing platform incorporating growth nutrients and resazurin. In the presence of viable bacteria, resazurin to reduced to resorufin producing fluorescence signal and visible colour change, while no signal and colour change are observed in the absence of infection. B) LOD for *E. coli*, demonstrating sensitive and concentration-dependent bacterial detection within the hydrogel matrix. C) LOD for *S. aureus*, confirming applicability across Gram-negative and Gram-positive pathogens. D) Release kinetics of ampicillin from NaAMPS hydrogels under static and orbital shaking conditions. Comparable kinetics observed, with ∼60% release within 5 minutes and complete release by 20 minutes. E) Schematic of the antibiotic-integrated hydrogel for AST. Resistant bacteria retain metabolic activity and generate signal, whereas susceptible bacteria show no response. Detection kinetics of *E. coli* in F) ampicillin-loaded, G) gentamicin-loaded, and H) meropenem-loaded hydrogels. I) Colorimetric visualization of antibiotic susceptibilities.

Quantitative evaluation of detection sensitivity was performed using a 24-well plate format to determine the limits of detection for both Gram-negative and Gram-positive pathogens. For *E. coli* (Fig. 4B), the assay exhibited a robust concentration-dependent response, enabling detection across clinically relevant bacterial loads (≥10^5^ CFU/mL). Similarly, *S. aureus* (Fig. 4C) was detected with comparable sensitivity, indicating that the hydrogel platform is not restricted to a single pathogen class and maintains broad-spectrum applicability for infection screening. Crucially, the distinct colorimetric shift from purple to pink was unambiguously discernible by eye across all positive samples within these clinical thresholds, validating the platform’s suitability for decentralized, instrument-free diagnostic deployment. As demonstrated by the raw multi-well assay profiles for *S. aureus* (Fig. 5A), high-density inoculations (10^7^ to 10^8^ CFU/mL) produced a complete shift to pink, while intermediate clinical thresholds (10^5^ to 10^6^ CFU/mL) yielded transitional purple-pink hues. In separate quantitative evaluations using *E. coli* (Fig. 5B), spectrophotometric detection via inverse absorbance kinetic curves (1/OD 600nm) showed a predictable right-shift in exponential signal as the initial bacterial load decreased. This metabolic readout correlates with macroscopic colony formation on the hydrogel surface, contrasting with the unreduced baseline of negative controls (Supplementary Fig. S6).

**Figure 5.**
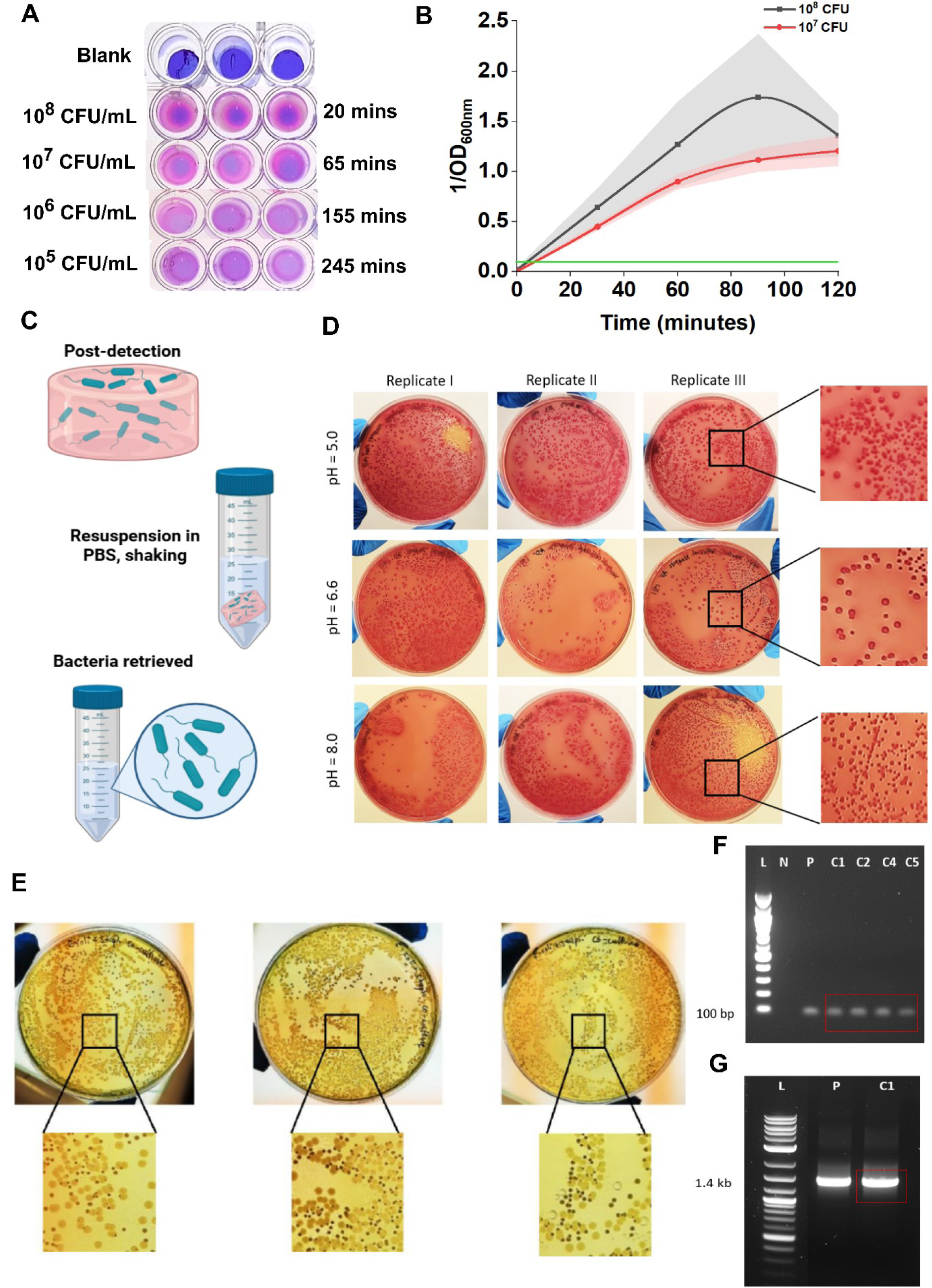
Downstream recovery of viable bacteria for phenotypic and genotypic analyses. A) Visual colorimetric assay for *S. aureus*, demonstrating naked-eye readout for 10^5^ to 10^7^ CFU/mL concentrations. B) Real-time quantitative spectrophotometric detection of *E. coli* (1/OD600nm), C) Schematic illustrating processing following hydrogel-bacterial detection. Post-assay, hydrogels are transferred into 1XPBS and subjected to orbital shaking (110 rpm), enabling release of bacteria grown on and within the hydrogel. D) Recovery of *E.coli* from NaAMPS hydrogels following detection in artificial urine across varying pH conditions (pH 4, 5.5, and 8). Retrieved bacteria were plated on MacConkey agar, confirming preservation of *E. coli* viability and culturability. E) Recovery from mixed-species samples containing *E. coli* and *S. aureus*. Recovered suspensions plated on CLED agar show distinct colony morphologies, enabling differentiation between species. D) Genotypic confirmation of recovered *S. aureus* by colony PCR targeting *esxA* gene (100 bp amplicon). F) Genotypic confirmation of recovered *E. coli* by colony PCR targeting the 16S rRNA gene (∼1.4 kb amplicon).

**Figure 6.**
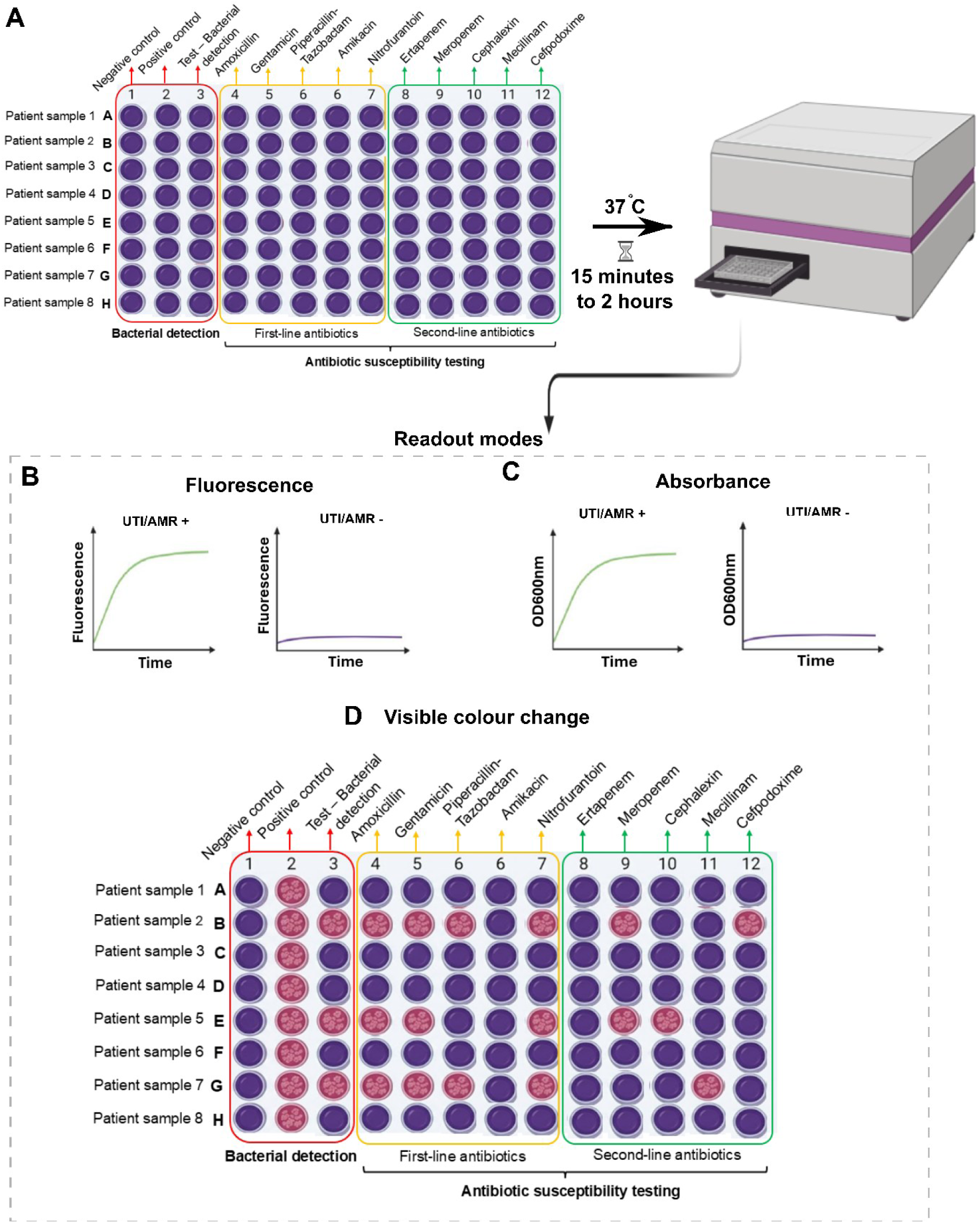
High-throughput screening and multi-mode readout profiles for rapid UTI and AST. **(A)** Schematic workflow of the hydrogel integrated 96-well plate assay configuration for parallel evaluation of eight patient samples (Rows A-H). Columns 1-3 screen for baselines bacterial presence (negative control, positive control, and UTI detection), while columns 4-7 and 8-12 monitor susceptibility against first-line and second-line antibiotic panels, respectively. Assays are incubated at 37°C with readouts captured across accelerated timelines ranging from 15 min to 2 hours. (B-D) Multi-mode diagnostic readout channels. **(B)** Kinetic fluorescence tracking and **(C)** spectrophotometric optical density absorbance kinetic curves (OD600nm) demonstrating distinct signal trajectories between (UTI/AMR +) and negative (UTI/AMR -) samples. **(D)** Colorimetric end-point profiling demonstrating visual metabolic discrimination; active bacterial proliferation drives a metabolic shift from unreduced resazurin (purple) to reduced resorufin (pink), indicating specific antibiotic resistance profiles across the clinical panels.

**Figure 7.**
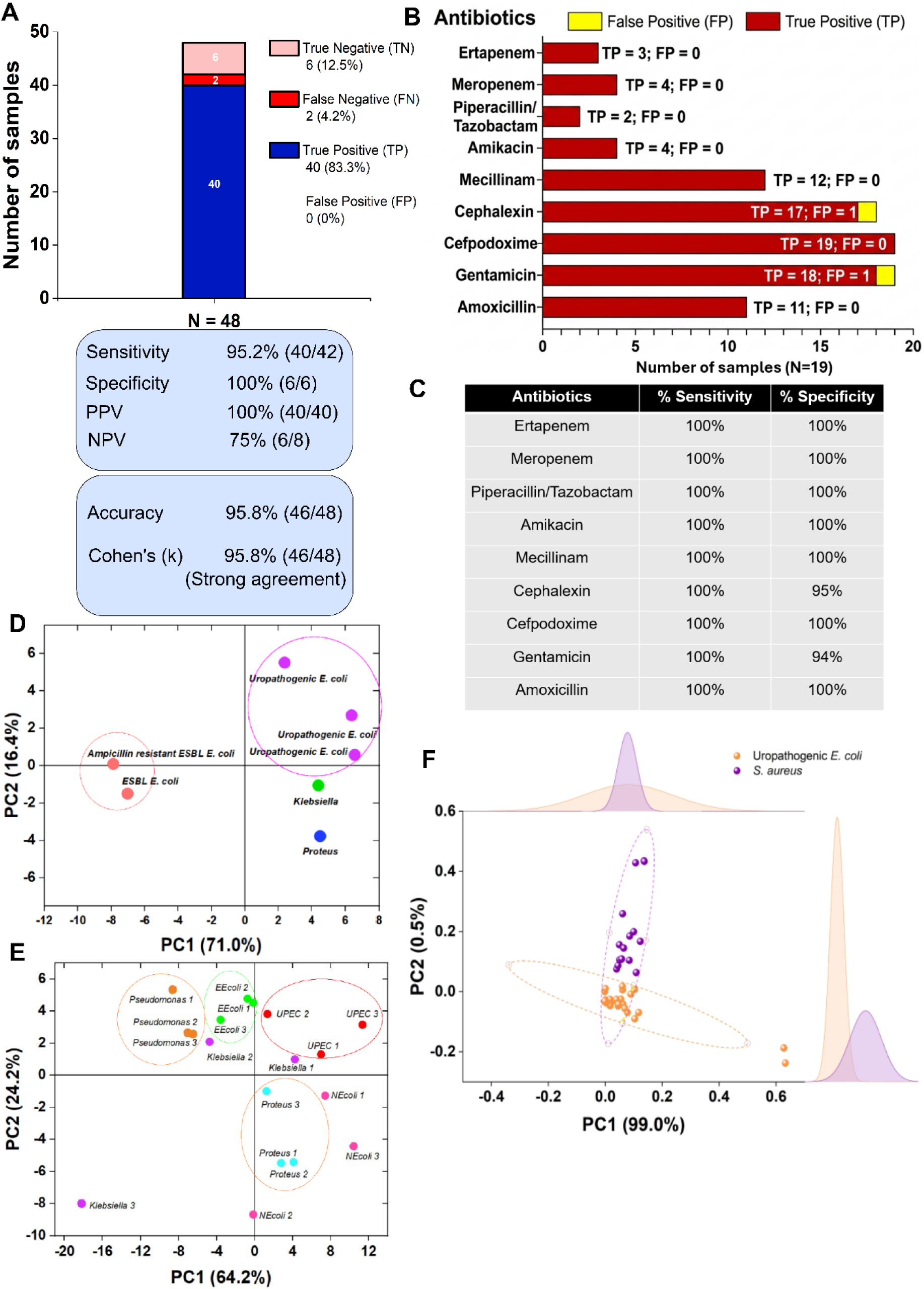
Diagnostic performance and pathogen identification. A) Overall diagnostic evaluation matrix across clinical isolates (N=48). The stacked bar chart categorizes sample outcomes into true positive (TP), false negative (FN), true negative (TN), and false positive (FP) cohorts. Calculated diagnostic parameters include sensitivity, specificity, positive predictive value (PPV), negative predictive value (NPV), total accuracy, and Cohen’s kappa (k) coefficient. (B, C) Susceptibility profiles for individual antibiotics (N=19). (B) Distribution of true positive versus false positive detections across the panel, and (C) corresponding clinical sensitivity and specificity metrics calculated per antibiotic class. (D-F) Principal Component Analysis (PCA) score plots for multi-species and strain-level differentiation based on metabolic signal trajectories. (D) PCA plot demonstrating clustering separation between susceptible uropathogenic *E. coli*, extended-spectrum β-lactamase (ESBL)-producing *E. coli*, Klebsiella, and Proteus species along PC1 (71.0% variance) and PC2 (16.4% variance). (E) Expanded strain-level clustering mapping distinct clinical isolates across diverse bacterial families. (F) High-resolution discrimination between Gram-negative *E. coli* and *S. aureus.* PC1 captures 99% of the variance, illustrating the extreme precision of the transport-enhanced kinetic signal in resolving fundamental differences in bacterial physiology.

Pre-analytical urine centrifugation drastically truncated the initial signal lag phase during real-time fluorometric monitoring. Concentrating the local uropathogenic load at the hydrogel interface enabled rapid, robust metabolic conversion within the critical initial 120 minutes of incubation, whereas raw, uncentrifuged controls exhibited a protracted signal delay (Supplementary Fig. S7). This enhances kinetic trajectory yields an acceleration in early signal-to-noise dynamics, significantly shortening the platform’s practical sample-to-answer timelines. Assay kinetics were similarly modulated by the initial sample inoculation volume under low pathogen loads (10^3^ CFU/mL). Increasing the input volume from 20 µL to 100 µL yielded an accelerated sigmoidal trajectory, substantially minimizing the TTD (Supplementary Fig. S8). Real-time monitoring across the initial 120 minutes highlighted that while the 20 µL inoculation volume remained suppressed at the baseline, the 100 µL volume achieved rapid signal onset within 60 minutes. This kinetic advantage stems from the increased absolute bacterial count delivered to the hydrogel sensor core, indicating that adjusting sample volume can effectively circumvent early metabolic latency without requiring pre-incubation steps. Furthermore, the hydrogel platform maintained diagnostic specificity when evaluated against potential metabolic signal cross-reactivity from background mammalian cellular debris. Inoculation with high-density epithelial cells (A549 line), acting as a surrogate model for exfoliated urinary tract shedding, yielded a completely flat, negligible fluorescence profile over the entire diagnostic timeline like the negative control baseline. This selective metabolic restriction confirms that background mammalian cellular components do not compromise assay fidelity or generate false-positive readouts (Supplementary Fig. S9).

To determine whether antibiotic incorporation affects molecular transport and release within the hydrogel, we next utilized a 12-well plate setup to characterize ampicillin release under both static and orbital conditions, (Fig. 4D). Release profiles were indistinguishable between conditions, indicating that external mixing does not significantly influence diffusion from the hydrogel network. Rapid release was observed, with approximately 60% of loaded antibiotic released within 5 minutes and complete release was achieved within 20 minutes, confirming fast and predictable drug availability within the sensing environment. Building on this, we integrated antibiotics into the hydrogel in the same multi-well configurations to enable direct antimicrobial susceptibility testing (Fig. 4E). In this configuration, nutrients, resazurin, and antibiotics are co-encapsulated, allowing bacterial viability to be directly assessed under antibiotic pressure through the same metabolic readout. In the case of susceptible bacteria, antibiotic activity suppresses metabolic conversion of resazurin, resulting in no observable signal, whereas resistant bacteria maintain metabolic activity and produce a clear colour and fluorescence response. Functional validation of this approach was performed using *E. coli* exposed to different antibiotics. In ampicillin-load hydrogels (Fig. 4F), no detectable signal was observed, consistent with effective inhibition of bacterial growth and metabolism under antibiotic pressure. In contrast, gentamicin-loaded hydrogels (Fig. 4G) also suppressed bacterial detection, although a low-level background signal was observed, suggesting minor assay interference likely arising from antibiotic-reporter interactions rather than bacterial activity. A similar suppression of metabolic signal was observed with meropenem (Fig. 4H), confirming robust inhibition of bacterial growth across multiple antibiotic classes within the hydrogel matrix.

Together, these results demonstrate that the hydrogel platform enables sensitive bacterial detection and direct phenotypic AST within a unified sensing microenvironment. Importantly, antibiotic release occurs rapidly and independently of external transport conditions, ensuring that observed diagnostic readouts reflect true bacterial viability responses rather than limitations in drug availability or diffusion.

### Post-detection bacterial recovery and downstream analysis

To assess whether the hydrogel platform supports downstream microbiological analysis beyond bacterial detection and AST, we investigated the recovery of viable bacteria following completion of the sensing assay. As shown schematically in Fig. 5C, post-detection hydrogels were simply transferred into 1x PBS (5-10 mL) and subjected to orbital agitation (110 rpm) to facilitate bacterial release. This process exploits the inherent swelling behaviour of the hydrogel, combined with diffusion and capillary-driven transport, enabling efficient retrieval of bacteria from the matrix without enzymatic or chemical degradation steps.

We first evaluated recovery efficiency using *E. coli* spiked artificial urine samples across pH conditions (pH 4, 5.5, and 8). Following hydrogel-based detection, bacteria were successfully retrieved and plated on MacConkey agar for confirmation (Fig. 5B). Across all tested pH conditions, characteristic lactose-fermenting *E. coli* colonies were observed, confirming that the hydrogel microenvironment and detection process do not compromise bacterial viability or culturability. We next assessed recovery in mixed-species samples containing both *E. coli* and *S. aureus* at 10^3^ CFU/mL (Fig. 5D, 5E). After retrieval, samples were plated on CLED agar to enable differential colony morphology-based identification. Distinct colony phenotypes were observed, where the lactose-fermenting *S. aureus* appeared as smaller dark brown colonies and *E. coli* presented as yellow/white colonies due to its acidic fermentation of the medium. This demonstrates that the platform preserves species-level differentiation after recovery and supports downstream culture-based analysis.

To further confirm compatibility with molecular diagnostics, we performed colony PCR on bacteria retrieved from NaAMPS hydrogels. Following orbital agitation and release, a standard silica-column DNA extraction protocol yielded dense, distinctly visible white nucleic acid precipitates prior to the final ethanol washing steps (Supplementary Fig. S10, Table S4). This visual presence of these robust pellets confirms that the non-destructive hydrogel elution process yields high concentrations of clean, intact template DNA. Genotypic identification of *S. aureus* was confirmed by amplification of esxA (100 bp amplicon), validating species-specific detection from recovered colonies (Fig. 5F). Similarly, PCR amplification of the 16S rRNA gene (1.4 kb) confirmed the identity of recovered E. coli populations (Fig. 5G). In both cases, clear and specific amplification products were obtained, demonstrating that bacteria recovered from the hydrogel remain suitable for standard molecular workflows.

Therefore, these results demonstrate that the multifunctional NaAMPS sensing platform is fully compatible with downstream phenotypic culture and genotypic analyses. Importantly, bacterial recovery is achieved through a simple resuspension step without specialised reagents or equipment, enabling seamless integration of rapid phenotypic detection with conventional microbiological and molecular diagnostics.

### Hydrogel platform enables rapid and accurate clinical phenotypic diagnostics

To evaluate the clinical utility of the transport-enhanced hydrogel, we transitioned the technology into a high-throughput 96-well format and validated its performance against gold-standard urine culture method. This platform demonstrates a robust capacity for rapid UTI detection and AST by overcoming the diffusion-limited bottlenecks typical of clinical diagnostics.

In a cohort of N = 48 clinical patient samples, the NaAMPS hydrogel platform achieved a diagnostic accuracy of 95.8%. The system identified infections with 95.2% sensitivity (40/42) and 100% specificity (6/6), yielding a Positive Predictive Values (PPV) of 100%. Statistical analysis confirmed strong agreement with conventional urine culture method, evidenced by a Cohen’s Kappa (k) of 0.958. We further assessed the platform’s ability to perform phenotypic AST across nine major antibiotic classes. The hydrogel matrix maintained 100% sensitivity for all tested conditions, correctly identifying every resistant isolate in the N = 25 subset. Specificity remained equally high, reaching 100% for seven of the nine drugs, including Ertapenem, Piperacillin/Tazobactam, and Gentamicin, and only dropping slightly to 94-95% for Amikacin and Meropenem (Supplementary Figs. S11 to S43, Table S5). By embedding convective transport directly into the sensing environment, the platform yields these gold standard matching results at a fraction of the time required for traditional workflows, reducing the diagnostic timeline from 24-48 hours for standard urine culture and an additional 16-24 hours for broth microdilution down to under 2 hours with the hydrogel matrix.

To further safeguard diagnostic specificity against common microbial confounders found in clinical urine^26,27^, we evaluated potential metabolic signal cross-reactivity from commensal urinary fungal species (Supplementary Figs. S44-S46). Because active fungal proliferation^28,29^ could theoretically reduce the encapsulated redox reporter and compromise bacterial readout, we investigated whether the hydrogel matrix could be modified to selectively isolate bacterial signals. Clinical fungal isolates grown on the hydrogel formed distinct surface mycelial structures accompanied by colorimetric reduction from purple to cream/pink (Supplementary Fig. S44A). However, the co-encapsulation of cycloheximide completely arrested this fungal metabolic pathway, yielding a flat kinetic profile over a 1400-minute window and locking the hydrogel matrix in its unreduced purple state (Supplementary Fig. S44B, S45A). Similarly, incorporating the clinical antifungal isavuconazole induced a high-resolution, dose-dependent kinetic suppression, where progressive concentration increases from 200 µg/mL to 250 µg/mL delayed metabolic conversion, culminating in total fungal inhibition 300 µg/mL (Supplementary Fig. S46). This modular integration of antifungals provides a reliable method to suppress fungal metabolic backgrounds, ensuring that the platform’s rapid diagnostic readouts can be dedicated exclusively to the definitive screening of bacterial uropathogens.

The preservation of metabolic kinetic data within the transport-enhanced hydrogel microenvironment allows for the high-resolution discrimination of bacterial species. Using Principal Component Analysis (PCA) to map the interaction between bacteria and the redox reporter, we observed distinct metabolic “fingerprints” for major uropathogens. Clinical isolates of *Klebsiella*, *Proteus*, and *E. coli* occupy clearly defined clusters in the PCA space, with PC1 and PC2 accounting for 71% and 16.4% of the variance, respectively indicating a clear species level separation. Higher-order analysis successfully resolved individual strains of *Pseudomonas*, *Klebsiella*, and *Proteus*, as well as different categories of *E. coli* (including ESBL and ampicillin-resistant strains), demonstrating that the hydrogel matrix amplifies subtle phenotypic differences. The platform achieved nearly absolute separation between Gram-negative *E. coli* and Gram-positive *S. aureus*, where PC1 alone captured 99% of the variance.

To establish a rigorous mathematic foundation for this phenotypic discrimination, the real-time fluorometric kinetic profiles were evaluated using non-linear mixed-effects modelling (NLME). By reparametrizing classical sigmoidal equations to extract explicit biological identifiers – specifically asymptotic signal (A), maximum growth rate (µ_m_), and lag phase duration (λ) – the data were subjected to comparative fitting criteria. Iterative testing across 18 specific model variations altering random-effect structures across Logistic, Gompertz, and Baranyi-Roberts formulations demonstrated that incorporating random effects on maximum growth rate and lag time yielded the optimal statistical fit (Supplementary Table S6). Across all datasets, the Gompertz growth model demonstrated superior descriptive fidelity over Logistic and Baranyi-Roberts formulations (Supplementary Table S7), consistently yielding the lowest Akaike Information Criterion (AIC) scores and minimum root-mean-square error (RMSE). This mathematical framework cleanly captured raw sigmoidal growth kinetics mapped both collectively (Supplementary Fig. S47) and across individual species panels (Supplementary Fig. S48). Furthermore, a deep-dive correlation matrix analysis of initial bacterial loads (10^1^ to 10^7^ CFU/mL) (Supplementary Fig. S49) revealed a near-perfect negative correlation between concentration and lag time (λ = -0.991, p < 0.001). Strikingly, the maximum specific growth rate (µ_m_) remained entirely independent of the initial bacterial concentration (p > 0.05). This mathematically demonstrates that while the initial pathogen load dictates when the sensor registers an exponential shift in log population size (fluorescence increase) ln (N_t_/N_0_), the slope of the kinetic trajectory (µ_m_) acts as a highly conserved, concentration-invariant, species-specific fingerprint that validates high PCA sorting accuracies.

To establish the operational robustness, assay generalizability, and inter-user portability of this metabolic profiling framework, cross-operator validation experiments were conducted in artificial urine. Standard concentration-dependent dose curves spanning a broad, nine-log linear dynamic range (10^1^ to 10^9^ CFU/mL) were independently reconstructed across multiple clinical uropathogenic species utilizing both real-time fluorometric kinetic monitoring and colorimetric verification workflows. Crucially, the independently generated sigmoidal profiles and derived calibration curves demonstrated exceptional fidelity to the primary data (Supplementary Fig S50). This confirms the quantitative analytical performance and detection sensitivity are fully preserved irrespective of operator-dependent variation, highlighting the high translational viability and ease-of-use of the platform for decentralized clinical settings. Importantly, the non-destructive nature of the hydrogel core allowed for the post-detection recovery of bacteria directly from positive patient samples. Following raw clinical screening, captured bacterial uropathogens were successfully retrieved from the 96-well hydrogel matrix via saline agitation and re-cultured on chromogenic agar, demonstrating that the diagnostic microenvironment preserves cellular culturability and viability during clinical workflows (Supplementary Figs. S51-S52). These results establish that the swelling-driven convective transport not only accelerates detection time by 12-48-fold but also provides a high-fidelity diagnostic signal. This capability enables a single-step workflow for simultaneous pathogen detection, identification, and susceptibility profiling suitable for both laboratory instrumentation and point-of-care deployment.

## Discussion

While phenotypic testing remains irreplaceable for capturing non-genetic and hetero-resistant traits that genome sequencing alone cannot predict^30–32^, its operation is often quietly constrained by the physics of mass transport rather than the intrinsic limits of cellular metabolism. In traditional diagnostic formats, passive molecular diffusion dictates the rate of bacteria-reporter interactions, inadvertently stretching clinical timelines to hours or days. Here, we demonstrate that this transport limitation is an engineering hurdle rather than an intrinsic constraint of phenotypic testing. By delegating active fluidic handling to the intrinsic thermodynamics of responsive ionic hydrogel, we introduce material-driven transport as a core design principle for biosensors, shifting transport control from complex external hardware directly to the autonomous sensing matrix itself. We leverage this self-contained transport mechanism to rapidly isolate and capture metabolic signals directly from clinical urinary tract infections (UTIs), compressing diagnostic timelines from days to minutes.

The 12-48-fold acceleration in bacterial detection times marks a critical transition away from classical diffusion-limited kinetics. In standard solution-phase or static matrix assays, the localised consumption of reporter molecules around bacteria rapidly creates a depleted boundary layer, yielding a typical √*t* signal plateau that delays diagnostic timescales. The dynamic NaAMPS matrix fundamentally bypasses this bottleneck, driving an apparent transport rate that outpaces the solution phase-diffusion by 4.45-fold. This transport enhancement aligns with the high density of fixed negative sulfonate groups within the NaAMPS network, which generates localized osmotic driving forces and capillary flow during swelling. This matrix-mediated convective transport model is strongly supported by the system’s macro-scale geometric dependence, providing a physical mechanism for the continuous redistribution of reporter molecules toward metabolically active cells, mitigating local substrate exhaustion, and sustaining high-frequency interactions.

Optimizing this materials-driven sensing environment relies on balancing the hydrogel’s physical structure with its chemical composition. Volumetric swelling, while vital, does not act in isolation; as demonstrated by the neutral PAAm network, the lack of fixed ionic charges prevents the generation of the electrostatic pressure differentials that assist advective transport. At the same time, the network’s structural crosslinking density plays a defining role in shaping the assay. While a lower crosslinking density can compromise macroscopic mechanical stability, a highly crosslinked network introduces physical confinement that restricts natural bacterial growth patterns into crowded, confluent configurations. By balancing these structural constraints (0.2 mM PEGDA) and integrating a divalent ion-rich nutrient media (M2), the system compresses the bacterial time-to-detection to 75 minutes. This demonstrates how the physical architecture and chemical microenvironment of the hydrogel can be tuned in tandem to enhance phenotypic signal transduction.

When configured for high-throughput clinical evaluation, this platform handles the heterogeneous nature of patient samples with exceptional robustness. The native buffering capacity of the pre-encapsulated medium effectively normalizes extreme physiological fluctuations in urine pH (4.5-8.0), while the integration of targeted antimycotics (cycloheximide and isavuconazole) completely isolates the bacterial metabolic signal from potential fungal backgrounds. Crucially, because antibiotic release occurs rapidly and independently of external mixing, the resulting susceptibility profiles reflect true phenotypic cellular responses rather than drug diffusion artifacts. Furthermore, the highly hydrated, non-destructive architecture of the matrix permits the effortless, reagent-free elution of viable, intact pathogens, serving as a universal bridge connecting rapid, point-of-care screening directly to downstream molecular workflows.

While the platform demonstrates exceptional clinical accuracy (95.8%), sensitivity (95.2%) and specificity (100%) across a 48-patient cohort, certain molecular boundaries warrant further exploration. The low-level background signal observed in gentamicin-loaded matrices suggests minor chemical cross-reactivity between specific aminoglycoside structures and the redox reporter, a phenomenon requiring systematic molecular mapping to refine baseline thresholds. Beyond UTIs, the fundamental nature of this matrix-mediated transport mechanism allows it to be readily adapted for the rapid detection and susceptibility profiling of other localised or systemic infections. Ultimately, this technology demonstrates that intrinsic material properties can bypass the need for complex, external fluidic infrastructure to solve transport limitations. By leveraging properties of NaAMPS hydrogels to overcome the mass transport bottleneck, this approach removes the dependency on active mixing or peripheral pumping hardware, providing a highly scalable architecture that seamlessly integrates rapid phenotypic testing into standard clinical plate-reader workflows.

## Materials

### NaAMPS synthesis

NaAMPS hydrogels were fabricated via free-radical cross-linking polymerization. The precursor formulation consisted of 2-acrylamido-2-methylpropane sulfonic acid sodium salt (NaAMPS) monomer (50 wt% in H_2_O, 0.10 mol;Sigma-Aldrich), poly(ethylene glycol) diacrylate cross-linker (PEGDA, Molecular weight = 575 g/mol, 0.2 mmol; Sigma-Aldrich), and 2-hydroxy-2-methylpropiophenone (Irgacure 1173/HMPP, Sigma-Aldrich) as the water-soluble photo-initiator. To initiate polymerization, the homogenous precursor solution was exposed to ultraviolet (UV) light. Photolysis of the Irgacure 1173 initiator generates reactive free radicals, which attack the terminal alkene double bonds (C=C) of both the PEGDA cross-linker and the NaAMPS monomers. This radical addition propagates a continuous chain growth polymerization and crosslinking network, extending the reaction until a stable, covalently interconnected three-dimensional NaAMPS hydrogel matrix is formed. 2mL of precursor solution was transferred into round bottom silicone moulds and placed in belt conveyor UV curing system (GEW engineering UV with E2C lamp, power of 140W/cm). It was passed through UV conveyer system at least 6 cycles of 1 minute to ensure complete polymerisation.

### Swelling kinetics

Hydrogel swelling analysis was performed first in water for a total duration of 360 minutes (6 hours). For this duration, the weight of hydrogels was measured at 30-minute intervals. % swelling capacity was calculated using the formula:

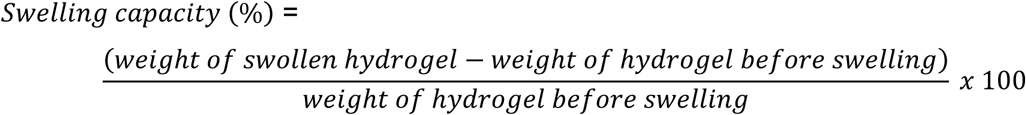

Similarly, the swelling behaviour was determined in 3 pH conditions. LB media (pH = 7) was adjusted to pH = 4 using 1M hydrochloric acid (HCl) and pH = 10 using 1M sodium hydroxide (NaOH). The weight of hydrogels was determined every 30 minutes over a total duration of 150 minutes.

### Resazurin preparation

Resazurin stock solution was prepared by dissolving 0.5 grams of resazurin powder in 100 mL sterile 1X PBS. For NaAMPS hydrogels, 100 µL of resazurin stock solution was introduced in 20 mL media. NaAMPS hydrogels were then introduced in the mixture for their uptake by the hydrogel.

### NaAMPS functionalisation

Post-synthesis, the hydrogels were introduced in 20mL of Mueller-Hinton Broth – 2 (M2) bacterial media containing 5mg/mL resazurin. The NaAMPS hydrogels were incubated in this solution for 12 hours. The resulting NaAMPS hydrogels are purple in colour. For antibiotics: Ampicillin 10 µg/mL was used as the testing concentration for AST against *E. coli* according the European Committee on Antimicrobial Susceptibility Testing (EUCAST). A stock solution of 100 mg/mL ampicillin solution was obtained from the media preparation facility at the Warwick Medical School. 24 µL from stock solution was added into a solution containing 20 mL M2 media and 1µg/mL resazurin. The solution was mixed thoroughly and transferred into a 250mL glass beaker. NaAMPS hydrogels were introduced in the beaker for uptake of the solution for 10 hours at room temperature. As required 15mm, 8mm, and 3mm hydrogels were punched from the swollen NaAMPS hydrogels using a metallic punch. The final resazurin concentration with the gel is 1 µg/mL. The final concentration of ampicillin in 10 µg/mL. The punched-out hydrogels were then introduced in multi-well plates. Gentamicin sulfate (code: 48760) and Meropenem (code: PHR1772) purchased from Sigma Aldrich, Merck were obtained. Similarly, 72 µL gentamicin from 10 mg/mL stock gentamicin solution was added into the solution so the final concentration is 30 µg/mL (EUCAST recommended). 48 µL from 50 mg/mL stock meropenem solution was added into the solution so that the final concentration is 10 µg/mL (EUCAST recommended). For clinical testing, the antibiotics were encapsulated within the hydrogels at the minimum inhibitory concentrations/clinical breakpoints as provided by the EUCAST guidelines. The breakpoints used for determining AST were: 30 µg cephalexin, 10 µg cefpodoxime, 30 µg piperacillin + 6 µg tazobactam, and 10 µg gentamicin, 30 µg amikacin, 20 µg amoxicillin, 10 µg meropenem.

### Preparation of artificial urine

Artificial urine was formulated by systematically dissolving the following chemical components in 1L of sterile, autoclaved Milli-Q water under constant magnetic stirring at room temperature: 0.651 g of calcium chloride dihydrate (CaCl_2_.2H_2_; SRL, Cat. No.70650), 0.651 g of magnesium chloride hexahydrate (MgCl_2_.6H_2_O;SRL, Cat. No. 91417), 4.6 g of sodium chloride (NaCl; Sigma-Merck, Cat. No. S9888), 2.3 g of sodium sulfate (Na_2_SO_4_; Sigma-Merck, Cat. No. 204447), 0.65 g of trisodium citrate (Na_3_C_6_H_5_O_7_; Sigma-Merck, Cat. No. S4641), 0.023 g of sodium oxalate (Na_2_C_2_O_4_; Sigma-Merck, Cat. No. 62760), 2.8 g of monopotassium phosphate (KH_2_PO_4_; Sigma-Merck, CAS. No. 7778-77-0), 1.6 g of potassium chloride (KCl; SRL, Cat. No. 38630), 1.0 g of ammonium chloride (NH_4_Cl; SRL, Cat. No. 25103), 25.0 g of urea (CO(NH_2_)_2_; Sigma-Merck, Cat. No. 57136), and 1.1 g of creatine (C_4_H_9_N_3_O_2_); Sigma-Merck, Cat. No. 6020877). Agitation was maintained until all solid compounds were completely solubilized, and the resulting matrix was used immediately or stored under sterile conditions for downstream validation.

### Antibiotic release

Ampicillin release kinetics was determined using ultraviolet (UV) -visible spectroscopy in FLUOstar OMEGA plate reader (BMG Labtech, UK). The hydrogels were suspended in 1X phosphate buffered saline (PBS) pH = 7 in glass beakers. From this solution, 200 µL was used for spectroscopy measurements and 200 µL of fresh 1X PBS was replaced in the antibiotic suspended PBS solution. The UV absorbance was measured every 5 minutes at 260 nm and the % antibiotic release was calculated using the formula:

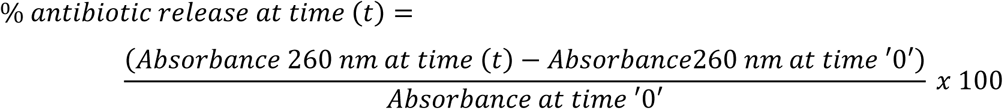

The effect of antibiotic release from hydrogels was also evaluated in orbital shaking and static conditions. In orbital shaking, the hydrogels suspended in 1X PBS in glass beakers were placed in orbital shaker at 110 RPM.

### Optimization of resazurin concentration

Photopolymerized NaAMPS hydrogels were introduced in 5 different solutions, each containing 20 mL M2 media, each with varying resazurin concentrations – 1 µg/mL, 2 µg/mL, 3 µg/mL, 4 µg/mL, and 5 µg/mL. NaAMPS hydrogels were swollen for 10 hours overnight. 1.5cm diameter hydrogels were punched-out using a sterile metallic punch and the hydrogels were introduced in 24-well plate for identifying the optimal resazurin concentration in one-experimental set-up. For each of the resazurin concentration condition, the experiments were performed in three replicates. Therefore, the experimental set-up consisted of 21 total number of hydrogels: 3x blank hydrogels, 3x positive control hydrogels, 3x (1 µg/mL, 2 µg/mL, 3 µg/mL, 4 µg/mL, and 5 µg/mL resazurin concentration) hydrogels. 20 µL of bacterial culture containing 10^7^ CFU/mL uropathogenic *E. coli* is introduced in the positive control. For hydrogels with different resazurin concentrations, 20 µL of artificial urine containing 10^6^ CFU/mL uropathogenic *E. coli* in artificial urine is introduced. The plate was then placed in the fluorescent plate reader (FLUOstar Omega BMG LABTECH). The fluorescence measurements were recorded at 560 nm excitation wavelength and 610 nm emission wavelength every 15 minutes for 18 hours. The resorufin fluorescence outputs were analysed in OriginPro 2021b. The end point qualitative images were captured in a smartphone camera. To identify the optimal resazurin concentration for encapsulation within NaAMPS hydrogels, we determined the maximum bacterial growth rate from the resorufin outputs. This was determined by using the Gompertz non-linear curve fitting in OriginPro 2021b. The resorufin outputs are arranged with x-axis = ‘time’ and y-axis = ‘fluorescence intensity’ for a time interval of 15 minutes over a period of 18 hours. The model fitting is performed using growth/sigmoidal non-linear curve fit tool with SGompertz function and Levenberg Marquardt iterative algorithm. The resulting bacterial growth rate (µ) is identified and compared across different multifunctional NaAMPS hydrogels with varying resazurin concentrations. The coefficient of determination R^2^-square value is noted, and the best fit is determined with R^2^-square values corresponding to ∼ 0.995.

### GFP-tagged *E. coli* culture

*E. coli* S17.1 (pUB7 λpir with pBAM7 plasmid expressing green fluorescent protein (GFP) and kanamycin resistance gene) was cultured aerobically in lysogeny broth (LB) at 37°C at 180 rpm overnight. The overnight cultures were further sub-cultured into fresh LB and grown to an optical density (OD)_600 nm_ = 1 and further diluted to (OD)_600 nm =_ 0.5. The culture was 10-fold serially diluted to obtain a colony count ∼1000 CFU/mL. 20 µL of this bacterial culture was plated onto LB agar control plates and NaAMPS hydrogels.

### LOD and bacterial concentrations

LOD was first identified by measuring the fluorescence of blank multifunctional NaAMPS hydrogels (hydrogels in which bacterial culture/artificial urine containing bacteria are not introduced). The fluorescence measurements are recorded every 5 minutes for 2 hours. The standard deviation of the blank fluorescence measurements was calculated, and the LOD value was determined by 3x standard deviation of the blank. A primary culture of uropathogenic *E. coli* is prepared by inoculating the culture from glycerol stock in LB media followed by overnight incubation at 37°C and 180 RPM under orbital shaking. The OD_600nm_ of the culture is measured and adjusted to 0.5 by diluting with LB media. 1:10 dilutions of uropathogenic *E. coli* with starting concentration of 10^7^ CFU/mL down to 10 CFU/mL is prepared. 1.5 cm multifunctional NaAMPS hydrogels encapsulated with 1µg/mL resazurin and M2 media were deposited in 24-well multi-well plate. 20 µL of each bacterial concentrations were added separately in hydrogels. All experiments were performed in three replicates.

### Optimization of encapsulated bacterial culture media

To determine the optimal media for encapsulation within NaAMPS hydrogels for bacterial detection, 5 bacterial culture media including NB (code: CM0001 from Oxoid, ThermoFisher Scientific), LB (code: L3522, Sigma Aldrich, Merck), TSB (code: 1.00525 Millipore), M1 (code: 70192, Millipore), and M2 (code: 90922, Millipore) are separately encapsulated within NaAMPS hydrogels via swelling. A sterile 24-well plate is used to accommodate 1.5 cm diameter multifunctional NaAMPS hydrogels, each with NB, LB, TSB, M1, and M2 in three replicates. Artificial urine containing10^6^ CFU/mL Uropathogenic *E. coli* is then introduced on the five types of bacterial culture media encapsulated hydrogels. 10^7^ CFU/mL Uropathogenic *E. coli* culture is introduced in multifunctional NaAMPS hydrogels which consisted of resazurin and M2 bacterial culture media. These hydrogels act as positive controls. The assay included blank multifunctional NaAMPS hydrogels which consisted of resazurin and M2 bacterial culture media to which bacteria/urine is not added. The fluorescence measurements were recorded as described earlier.

### Interference from epithelial cells

A549 type II human alveolar epithelial cells obtained from the American Type Culture Collection (ATCC) are cultured in T75 flasks (Falcon) in Dulbecco’s Modified Eagle Medium (DMEM) supplemented with 10% Fetal Bovine Serum (FBS) and 1% penicillin + streptomycin at 37°C and 5% CO2. When the epithelial cells are 80% confluent, they are trypsinised and introduced into new DMEM medium. The freshly transferred epithelial cells are diluted to a concentration of 5×104 cells with sterile 1X PBS and centrifuged at 3000 RPM for 3 minutes. The cell pellet is again washed with sterile 1X PBS and centrifuged at 3000 RPM for 3 minutes to completely remove trypsin/DMEM if any. The pellet is resuspended with 20mL artificial urine. 20 µL of artificial urine is introduced in multifunctional NaAMPS hydrogels encapsulated with 0.5X resazurin and M2 media, hydrogels pre-deposited in 12-well multi-well plate. The final concentration of alveolar epithelial cells introduced in the hydrogel is 50 cells (per 20 µL). The fluorescence measurements were recorded as described earlier.

### Bacterial retrieval and downstream testing

For bacterial retrieval, post-bacterial detection, the multifunctional NaAMPS hydrogels were transferred into a sterile 20 mL tube containing 1X PBS. The tubes were vortexed under shaking conditions 110 RPM at room temperature for 10 minutes to facilitate bacterial detachment from the hydrogels into the PBS solution. 100 µL of the retrieved solution was plated on MacConkey agar and incubated at 37°C for 18 hours. 10 µL of this solution was used for direct PCR. After retrieval, colonies were picked from MacConkey agar plates and resuspended in 50 µL sterile water. The mixture was heated at 98°C for 10 minutes and 10 µL was used for PCR .PCR was performed to amplify the 16S rRNA genes using Q5® High-Fidelity DNA Polymerase (New England BioLabs). 16 S rRNA primers were synthesized by Integrated DNA Technologies. The primer sequences are shown in table 8. PCR cycle conditions were set-up as follows: 98°C initial denaturation for 30 seconds, 98°C denaturation for 10 seconds, 65°C primer annealing for 30 seconds, 72°C extension for 30 seconds/kb, 72°C final extension for 2 minutes. Amplification reactions were performed in a total volume of 25 µL per well. Each reaction mixture contained 2.5 µL of 5X Q5 Reaction Buffer (1X final concentration), 0.5 µL of 10 µM dNTPs (200 µM final concentration), 1 µL each of 10 µM forward and reverse primers (0.5 µM final concentration each), 10 µL of template DNA, and 0.25 µL of Q5 High-Fidelity DNA Polymerase (0.02U). The remaining volume was brought to 25 µL using 9.75 µL of nuclease-free water. Primers used: esxA forward primer (5’-GCGGCTAGCATGGCAATGATTAAGATGAG-3’), esxA reverseprimer(5’-ATCGCGGCCGCCTAGCTGCTCCAGTGGCTCACGG CCCTCACCCTGTCGGGTTGCAAACCGAAATTATTA G -3’, 16S rRNA forward primer (5’ – AGAGTTTGGATCMTGGCTCAG -3’), and 16S rRNA reverse primer (5’ - CGGTTACCTTGTTACGACTT - 3’).

### DNA extraction

DNA extraction was performed using the DNeasy Blood & Tissue kit by following the specified protocol for bacterial DNA extraction. After bacterial retrieval, the bacteria were pelleted by centrifugation for 5 minutes at 8000 RPM. The pellets were resuspended in 200 µL PBS and 20 µL proteinase K was added. 2.5 µL RNase A (100 mg/mL) was added and mixed by vortexing. The solution was incubated for 2 minutes at room temperature. 200 µL Buffer AL was added and thoroughly vortexed. The solution was then incubated at 56°C for 10 minutes. Next, 200 µL ethanol was added to the mixture and vortexed to obtain a homogenous solution. The mixture was then pipetted into the DNeasy Mini spin column with a 2mL collection tube. It was centrifuged at 8000 RPM for 1 minute. The flow-through was discarded. The column was washed with 500 µL of Buffer AW1 and spun at 8000 RPM. Next, 500 µL of Buffer AW2 was added and spun at 14000 RPM for 3 minutes. Finally, 200 µL buffer AE was added to the column and incubated for 1 minute. The column was the centrifuged at 8000 RPM for 1 minute for DNA elution.

### Principal Component Analysis (PCA)

PCA was performed using OriginPro 2021b software. The input dataset consisted of hydrogel outputs which were categorized as x-axis = time (2 hours) and y-axis = resorufin fluorescence intensities for 2-hour duration. An enhanced version of PCA called PCA v1.50 application was used. The generated outputs included eigen values, scores, and loadings. The score plot and loadings plot were utilised to extract the principal components in two dimensions (PC1 and PC2). The components were plotted using the scatter plot. Furthermore, to determine the confidence interval for the clustered bacterial species, a two-dimensional confidence ellipse at 95% was superimposed on the PCA plot using the 2D confidence ellipse v1.40 application in originPro.

### Encapsulation of antifungal agents

50 µg/mL cycloheximide in water was encapsulated within NaAMPS hydrogels in addition to media and resazurin as described earlier. 100 µg/mL, 200 µg/mL, 250 µg/mL, and 300 µg/mL isavuconazole in DMSO was encapsulated within NaAMPS hydrogels in addition to media and resazurin as described before.

### Blind cohort study design

Clinical urine specimens (N=48) were obtained from the Arden Tissue Bank at University Hospitals Coventry and Warwickshire (UHCW) NHS Trust. Samples were collected and utilized in accordance with the ethical framework of the Arden Tissue Bank under REC reference 18/SC/0180. To ensure rigorous, unbiased evaluation, the validation was executed as a double-blind diagnostic study. We performed the hydrogel-mediated phenotypic screening, species identification, and AST profiling entirely blind to the patients’ clinical histories, symptomatic profiles, and existing microbiological diagnostic records. The independent gold-standard clinical urine culture and broth microdilution data generated by the hospital were withheld until we completed the entire hydrogel assay cohort. Once our testing was complete, we cross-checked our outputs against the UHCW records to calculate diagnostic accuracy.

Clinical urine specimens were inoculated directly onto the 96-well sensing platforms and incubated at 37°C inside a multi-mode microplate reader (BioTek Cytation 5 Cell Imaging Multimode Reader (Agilent). Real-time quantitative monitoring was captured simultaneously across two primary analytical channels over a timeline spanning 15 minutes to 2 hours: fluorometric kinetic tracking (excitation/emission mapped via relative fluorescence units) and spectrophotometric absorbance tracking (OD 600nm). Concurrently, end-point visual colorimetric tracking was documented to confirm a binary, instrument-free naked-eye readout (unreduced purple indicating negative infection or antibiotic susceptibility; metabolic conversion to pink/cream indicating active bacterial growth or resistance).

### Calculations

% sensitivity = (True Positives) / (True Positives + False Negatives) * 100 % specificity = (True Negatives) / (True Negatives + False Positives) * 100 The Positive Predictive Value (PPV) and Cohen’s Kappa (*κ*) coefficients were calculated using OriginPro software, where *κ* was determined via a cross-tabulation confusion matrix comparing the binary diagnostic classifications of the hydrogel assay directly against the UHCW gold standard culture results.

### Kinetic growth reparameterization and model fitting

The fluorometric kinetic profiles were evaluated using non-linear mixed-effects (NLME) modelling. Classical sigmoidal growth equations were reparametrized to extract explicit biological identifiers from the time-resolved fluorescence data, isolating the asymptotic signal (A), the maximum specific growth rate (µ_m_), and the lag phase duration (λ). Three foundational mathematical formulations were evaluated and compared:

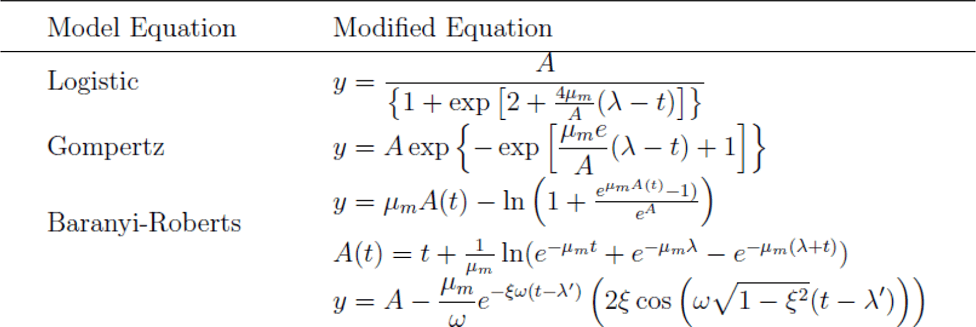

Iterative testing was conducted across 18 distinct structural model variations by systematic allocation of random-effect structures to the primary growth parameters (A, µ_m_, λ).

Model optimization, parameter estimation, and random-effects configurations were executed using R software. The descriptive fidelity and predictive performance of each candidate model evaluated using comparative statistical criteria. The optimal model formulation was selected by minimizing the Akaike Information Criterion (AIC) and the root-mean0square error (RMSE) across all aggregated patient and strain datasets. Following model fitting, the derived biological parameters (µ_m_, λ) for initial bacterial loads spanning a 7-log linear dynamic range (10^1^ to 10^7^ CFU/mL) were extracted for higher-order correlation analysis. Bivariate correlation matrices were constructed in R to map the interdependence between initial pathogen concentration and extracted kinetic parameters. Statistical significance was verified using Pearson Correlation Coefficients (r) and two-tailed Student’s t-tests, evaluating the concentration-dependence of λ against the concentration-invariance of the growth trajectory slope (µ_m_).

## Contributions

J.C. and A.D. conceived the study. J.C., C.C., and A.D. designed the experiments. A.D. performed the majority of the experimental work, conducted principal component analysis and clinical testing. A.D and J.C wrote the manuscript. A.D. synthesized the NaAMPS hydrogels under the guidance of D.M.H. and R.A.H. Y.Y. performed the model fitting and statistical analyses under the guidance of J.B. T.D. performed cross-operator validation experiments under guidance of M.U. and A.D. N.R. designed the blind clinical validation experiments. All authors reviewed and edited the manuscript.

## Acknowledgements

We sincerely thank the Arden Tissue Bank for providing the clinical urine samples used in this study. We are particularly grateful to Hiten Mistry and Mandip Hira at University Coventry & Warwickshire for their invaluable assistance with sample collection, sorting for clinical testing and logistics. We also extend our gratitude to Dr Spyros Efstathiou and Dr Gavin Kirby (University of Warwick) for initial discussions on NaAMPS hydrogel synthesis. We are grateful to Prof. Andrew McAinsh, Warwick Medical School for valuable discussions, encouragement, and continued support. Finally, we thank Warwick Innovations for their guidance and support regarding intellectual property management.

