## Supplementary information for "Embedded transport accelerates interaction-limited biosensing"

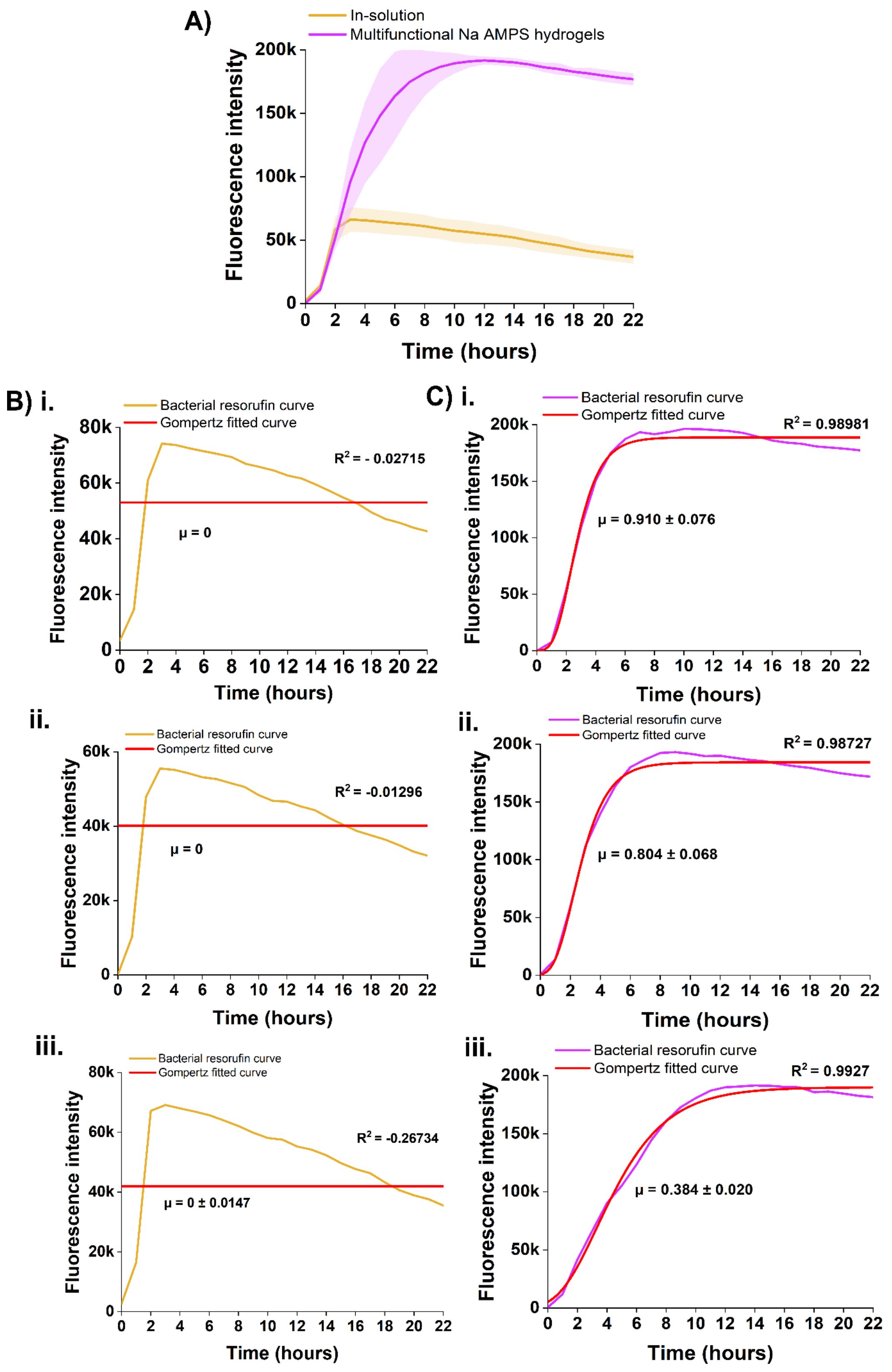


**Figure S1.** Comparison of bacterial resorufin kinetics and Gompertz model fitting between in-solution assays and multifunctional NaAMPS hydrogels. **(A)** Time-dependent fluorescence intensity profiles of bacterial resazurin-to-resorufin conversion over 22 hours, comparing conventional in-solution assays (yellow line) and mulitfunctional NaAMPS hydrogels (purple line). Shaded regions represent the standard deviation of replicates (n=3). **(B) i-iii.** Individual kinetic trajectories of bacterial resorufin production in-solution (yellow curves) across three distinct replicates, fitted against the classical Gompertz growth model (red lines). In all solution-phase replicates, the Gompertz model fails to capture the rapid signal decline following early saturation, resulting in poor goodness-of-fit metrics ($R^2$ = 0) and an estimated specific signal accumulation rate ($\mu$) of zero) (0±0.0147). This behaviour highlights a transport-limited regime and rapid local substrate depletion. **(C) i-iii.** Individual kinetic trajectories of bacterial resorufin production within the multifunctional NaAMPS hydrogel matrix (purple curves) across three distinct replicates, fitted against the Gompertz growth model (red lines). The hydrogel system exhibits a string sigmoidal fit ($R^2$ = 0.98 across all replicates) with robust, quantifiable specific signal accumulation rates ($\mu$). This continuous signal accumulation confirms that the dynamic, swelling-driven matrix mitigates passive diffusion limitations, preventing early local saturation and supporting prolonged bacteria-substrate interactions.

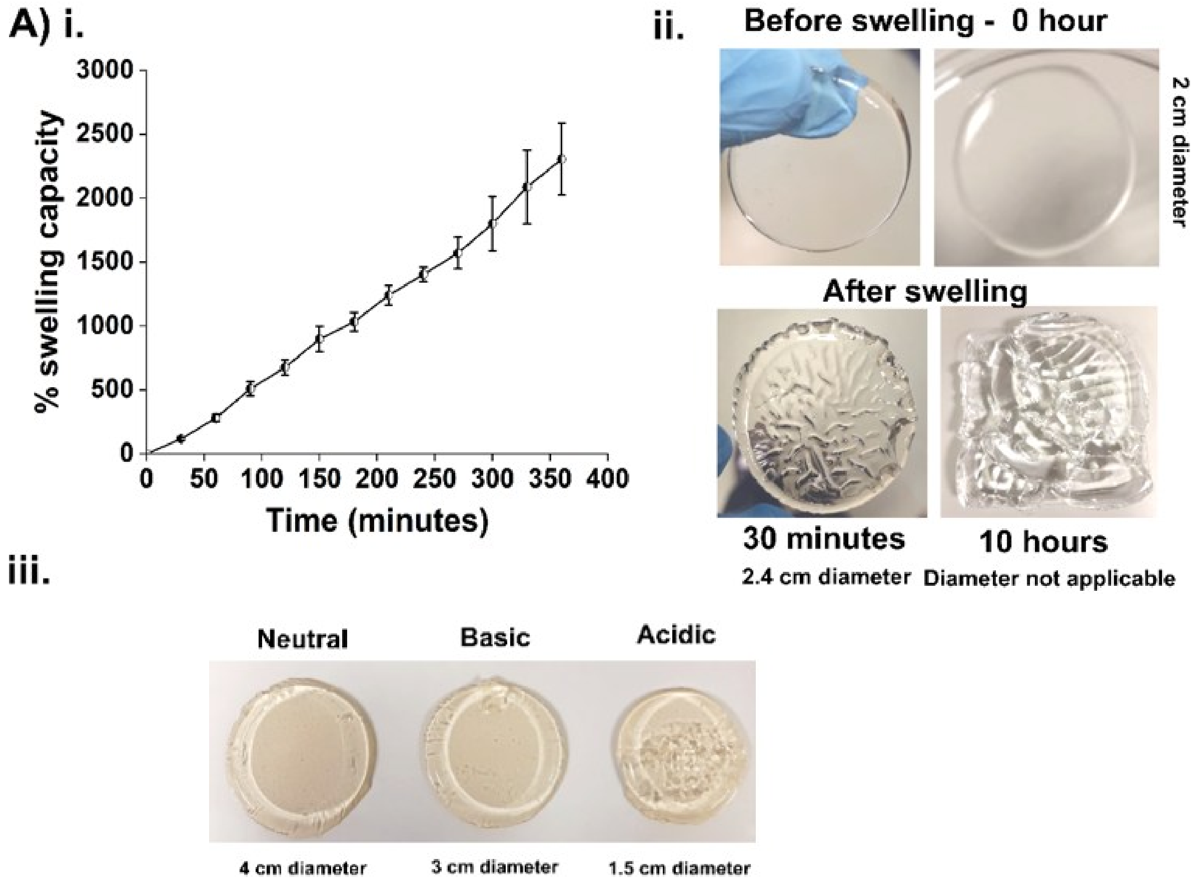


**Figure S2.** Characterization of temporal swelling kinetics and pH-dependent macro-scale expansion of NaAMPS hydrogels. **(A) i.** Quantitative swelling profile of NaAMPS hydrogel over a 360-minute timeline, expressed as percentage of swelling capacity. The network exhibits rapid, sustained hydration, surpassing 2000% volumetric/mass expansion by 6 hours. Data points represent mean ± standard deviation (n=3). **(ii).** Representative macroi-scale photographs illustrating the real-time physical expansion and structural changes of a hydrogel disk. The matrix transitions from a uniform 2 cm diameter disk at 0 hours (top panels) to an expanded 2.4 cm disk at 30 minutes (bottom left). Prolonged incubation (10 hours, bottom right) results in extreme hydration and network expansion where localized gel deforemation renders traditional boundary and diameter measurements non-applicable. **(iii).** Visual macro-scale comparison of NaAMPS hydrogel disk diameters and structural stability across varying environmental pH regimes. Maximum network expansion is achieved under neutral conditions (4 cm diameter). Basic media yields a slightly more constrained expansion (3 cm diameter), whereas acidic conditions severely suppress swelling (1.5 cm diameter) and induce internal network texturing/opacity. This environmental sensitivity directly highlights the role of sulfonate group ($-SO_3^-$) inonization in driving the electrostatic repulsion and osmotic forces required for matrix hydration.

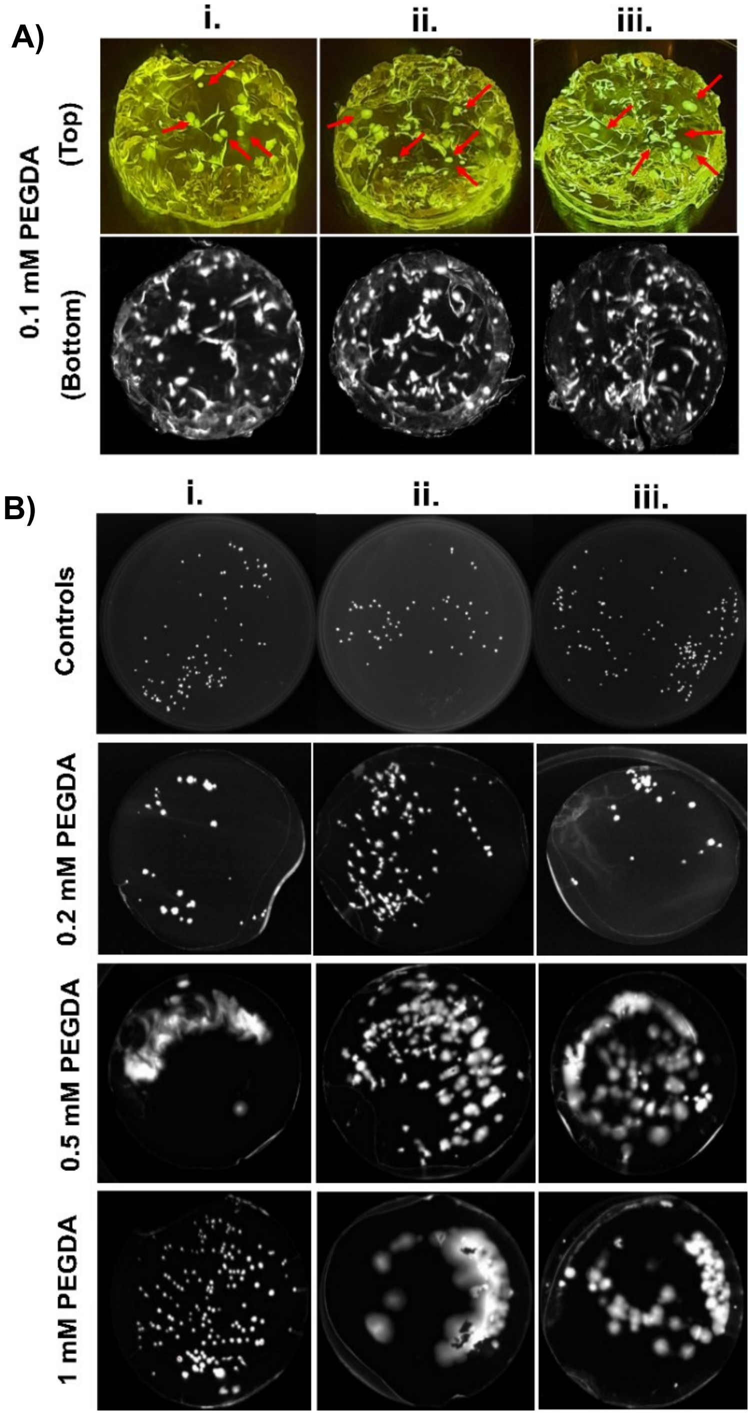


**Figure S3.** Macro-scale distribution and spatial organization of GFP-expressing E. coli colonies within NaAMPS hydrogels at varying crosslinking densities. **(A) i-iii.** Representative dual-view macro-images of three independent replicates of 0.1 mM PEGDA crosslinked hydrogel networks under UV excitation. The top panel (color) highlights physical matrix instability and uneven fragmentation, with distinct macroscopic pocketing. Red arrounds indicate bacterial growth and uneven colony settling across the hydrogel's internal structure due to deficient network structural integrity. **(B) i-iii.** Comprehensive spatial profiling of bacterial colony architecture across three independent replicates (i, ii, iii) as a function of PEGDA crosslinker concentration (0.2mM, 0.5mM, and 1mM) relative to conventional agar control platinge (top row). At 0.2mM PEGDA, the matrix preserves high spatial confinement, yielding sharp, discrete, and isolated fluorescent bacterial colonies. Increasing crosslinking density to 0.5 mM and 1 mM prompts a transition toward restrictive pore size, limiting deep spatial infiltration causing colonies to grow into localized, confluent sheets and merged, diffuse fluorescent zones along the boundaries.

| **Media** | **Uropathogenic *E. coli* growth rate (μ)** | **Lag time ($t_{lag}$)** | **Maximum fluorescence intensity** | **Slope of exponential region** |
|---|---|---|---|---|
| NB | 0.532 ± 0.023 | 2.521 ± 0.098 | 224094 ± 5162 | 39463.1 ± 2083 |
| LB | 0.610 ± 0.058 | 1.991 ± 0.054 | 339677.093 ± 3861 | 54012.395 ± 2174.55 |
| TSB | 0.656 ± 0.020 | 1.011 ± 0.132 | 344919.833 ± 2178 | 56531.852 ± 4960.393 |
| M1 | 0.872 ± 0.057 | 1.003 ± 0.044 | 28195.709 ± 2964 | 60753.616 ± 3895 |
| M2 | 0.915 ± 0.076 | 0.954 ± 0.024 | 262395.932 ± 3316 | 64071.483 ± 3303 |

**Table S1.** Comparative uropathogenic E. coli metabolic tracking parameters across various microbiological growth media encapsulated within NaAMPS hydrogels. Data represent the mean ± standard deviation (n=3). Kinetic traits (μ and $t_{lag}$) were derived via non-linear regression modelling using the classical Gompertz growth equation applied to the real-time fluorometric curves shown in the main text. Abbreviations: NB, Nutrient Broth; LB, Lysogeny Broth; TSB, Tryptic Soy Broth; M1, Mueller Hinton Broth 1 and M2, Mueller Hinton Broth 2 formulation.

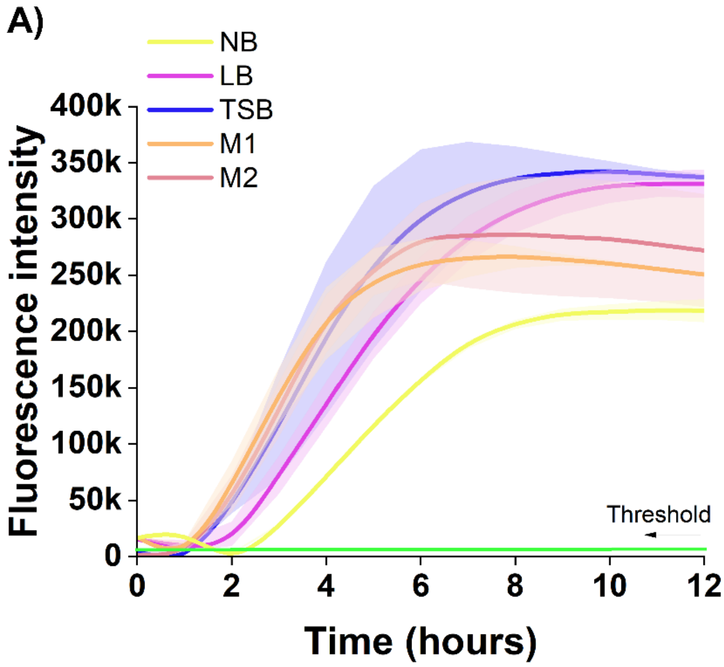


**Figure S4.** Comparative real-time fluorometric kinetic profiles of uropathogenic E. coli metabolic detection across various encapsulated growth media. (A) Time-dependent fluorescence accumulation trajectories monitored over 12 hours within NaAMPS hydrogel matrices formulated with alternative nutrient backgrounds: Nutrient Broth (NB, yellow line), Lysogeny Broth (LB, purple line), Tryptic Soy Broth (TSB, blue line), Mueller Hinton Broth -1 (M1, orange line), and Mueller Hinton Broth -2 (M2, pink line). Shaded regions signify the standard deviation across independent biological replicates (n=3). The baselines green line represents the negative control thershold. The M2 composition demonstrates fastest time-to-detection (TTD) and validating the quantitative kinetic advantages such as minimized lag times ($t_{lag}$) and maximized exponential slopes- tabulated in Supplementary Table S1.

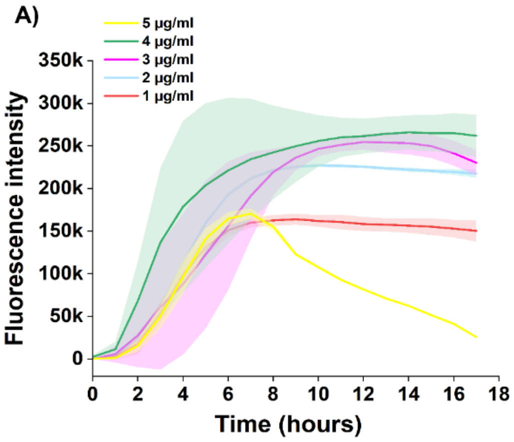

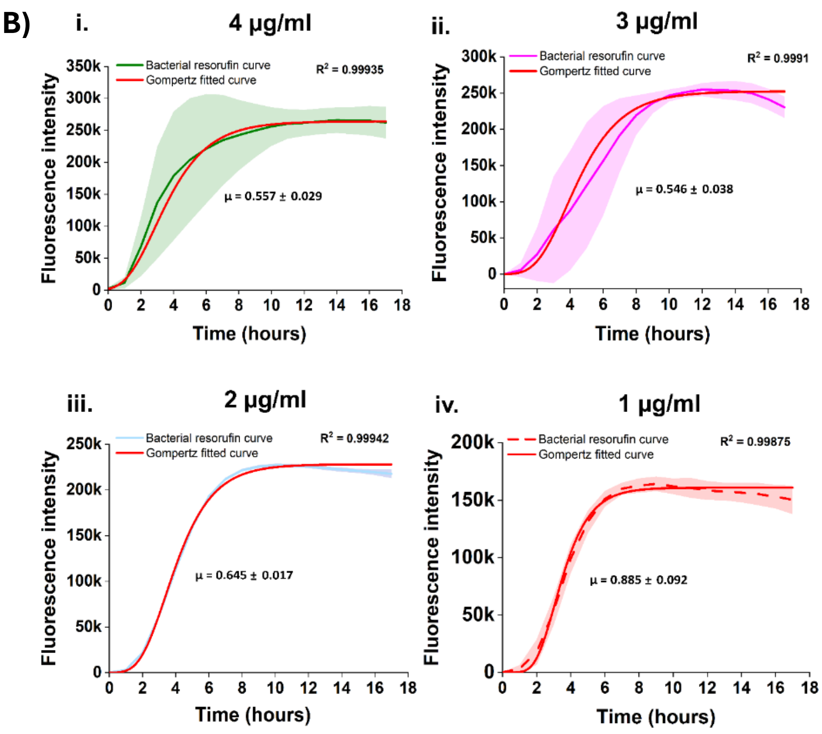


**Figure S5.** Optimization of resazurin reporter concentration and Gompertz modelling within multifunctional NaAMPS hydrogels. **(A)** Real-time fluoometric trajectories of uropathogenic E. coli tracking over a 17-hour timeline across varying initial resazurin concentration profiles (1,2,3,4, and 5 μg/mL). Shaded areas denote the standard deviation of experimental replicates (n=3). The highest concentration (5 μg/mL, yellow line) exhibits a clear distinction from standard sigmoidal growth, displaying an anomalous bell-shaped kinetic trajectory with late-stage signal decay. (**B) i-iv.** Non-linear regression profiles and classical Gompertz growth model fits (red lines) for the standard sigmoidal concentration regimes: **(i)** 4 μg/mL, showcasing robust mathematical convergence ($R^2$ = 0.99935) and an accumulation rate of μ=0.557 ± 0.029; **(ii)** 3 μg/mL ($R^2$ = 0.9991, μ=0.546 ± 0.038); **(iii)** 2 μg/mL ($R^2$ = 0.99942, μ = 0.645 ± 0.017); and **(iv)** 1 μg/mL ($R^2$ = 0.99875, μ=0.885 ± 0.092). The high coefficients of determination ($R^2$ > 0.99) validate stable, highly predictable conversion mechanics across the 1 to 4 μg/mL operational window, confirming that the network architecture maintains a non-destructive susbtrate supply up to its terminal plateau before the introduction of high-density resazurin constraints.

| Resazurin concentration | Uropathogenic *E. coli* growth rate (µ) | Resorufin intensity at 18 hours |
|---|---|---|
| 1 µg/mL | 0.885 ± 0.092 | 150K ± 12558 |
| 2 µg/mL | 0.645 ± 0.017 | 217K ± 4668 |
| 3 µg/mL | 0.546 ± 0.038 | 230 ± 14776 |
| 4 µg/mL | 0.557 ± 0.029 | 262 ± 24690 |

**Table S2.** Uropathogenic E. coli metabolic growth rates and terminal resorufin fluorescence intensities across varying resazurin reporter concentrations. Data represent the mean ± standard deviation (n=3). Specific signal accumulation rates (µ) were determined by fitting experimental real-0time fluorometric data to the classical Gompertz growth model ($R^2$ > 0.99). The 5 µg/mL experimental group is omitted from calculation due to its non-sigmoidal, bell-shaped profile.

| Resazurin concentration | Signal-to-noise (SNR) ratio at 2 hours | Uropathogenic *E. coli* growth rate (µ) |
|---|---|---|
| 1 µg/mL | 86.58 | 0.885 ± 0.092 |
| 2 µg/mL | 82.44 | 0.645 ± 0.017 |
| 3 µg/mL | 278.81* | 0.546 ± 0.038 |
| 4 µg/mL | 26.45 | 0.557 ± 0.029 |

**Table S3.** Early-stage (2 hours) signal-to-noise (SNR) and corresponding metabolic growth rates as a function of hydrogel resazurin loading. SNR were captured during the initial diagnostic window (t=2 hours) to evaluate early assay performance. Growth rates µ represent mean ± standard deviation (n=3). The asterisk (*) identifies an isolated peak in SNR resulting from a sharp, transient fluorescence differential between infected matrices and uninoculated baseline control hydrogels at the specific time interval.

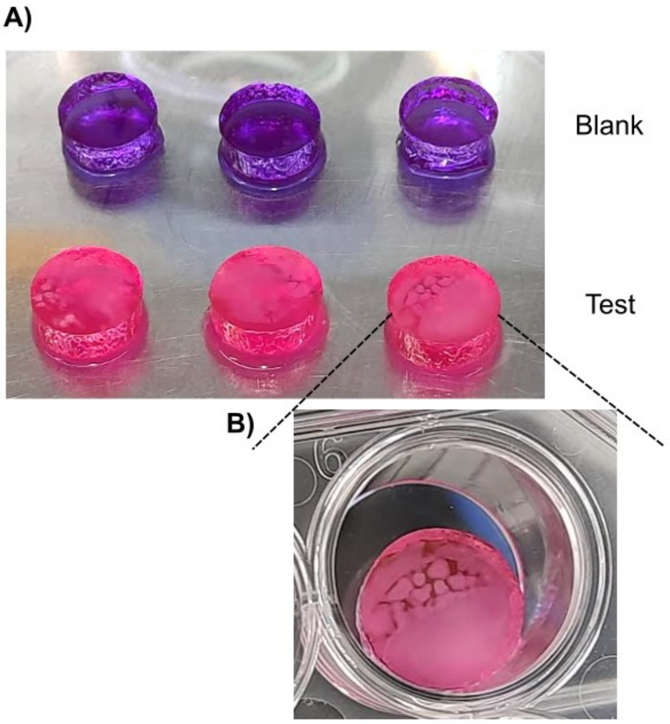


**Figure S6.** Visual macro-scale colorimetric response and surface colony distribution of *S. aureus* on NaAMPS hydrogels. **(A)** Macroscopic overview of multifunctional NaAMPS hydrogel discs following assay incubation. The top row (blank) demonstrates the stable, unreacted purple baseline of the resazurin-loaded matrix in the absence of infection. The bottom row (test) shows the robust, uniform colorimetric shift to pink indicating metabolic reduction to resoruin by *S. aureus*. **(B)** Magnified well-plate view of a positive test hydrogel disc, showing visible, distinct *S. aureus* macro-colonies across the hydrogel matrix surface post-detection.

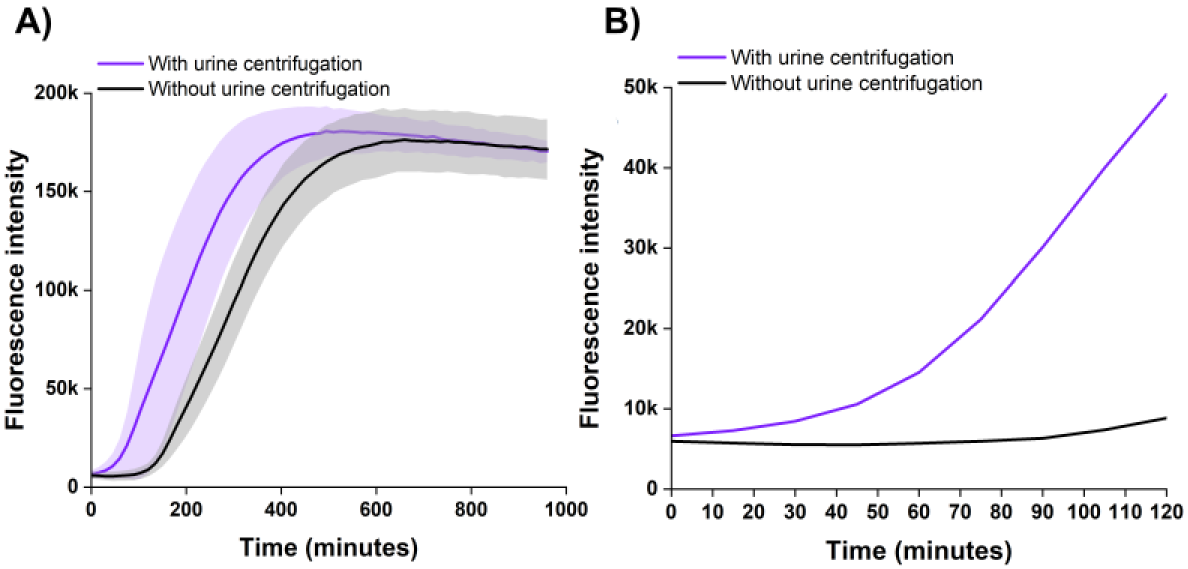


**Figure S7.** Influence of pre-analytical urine centrifugation on metabolic detection kinetics and turnaround time. **(A)** Full 1000-minute real-time fluorometric profiles of synthetic urine samples spiked with uropapthogenic E. coli processed with a pre-test centrifugation step (purple line) versus raw, uncentrifuged controls (black line). Shaded regions represent the standard deviation of experimental replicates (n=3). While both processing conditions ultimately converge on equivalent maximum fluorescence intensities at later stages, the centralised processing step yields an accelerated sigmoidal curve. **(B)** Maginifed kinetic view of the initial 120-minute diagnostic window. The centrifuged samples demonstrate an immediate, steep exponential signal departure from the baseline, while the raw, uncentrifuged matrix remains suppresed at baselines values. This highlights that concentrating uropathogenic loads at the hydrogel interface effectively eliminates early latency, significantly shortening the platform's Time-To-Detection (TTD).

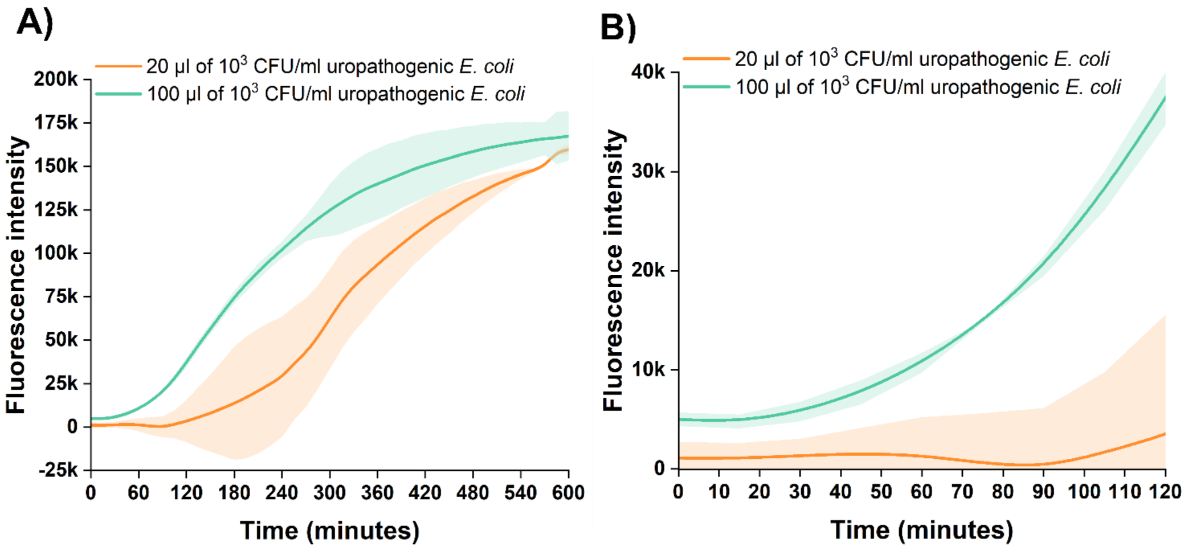


**Figure S8.** Effect of sample inoculation volume on the metabolic detection kinetics of low-concentration uorpathogenic E. coli. **(A)** Full 600-minute real-time fluorometric trajectories tracking a low-density bacterial load ($10^3$ CFU/mL) inoculated at volumes of 20 µL (orange line) versus 100 µL (green line) within the hydrogel sensing platform. Shaded regions signify the standard deviation across independent experimental replicates (n=3). **(B)** Magnified kinetic view of the data presented in panel A across the 120-minute diagnostic phase. The 100 µL volume configuration demonstrates an accelerated baseline departure within 60 minutes, driving strong fluorescence accumulation by 2 hours. Conversely, the lower 20 µL exhibits a pronounced latent period, remaining unreduced at the baseline due to mass transport limitations and a lower initial absolute cell count at the hydrogel interface. This confirms that optimization of the initial sample volume presents an accessible approach to shorten diagnostic turnaround times for low bacterial concentrations.

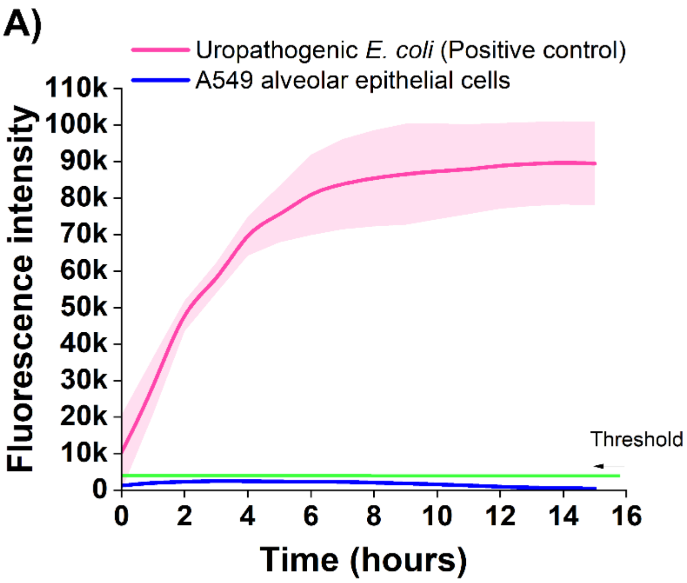


**Figure S9.** Diagnostic specificity and metabolic matrix interference validation using mammalian epithelial cell lines. Time-dependent fluorometric tracking of multifunctional NaAMPS hydrogels exposed to mammalian epithelial cells (A549 cell line) compared against uropathogenic E. coli positive control, and control baseline hydrogels. Inoculation with epithelial populations yields a flat, negligible kinetic profile that overlaps with the negative control baseline, demonstrating that background non-bacterial cellular matter does not cross-react with or prematurely reduce the encapsulated resazurin reporter. This selective metabolic restriction validates the platform's diagnostic accuracy for direct urinary pathogen deployment without risk of cell-mediated false positives.

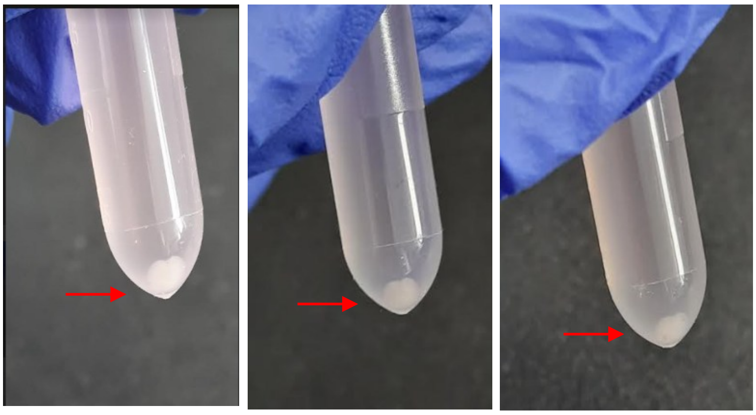

**Figure S10.** Visual confirmation of isolated bacterial DNA pellets recovered post-assay NaAMPS hydrogels. Images showing triplicate experimental replicates of dense, well-defined nucleic acid precipitates (indicated by red arrouws) during the downstream DNA extraction workflow. Images were captured immediately following initial precipitation steps and prior to the final ethanol addition/wash phases. The formation of these highly visible, substantial pellets demonstrates that the passive swelling-mediated bacterial elution process into 1XPBS successfully retrieves an abundance quantity of high-purity cellular biomass from the hydrogel network. This extraction yield provides an ideal, contaminant-free template concentration fro sensitive downstream genotypic analysis and molecular amplification.

| Samples | Concentration (ng/µL) (DNA eluted volume = 100 µL) | Purity (260/280 nm) |
|---|---|---|
| Replicate 1 | 25.892 | 1.789 |
| Replicate 2 | 27.690 | 1.898 |
| Replicate 3 | 33.251 | 1.845 |

**Table S4.** Quantitative yield and purity metrics of bacterial genomic DNA extracted from post-sensing NaAMPS hydrogels. All genomic DNA samples were eluted in a fixed final volume of 100 µL. Absorbance ratios of 260/280 nm falling between 1.8 and 1.9 signify high-purity nucleic acid recovery with negligible protein or matrix reagent co-purification. These quantitative yields match the visual pellet densities documents in Supplementary Figure S9 and confirm the preservation of genetic material optimized for direct downstream molecular diagnostic deployment.

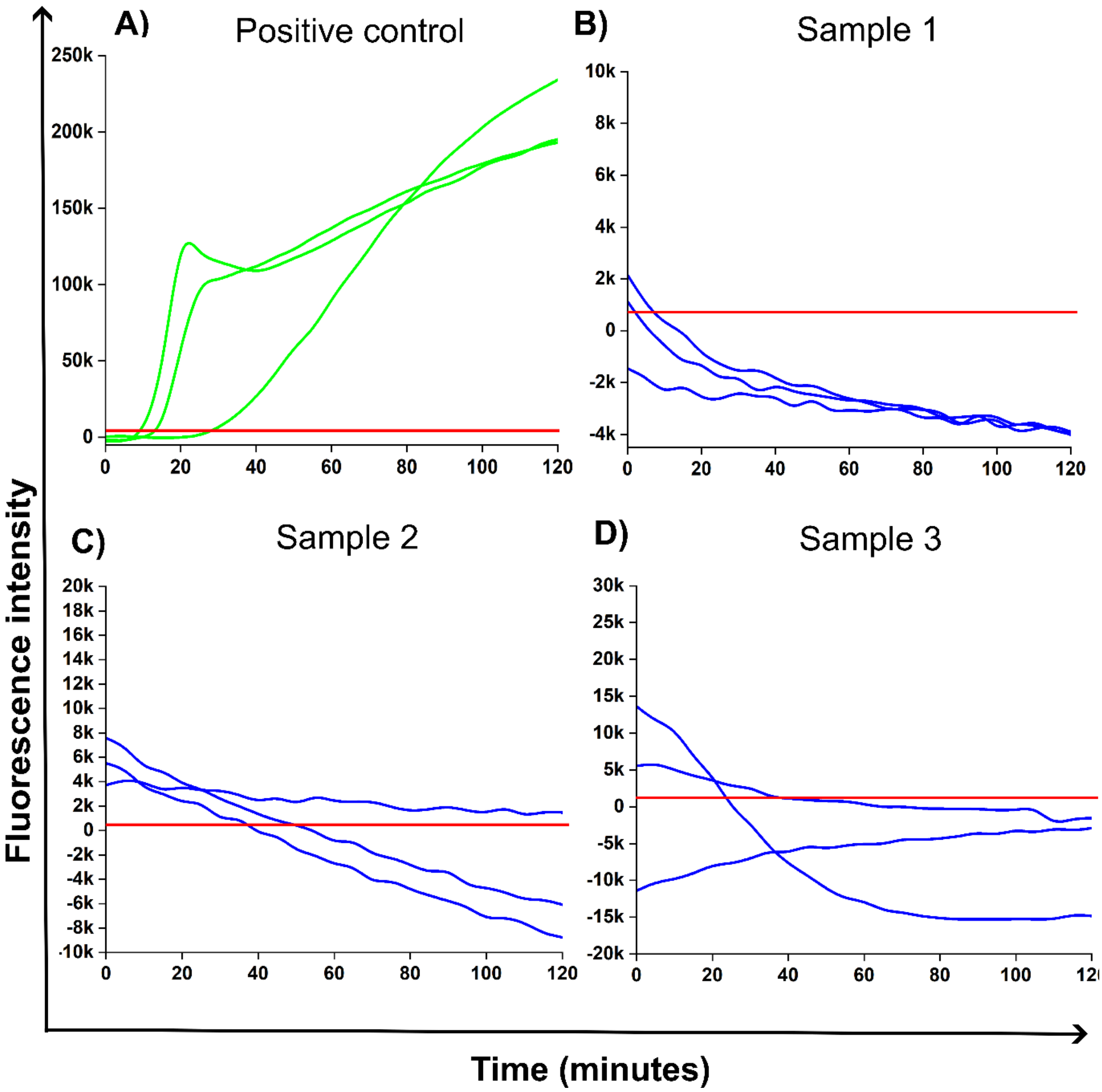


**Figure S11.** Real-time clinical urine sample screening and early kinetic diagnostic tracking within the high-throught matrix. Real-time fluorometric screening trajectories capturing the 120-minute window for clinical patient samples processed within the 96-well hydrogel diagnostic platform. **(A)** Positive control validation runs utilizing artificial urine spiked with confirmed clinical isolate uropathogenic E. coli, demonstrating rapid, exponential metabolic resazurin reduction and an immediate increase in signal from the baseline detection threshold (indicated by the solid red reference lines) within 15 to 35 minutes of sample incubation. **(B)** Sample 1, **(C)** Sample 2, and **(D)** Sample 3 present negative clinical urine patient profiles. Across entire 2-hour monitoring phase, the metabolic fluorescence trajectories remain completely suppressed below or within the initial baseline threshold boundaries. The mild downward sloped in relative fluorescence units (RFU) characterize typical non-specific background matrix equilibration in the absence of active bacterial replication, establishing a clear, instrument-read separation between positive and negative specimens.

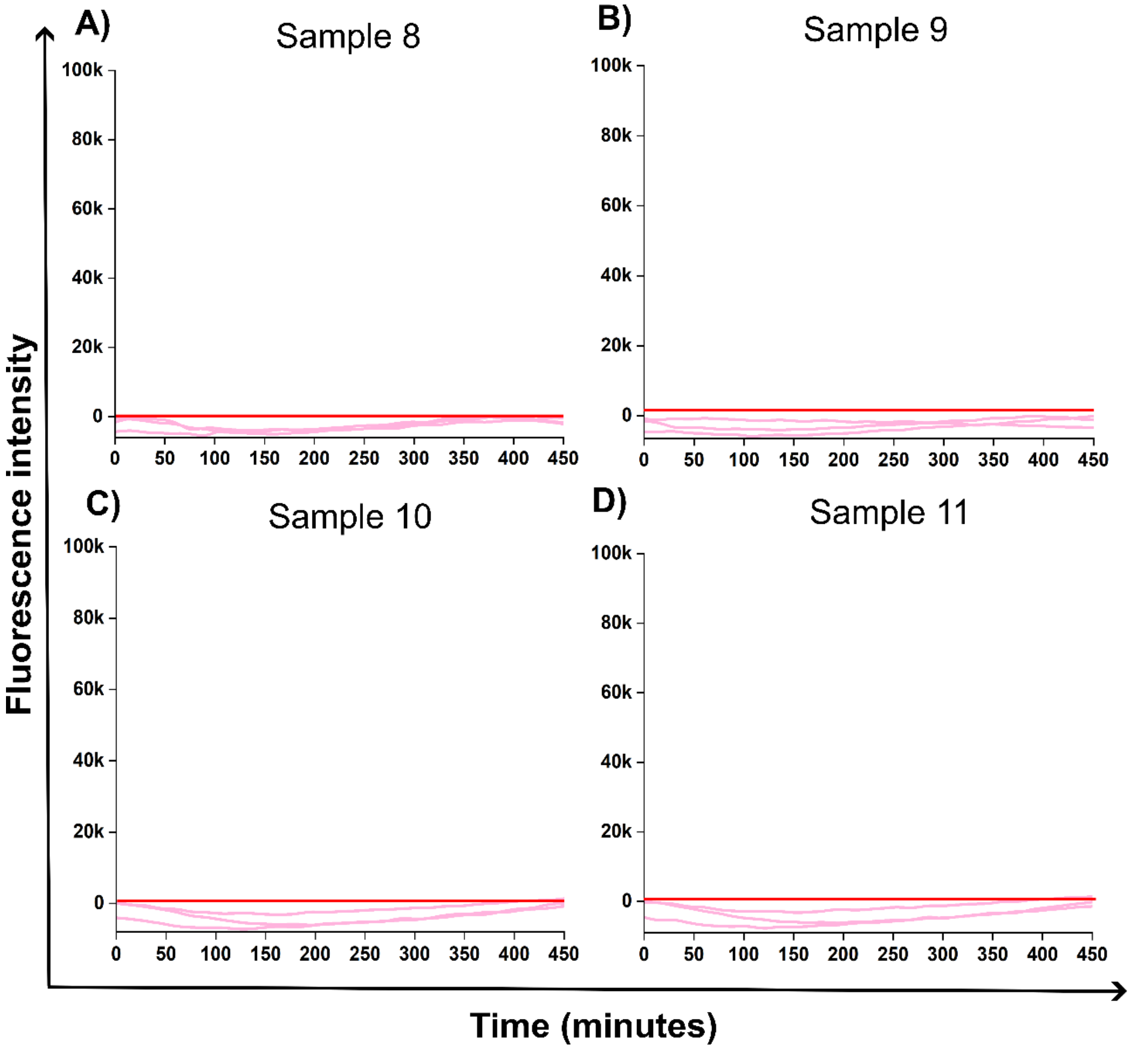


**Figure S12.** Sample 8 **(A)**, Sample 9 **(B)**, Sample 10 **(C)**, and Sample 11 **(D)** = no UTI detected.

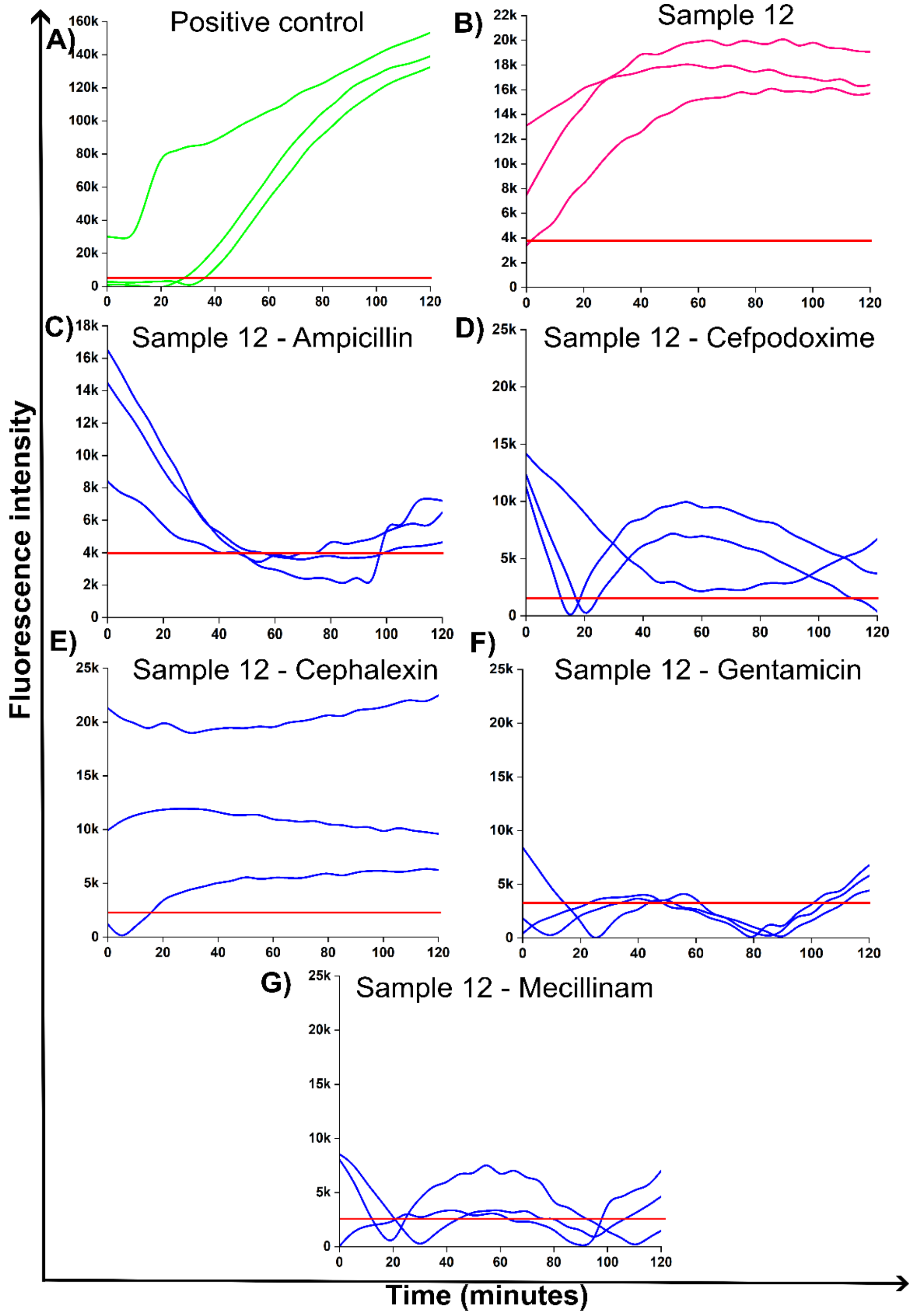


**Figure S13. A)** Positive control = uropathogenic *E. coli*, **B)** Sample 12 = UTI, **C)** Sample 12 = Ampicillin sensitivity, **D)** Sample 12 = Cefpodoxime resistance, **E)** Sample 12 = Cephalexin sensitivity, **F)** Sample 12 = sensitivity, and **G)** Sample 12 = Mecillinam sensitivity.

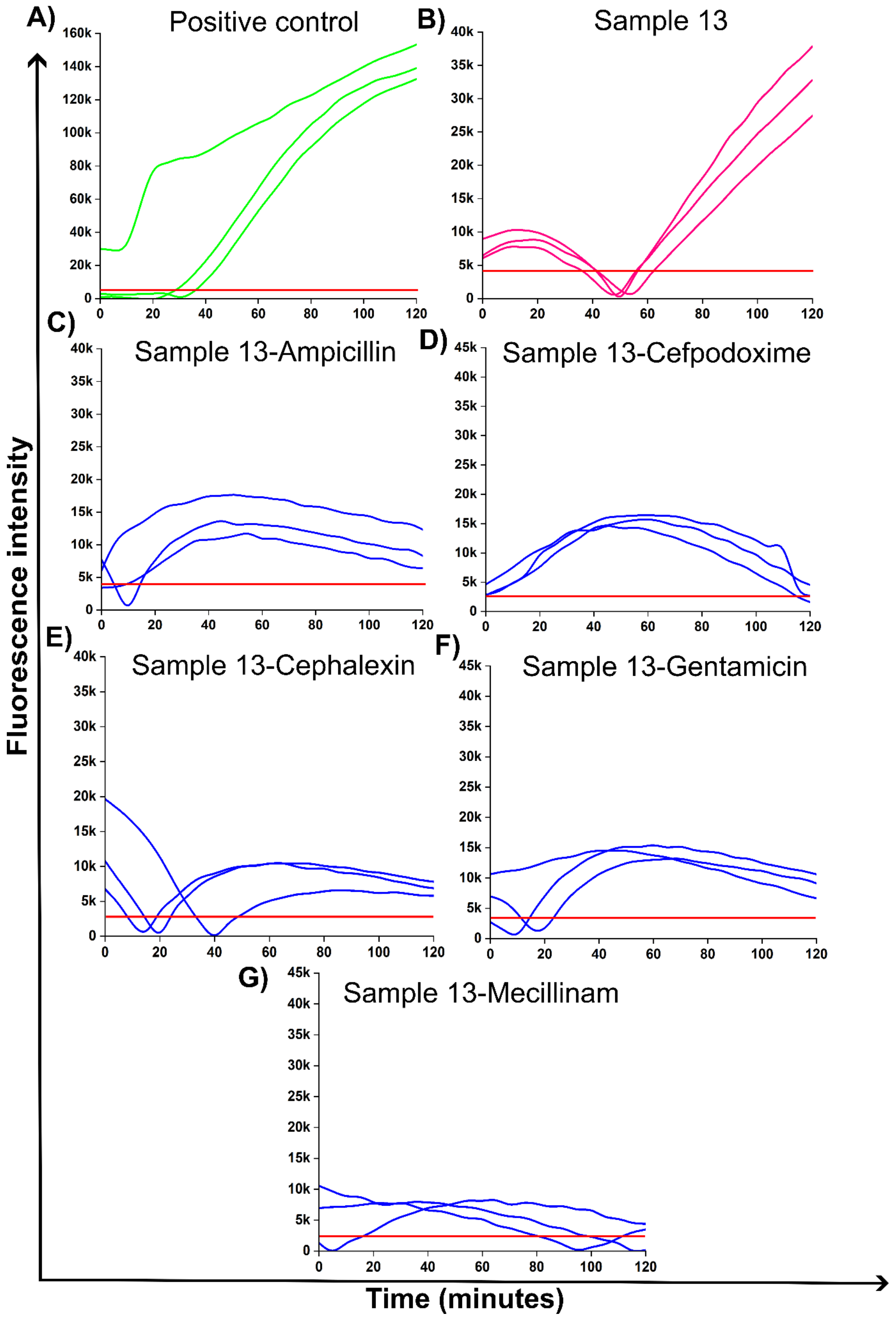


**Figure S14 .A)** Positive control = uropathogenic *E. coli*, **B)** Sample 13 = UTI, **C)** Sample 13 = Ampicillin resistance, **D)** Sample 13 = Cefpodoxime resistance, **E)** Sample 13 = Cephalexin resistance, **F)** Sample 13 = Gentamicin resistance, and **G)** Sample 13 = Mecillinam sensitivity.

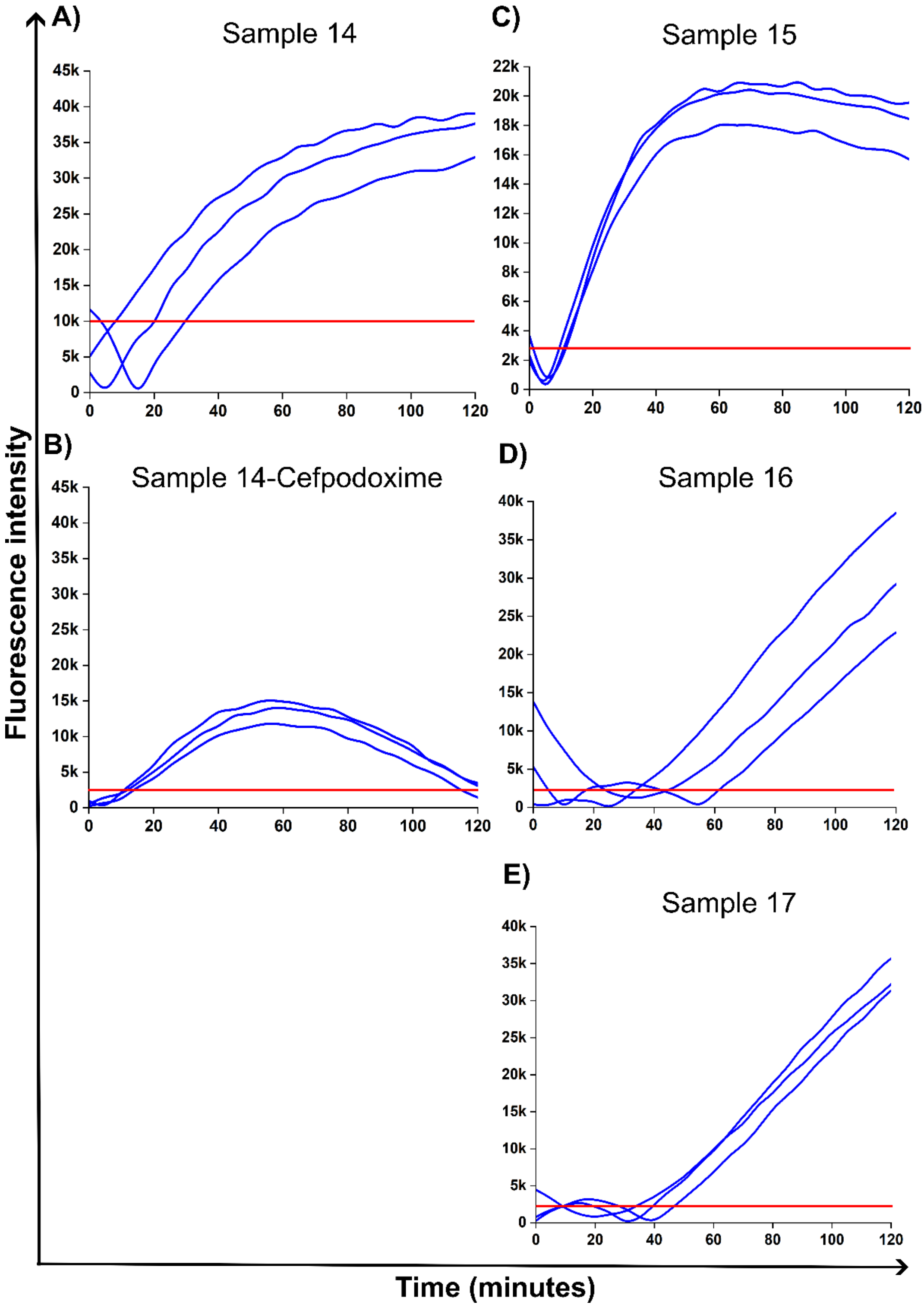


**Figure S15. A)** Sample 14 = UTI, **B)** Sample 14 = Cefpodoxime sensitivity, **C)** Sample 15 = UTI, **D)** Sample 16 = UTI, and **E)** Sample 17 = UTI.

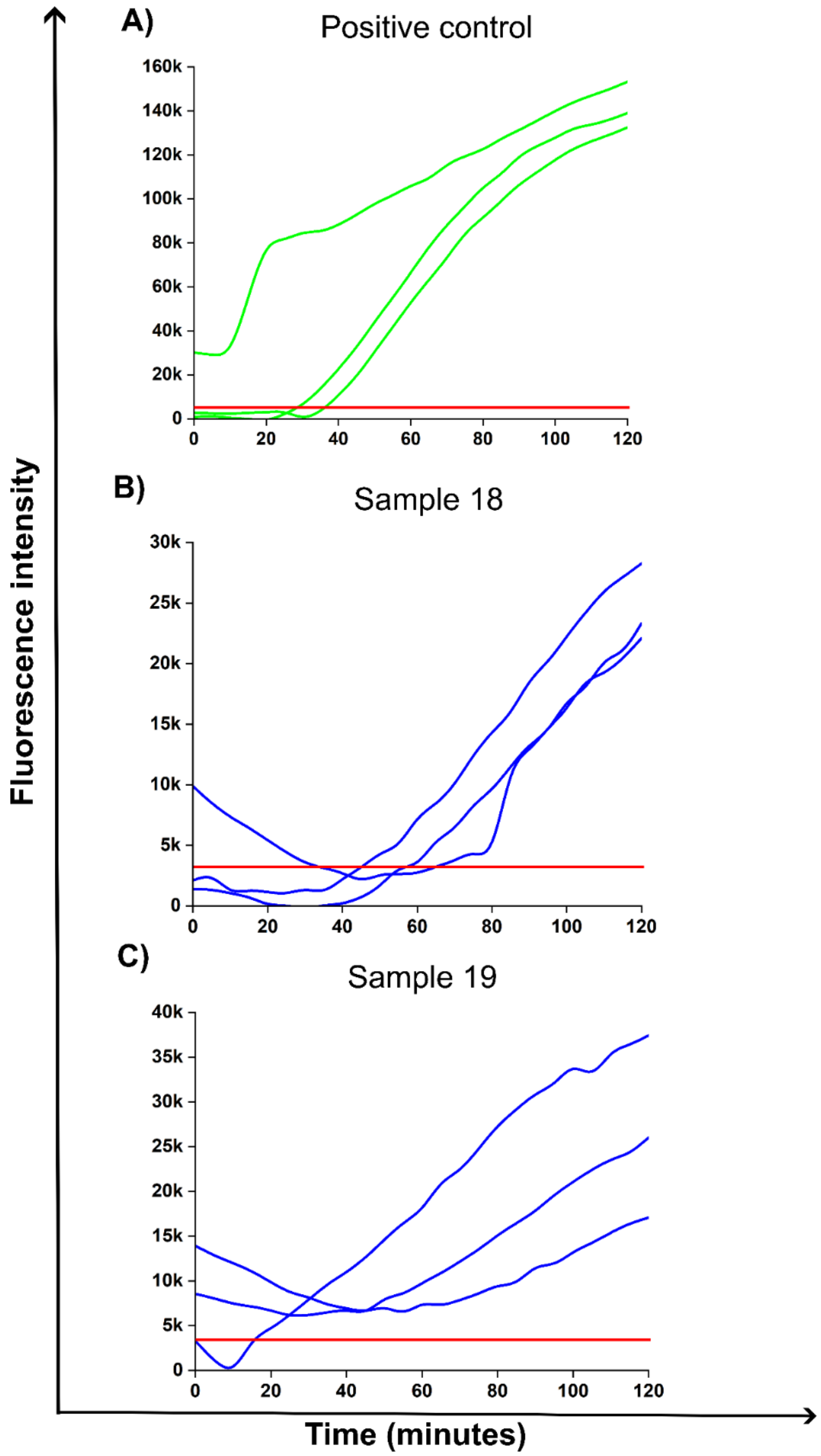


**Figure S16. A)** Positive control, **B)** Sample 18 = UTI and **C)** Sample 19 = UTI.

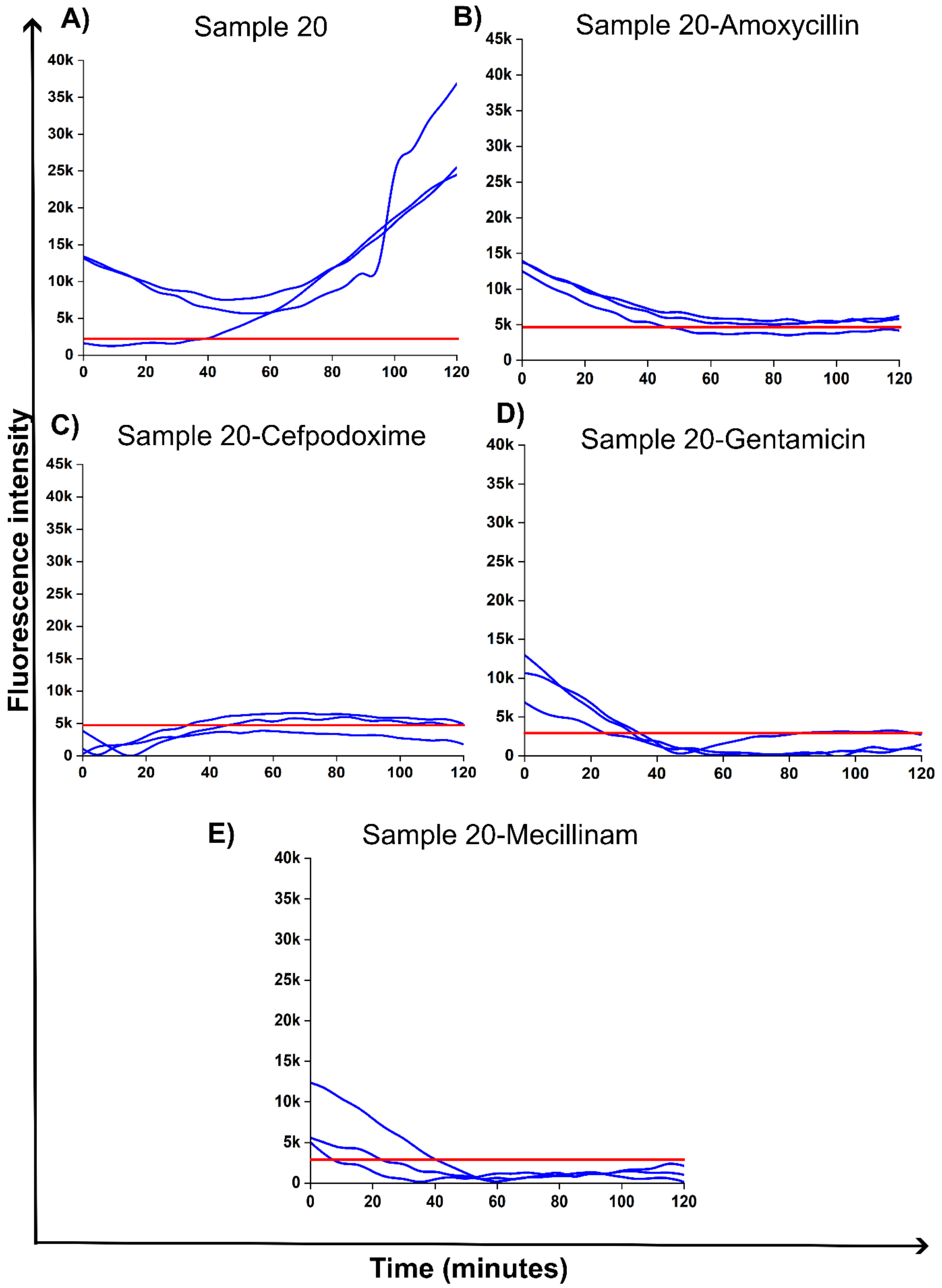


**Figure S17. A)** Sample 20 = UTI, **B)** Sample 20 – Amoxicillin sensitivity, **C)** Sample 20 – Cefpodoxime sensitivity, **D)** Sample 20 – Gentamicin sensitivity, and **E)** Sample 20 – Mecillinam sensitivity.

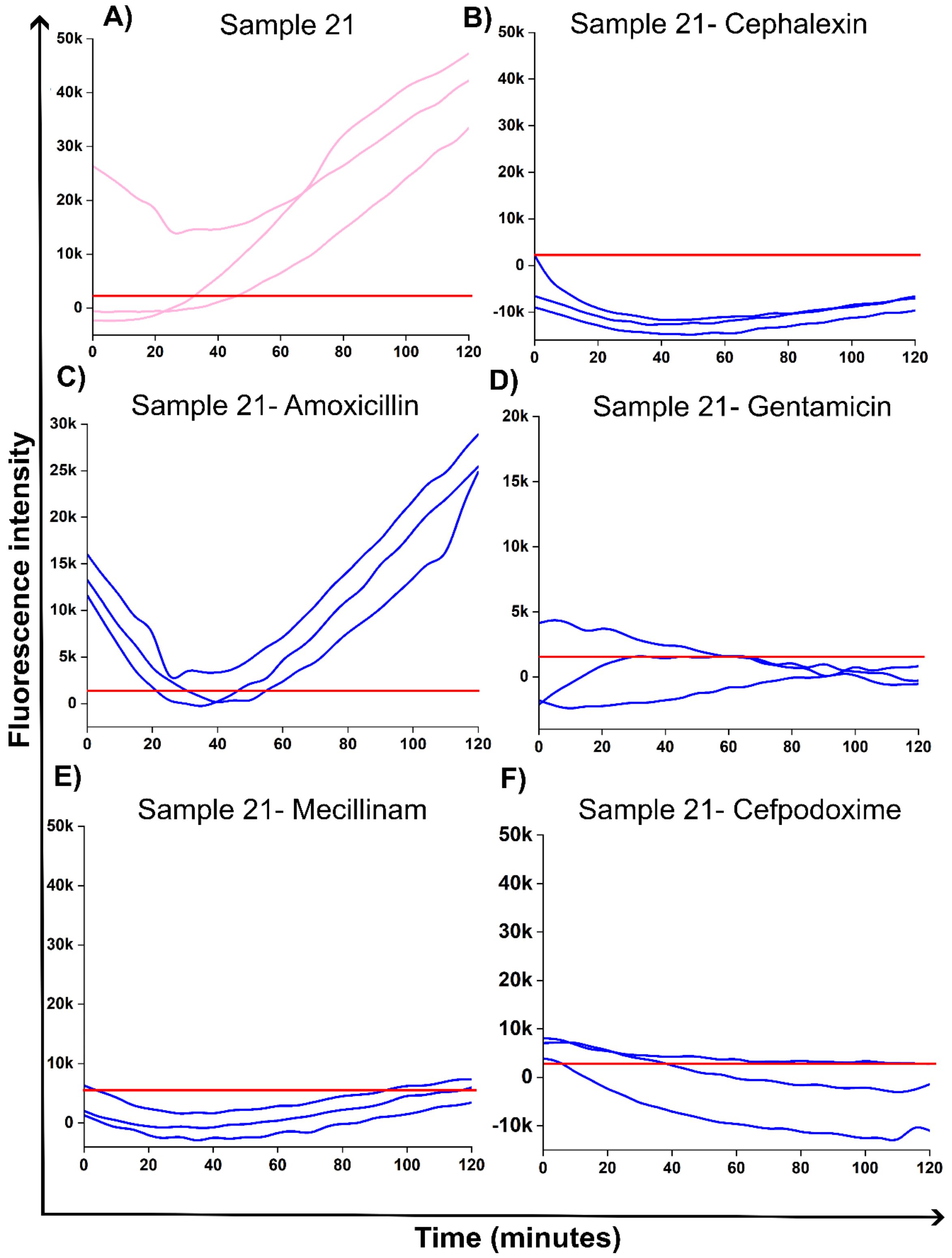


**Figure S18. A)** Sample 21 = UTI, **B)** Sample 21 = Cephalexin sensitivity, **C)** Sample 21 = Amoxicillin resistance, **D)** Sample 21 = Gentamicin sensitivity, **E)** Sample 21 = Mecillinam sensitivity, and **F)** Sample 21 = Cefpodoxime sensitivity.

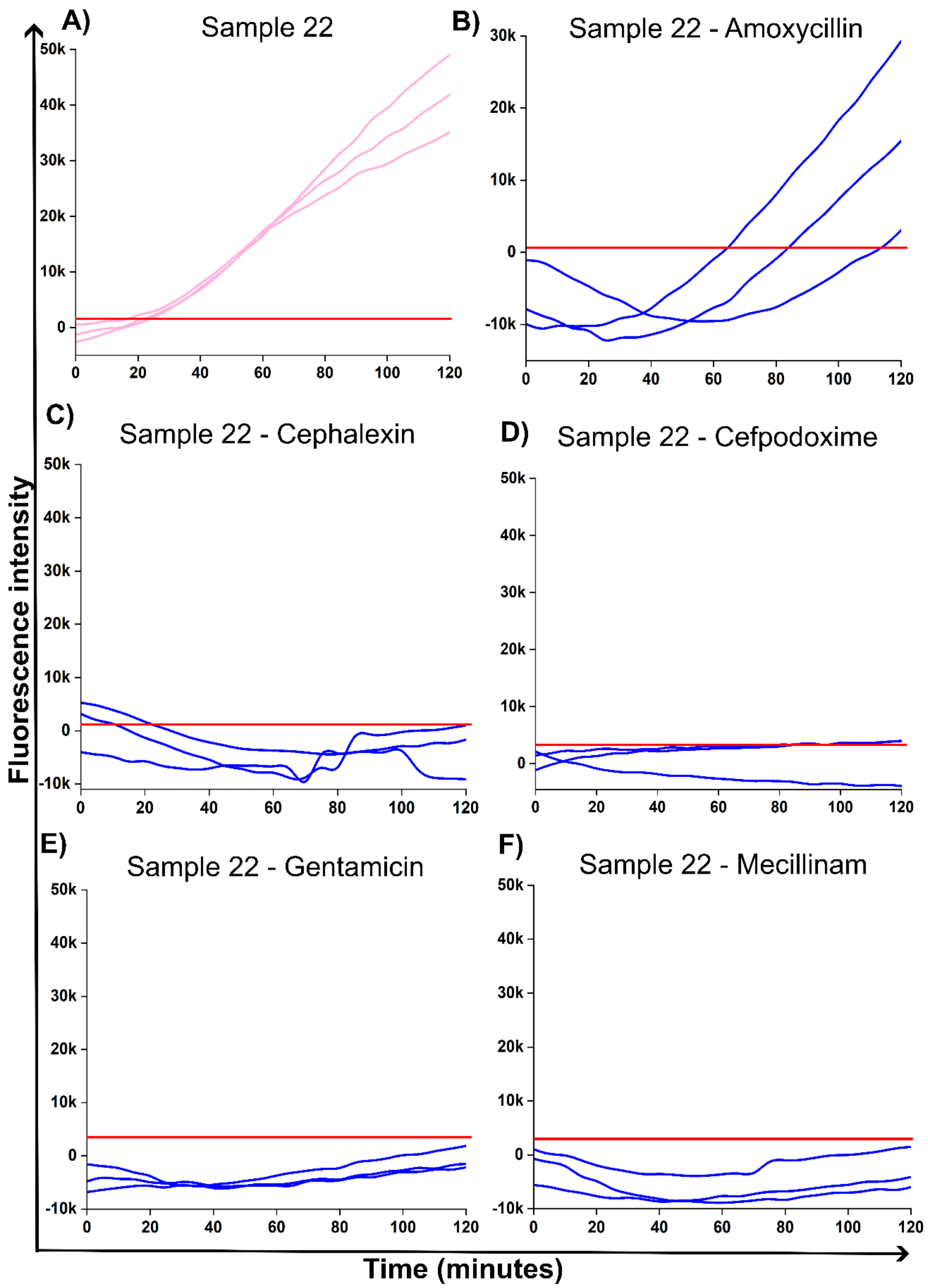


**Figure S19. A)** Sample 22 = UTI, **B)** Sample 22 = Amoxicillin resistance, **C)** Sample 22 = Cephalexin sensitivity, **D)** Sample 22 = Cefpodoxime sensitivity, **E)** Sample 22 = Gentamicin sensitivity, and **F)** Sample 22 – Mecillinam sensitivity.

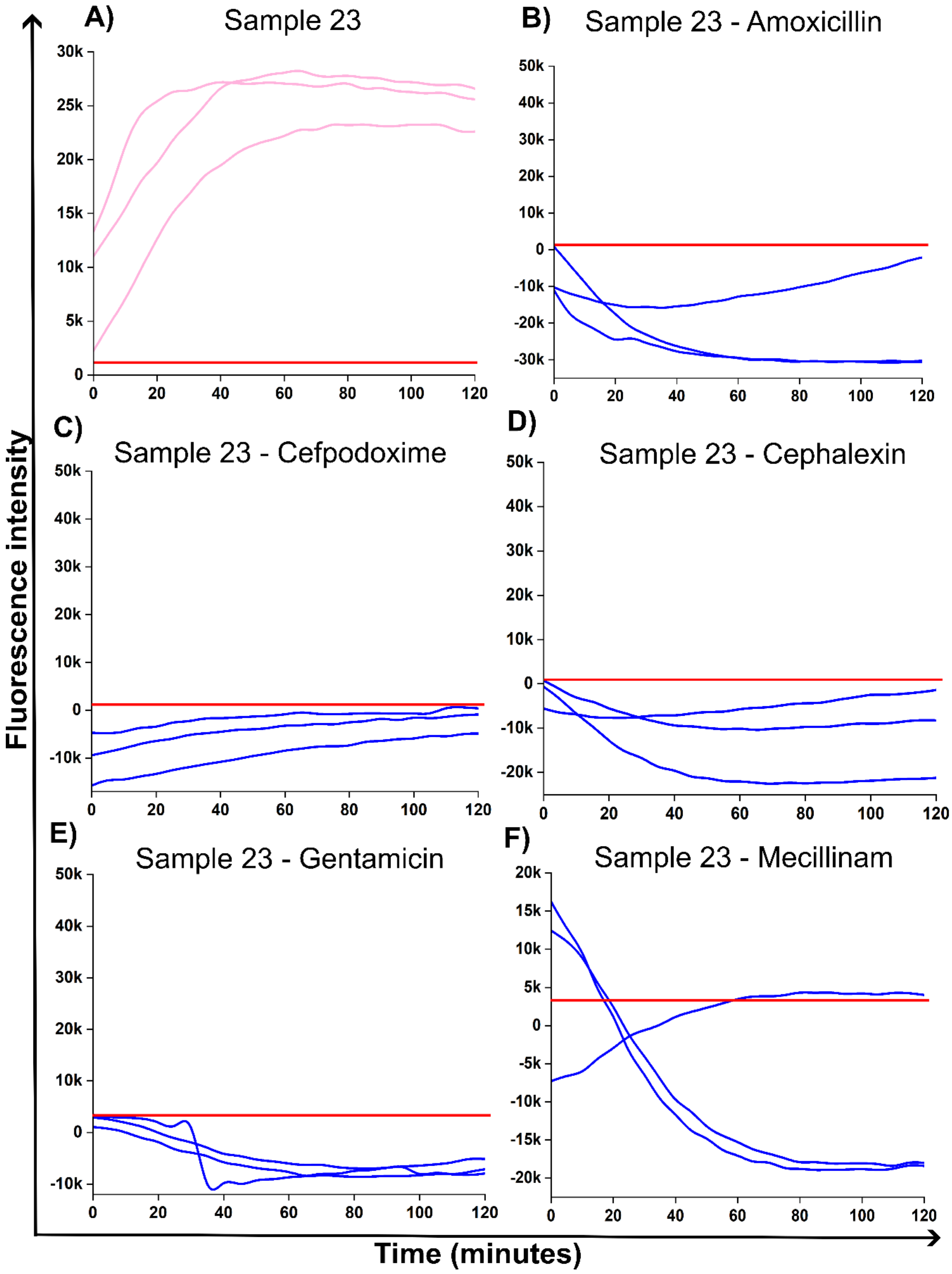


**Figure S20. A)** Sample 23 = UTI, **B)** Sample 23 = Amoxicillin sensitivity, **C)** Sample 23 = Cephalexin sensitivity, **D)** Sample 23 = Cefpodoxime sensitivity, **E)** Sample 23 = Gentamicin sensitivity, and **F)** Sample 23 – Mecillinam sensitivity

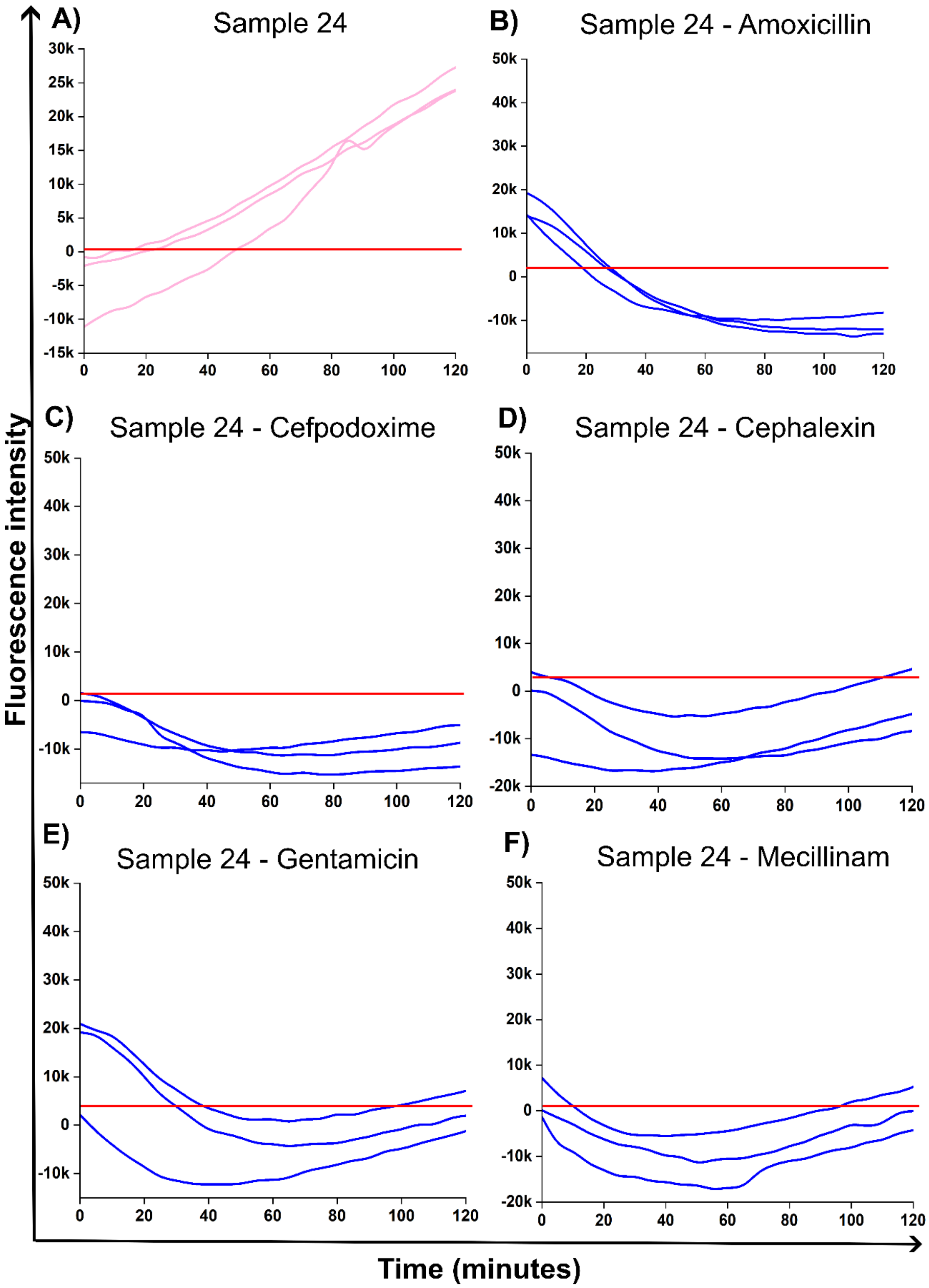


**Figure S21. A)** Sample 24 = UTI, **B)** Sample 23 = Amoxicillin sensitivity, **C)** Sample 24 = Cephalexin sensitivity, **D)** Sample 24 = Cefpodoxime sensitivity, **E)** Sample 24 = Gentamicin sensitivity, and **F)** Sample 24 – Mecillinam sensitivity

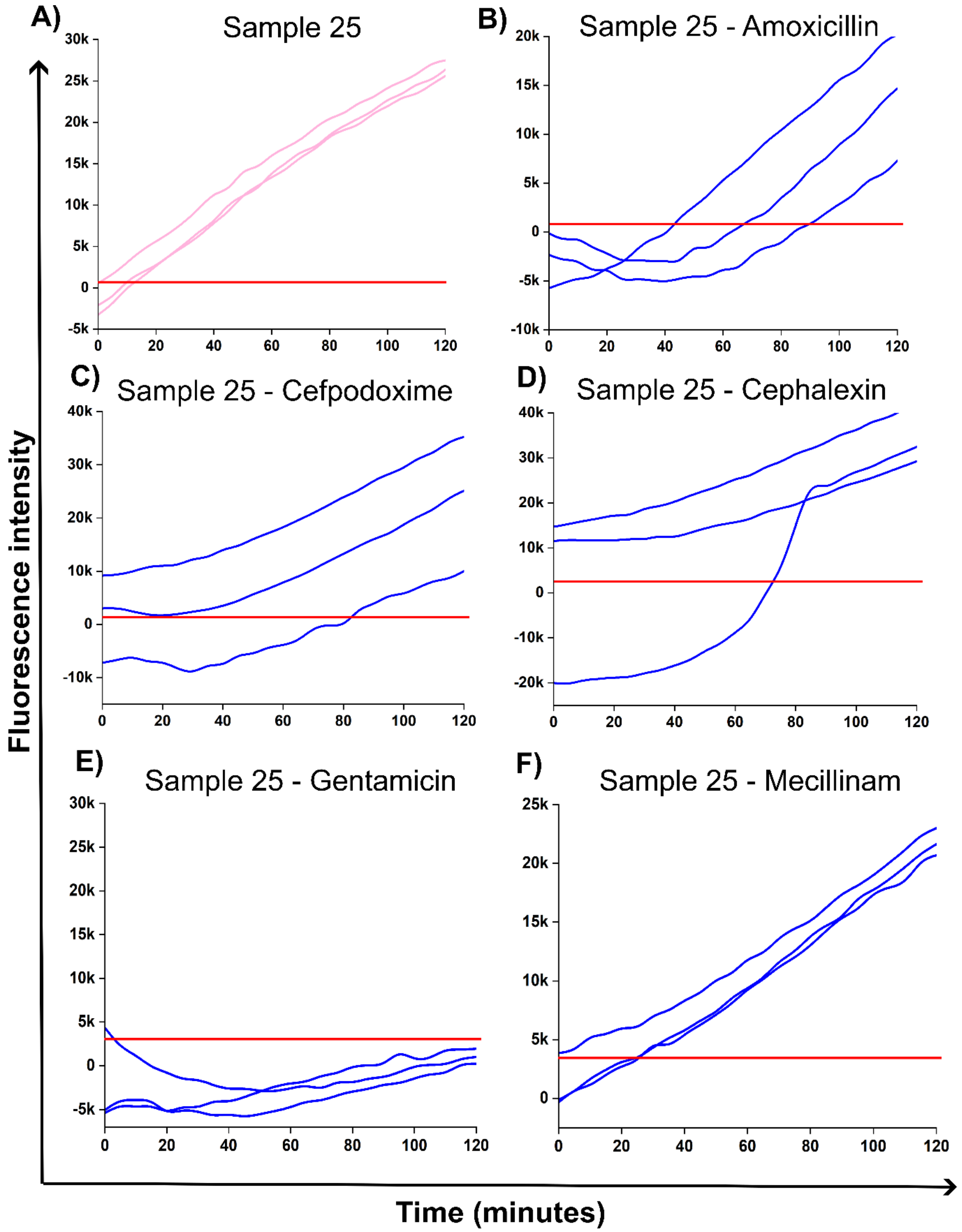


**Figure S22. A)** Sample 25 = UTI, **B)** Sample 25 = Amoxicillin resistance, **C)** Sample 25 = Cefpodoxime resistance, **D)** Sample 25 = Cephalexin resistance, **E)** Sample 25 = Gentamicin sensitivity, and **F)** Sample 25 = Mecillinam resistance.

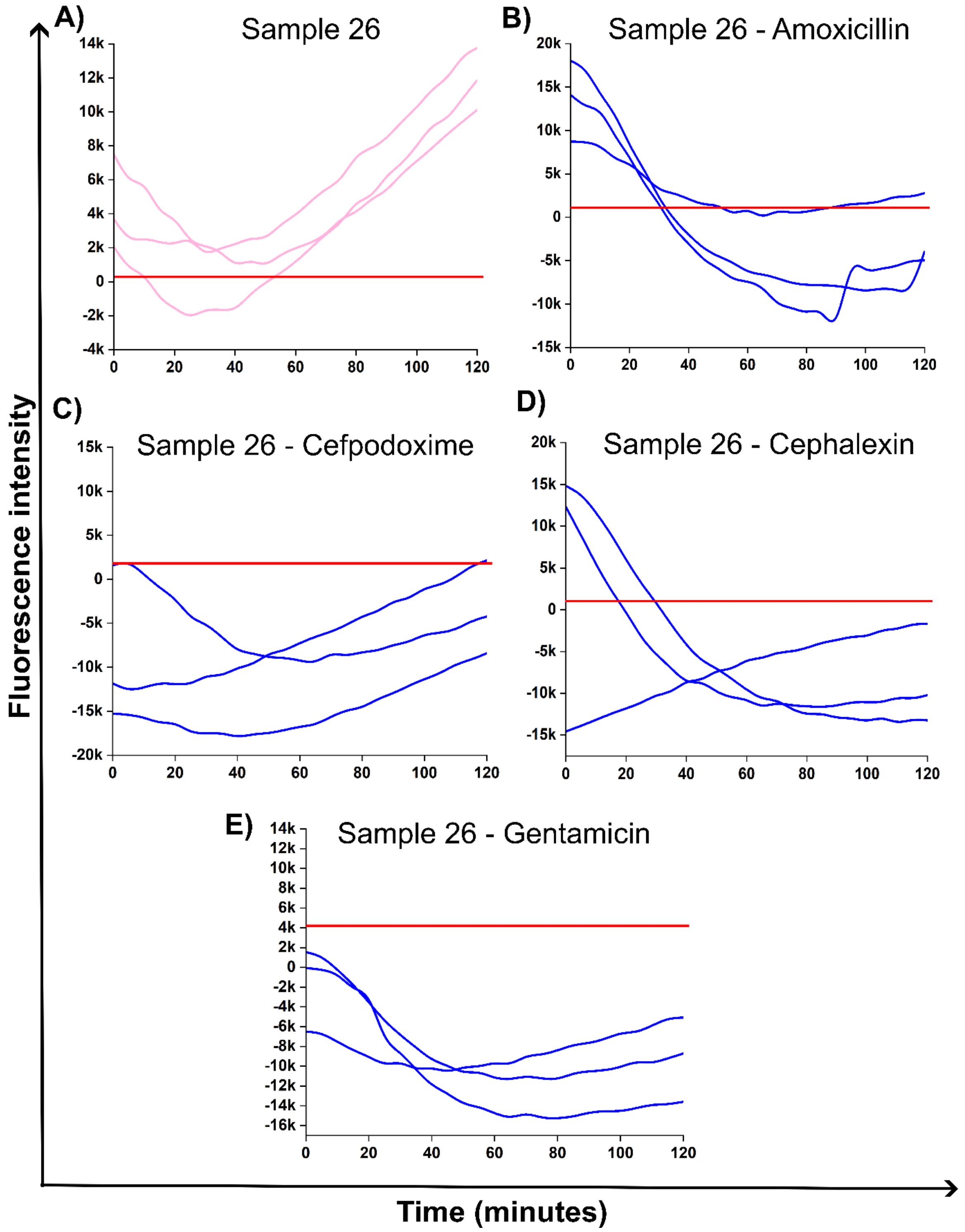


**Figure S23.** Clinical validation of *microAmp Dx.* **A)** Sample 26 = UTI, **B)** Sample 26 = Amoxicillin sensitivity, **C)** Sample 26 = Cefpodoxime sensitivity, **D)** Sample 26 = Cephalexin sensitivity, and **E)** Sample 26 = Gentamicin sensitivity.

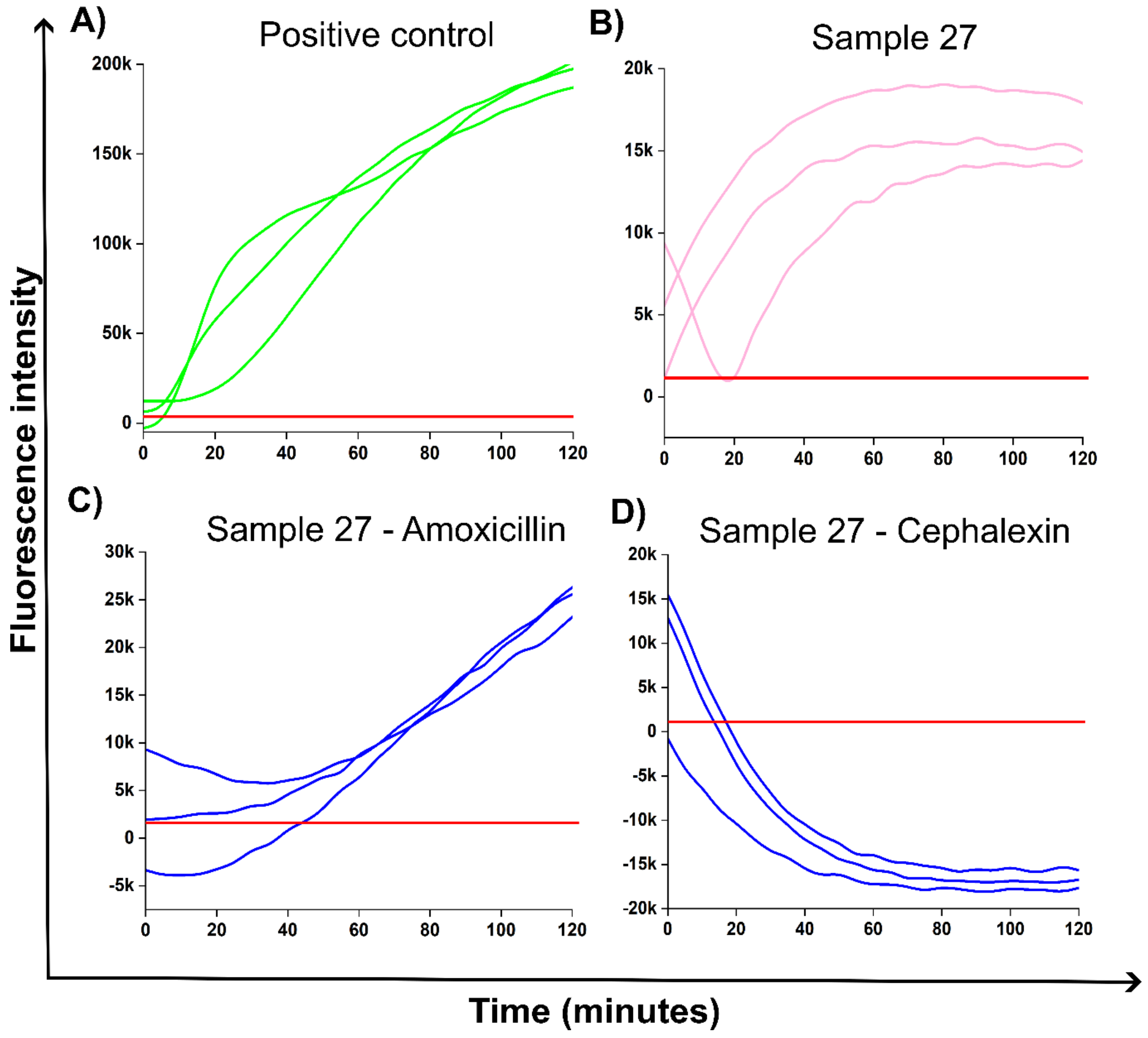


**Figure S24. A)** Positive control, **B)** Sample 27 = UTI, **C)** Sample 27 = Amoxicillin resistance, and **D)** Sample 27 = Cephalexin sensitivity.

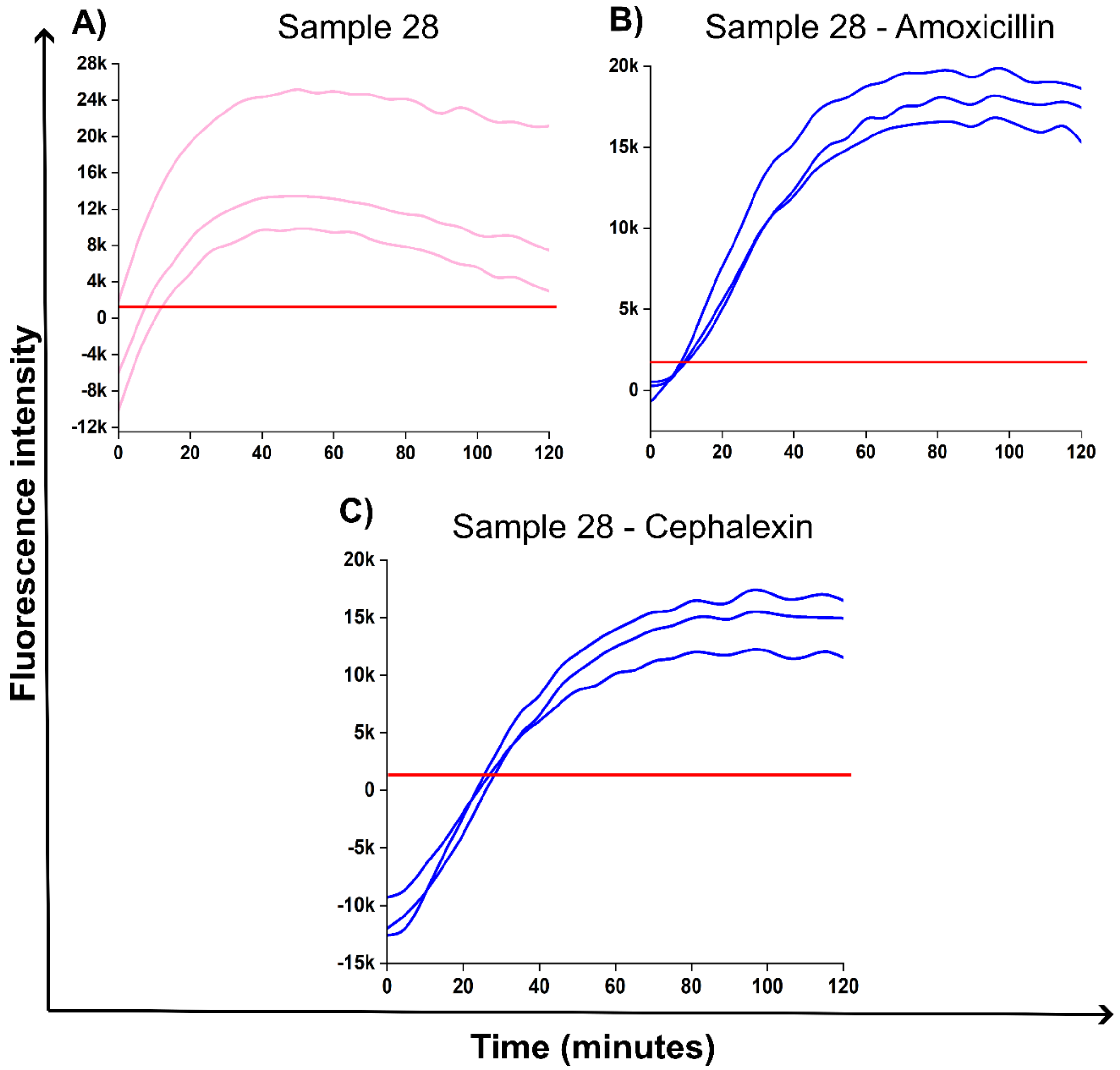


**Figure S25.** **A)** Sample 28 = UTI, **B)** Sample 28 = Amoxicillin resistance, and **C)** Sample 28 = Cephalexin resistance.

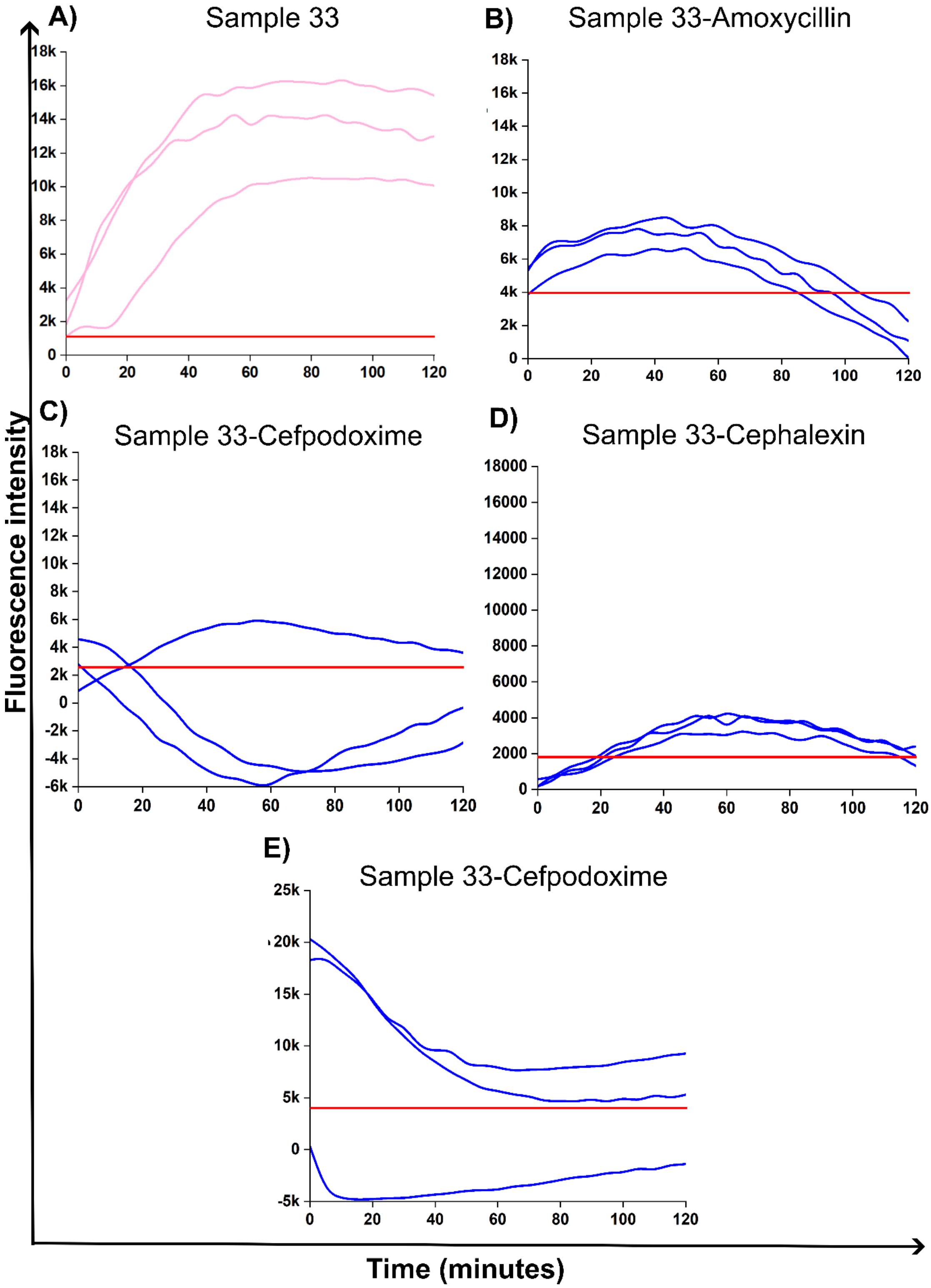


**Figure S26.** **A)** Sample 33 = UTI, **B)** Sample 33 = Amoxicillin sensitivity, and **C)** Sample 33 = Cefpodoxime sensitivity, **D)** Sample 33 = Cephalexin sensitivity, and **E)** Sample 33 = Gentamicin sensitivity.

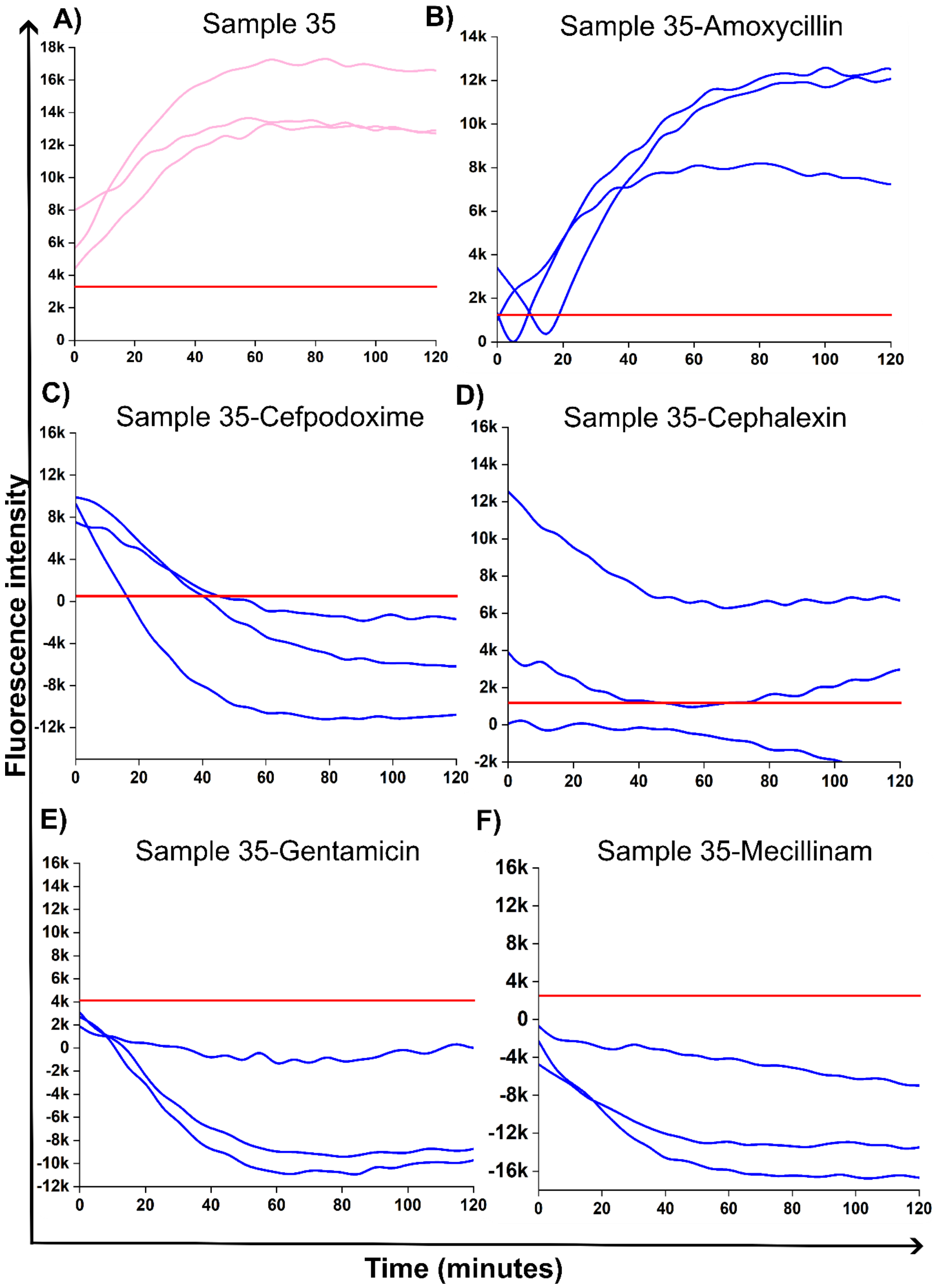


**Figure S27.** **A)** Sample 35 = UTI, **B)** Sample 35 = Amoxicillin resistance, and **C)** Sample 35 = Cefpodoxime sensitivity, **D)** Sample 35 = Cephalexin sensitivity, **E)** Sample 35 = Gentamicin sensitivity, and **F)** Sample 35 = Mecillinam sensitivity.

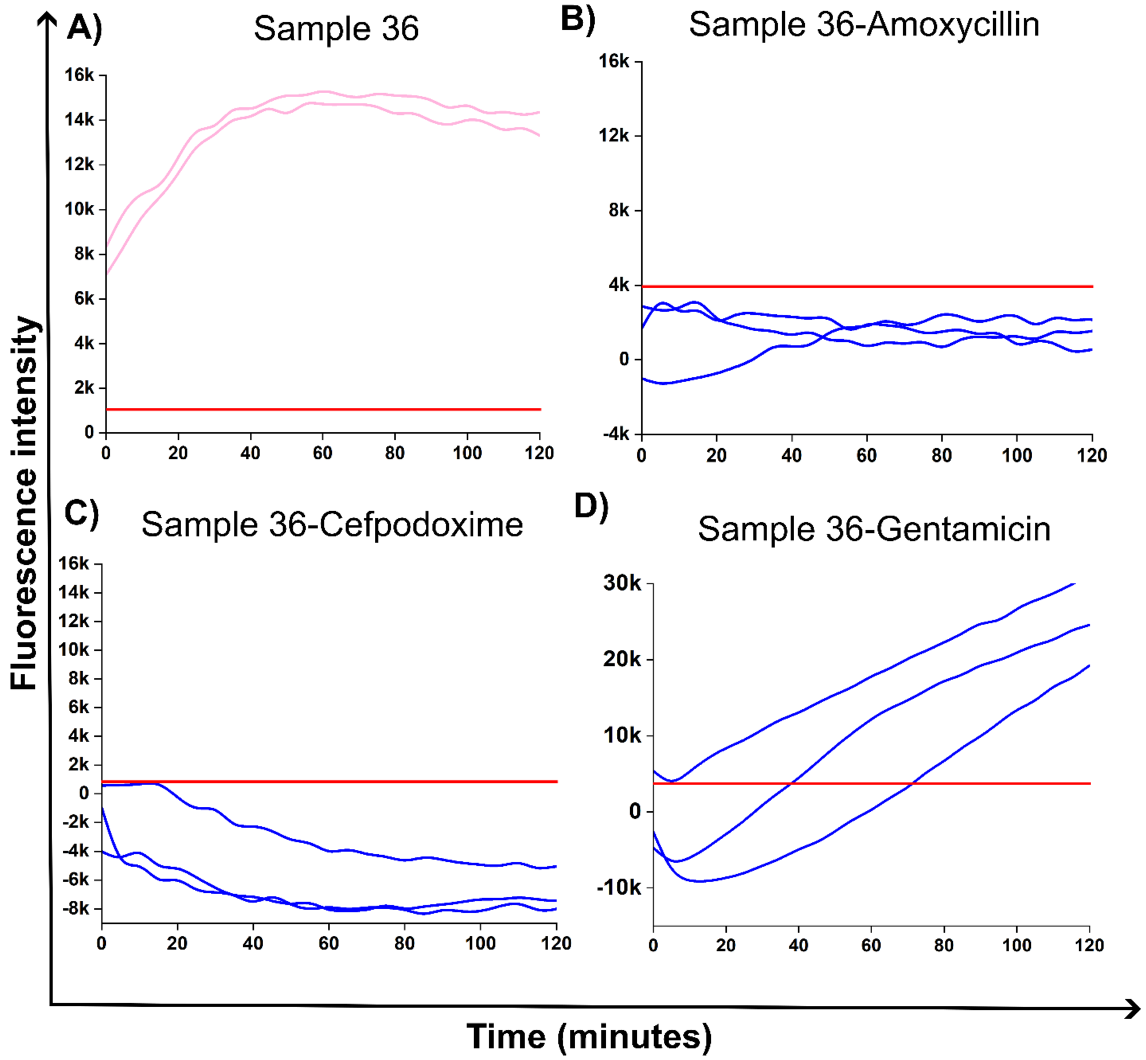


**Figure S28.** **A)** Sample 36 = UTI, **B)** Sample 36 = Amoxicillin sensitivity, and **C)** Sample 36 = Cefpodoxime sensitivity, and **D)** Sample 36 = Gentamicin resistance

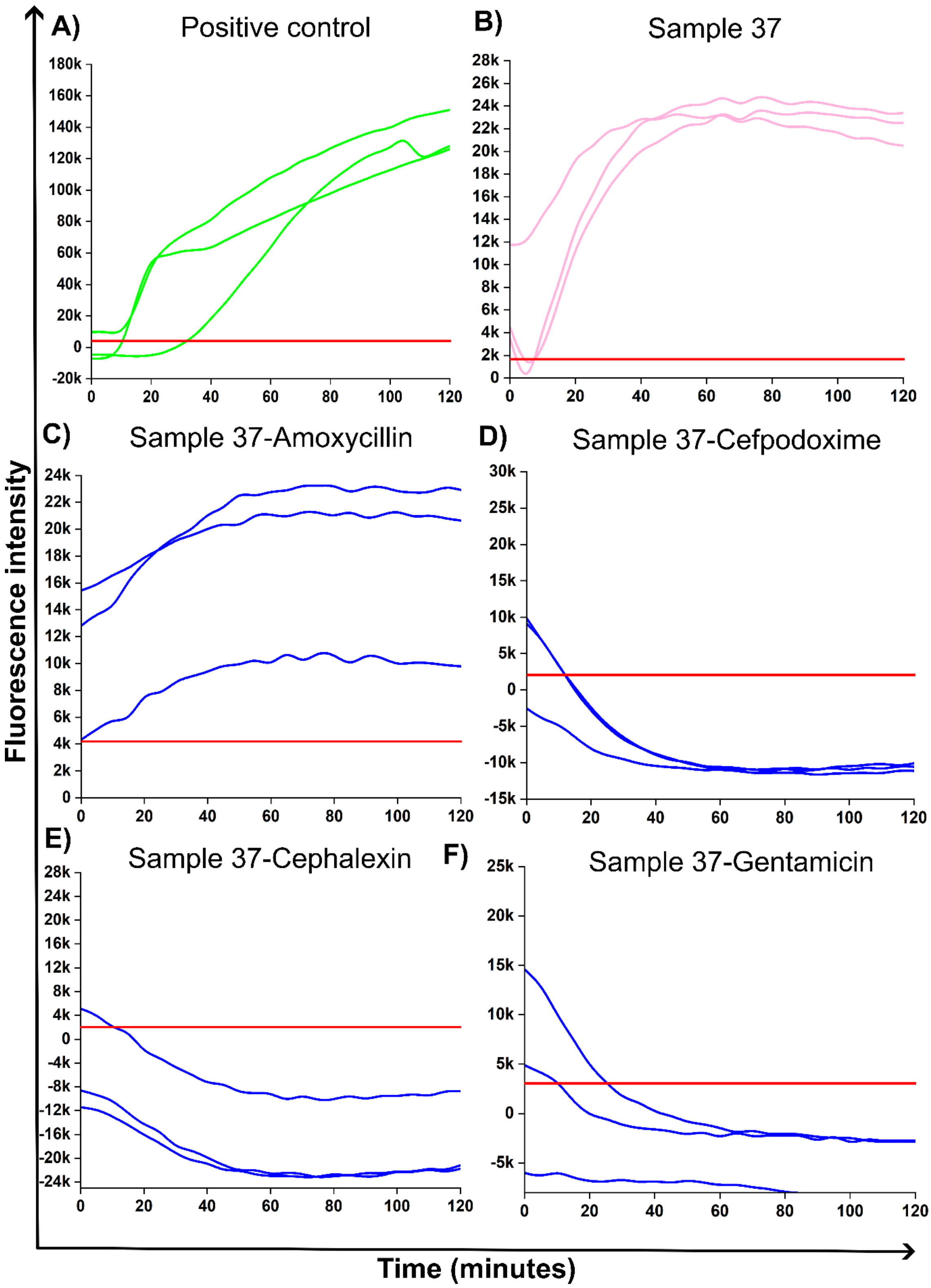


**Figure S29.** **A)** Positive control, **B)** Sample 37 = UTI, **C)** Sample 37 = Amoxicillin resistance, **D)** Sample 37 = Cefpodoxime sensitivity, **E)** Cephalexin sensitivity, and **F)** Sample 37 = Gentamicin sensitivity.

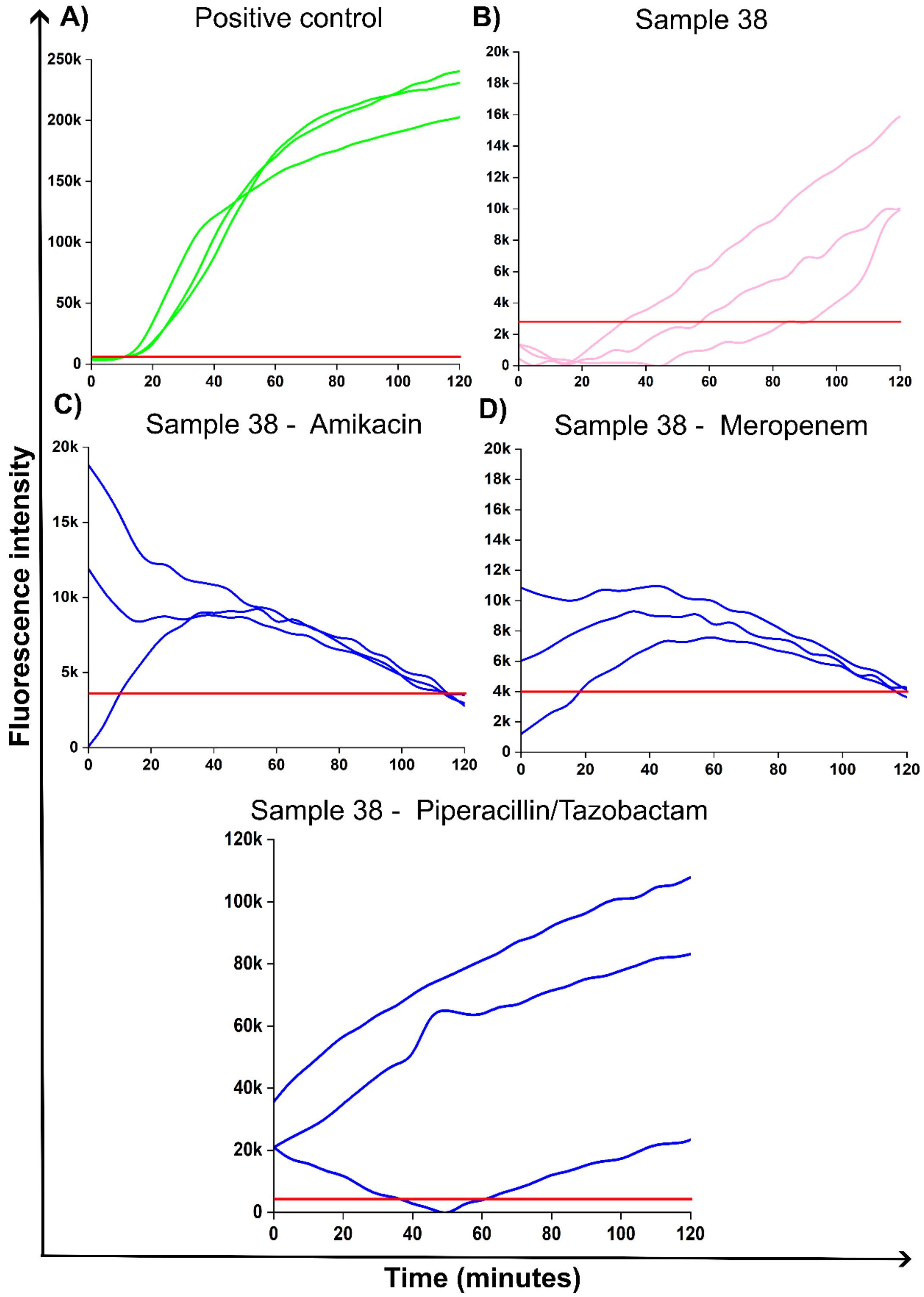


**Figure S30. A)** Positive control, **B)** Sample 38 = UTI, C) Sample 38 = Amikacin sensitivity, D) Sample 38 = Meropenem sensitivity, and E) Sample 38 = Piperacillin/Tazobactam resistance.

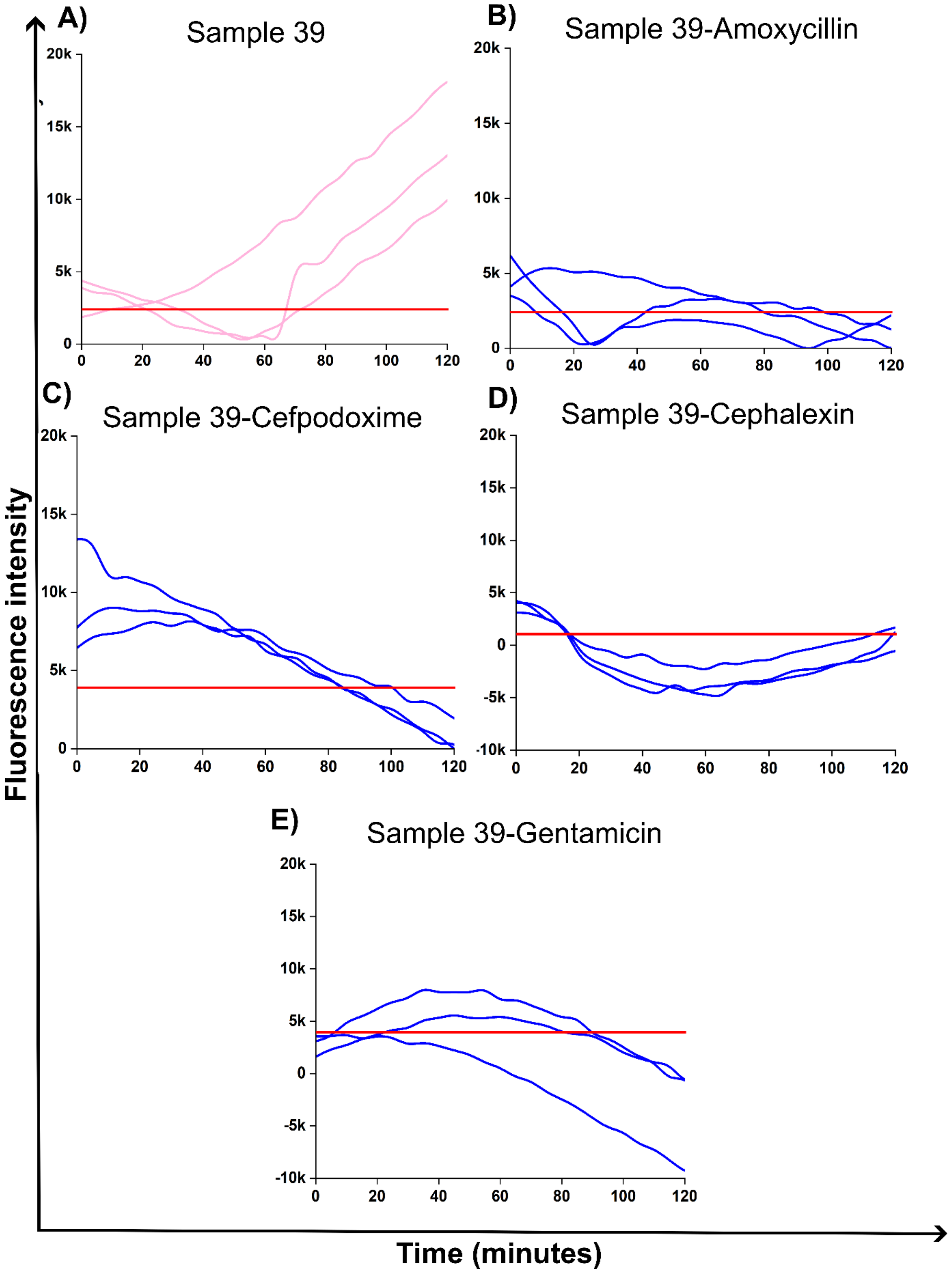


**Figure S31.** **A)** Sample 39 = UTI, **B)** Sample 39 = Amoxicillin sensitivity, **C)** Sample 39 = Cefpodoxime sensitivity, **D)** Sample 39 = Cephalexin sensitivity, and **E)** Sample 39 = Gentamicin sensitivity.

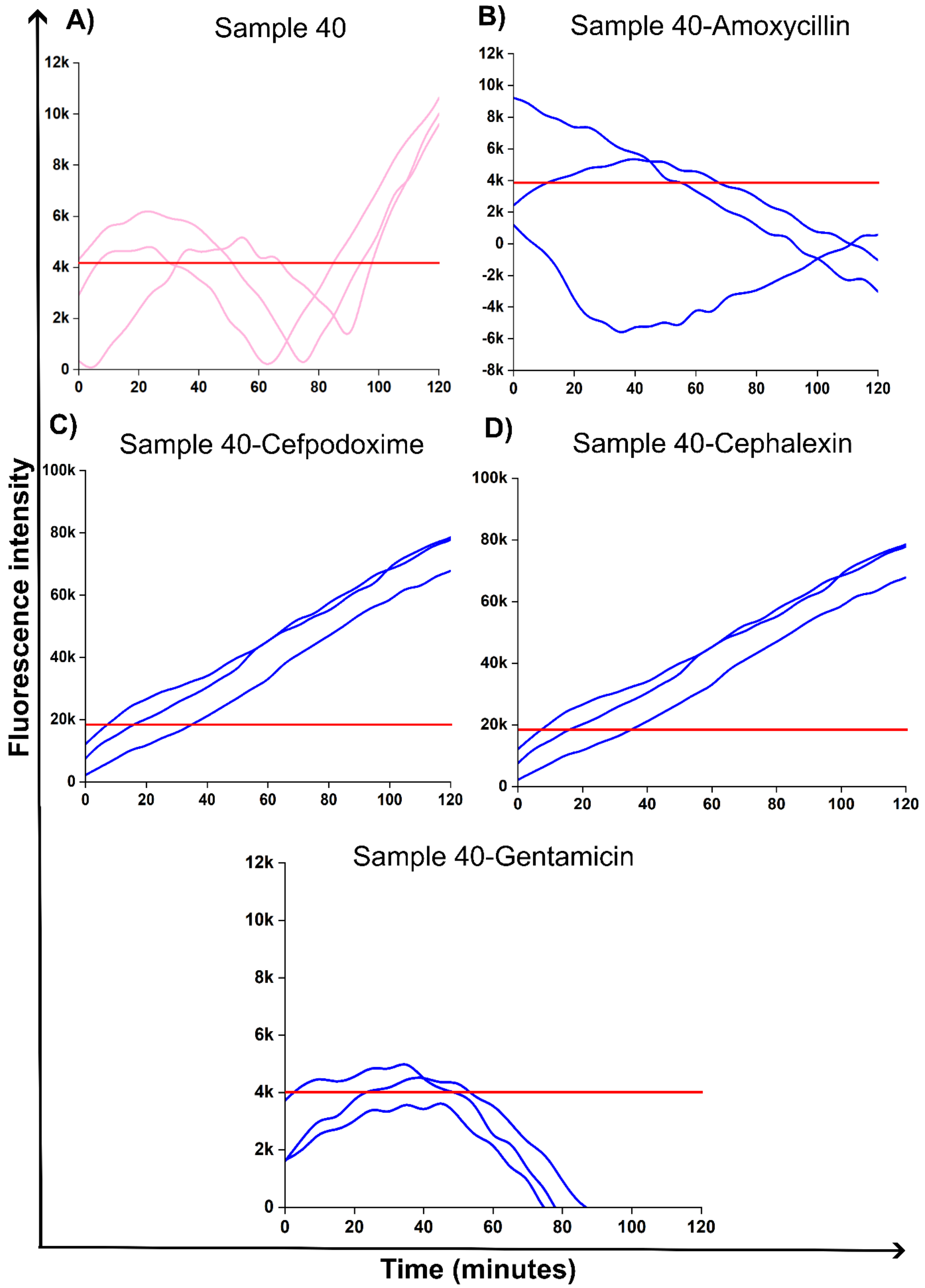


**Figure S32.** **A)** Sample 40 = UTI, **B)** Sample 40 = Amoxicillin sensitivity, **C)** Sample 40 = Cefpodoxime resistance, **D)** Sample 40 = Cephalexin resistance, and **E)** Sample 40 = Gentamicin sensitivity.

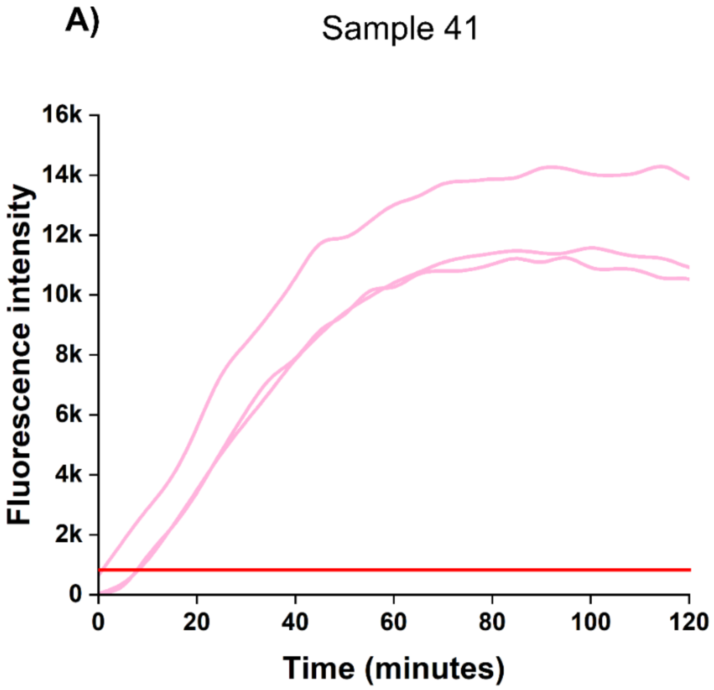


**Figure S33.** **A)** Sample 41 = UTI.

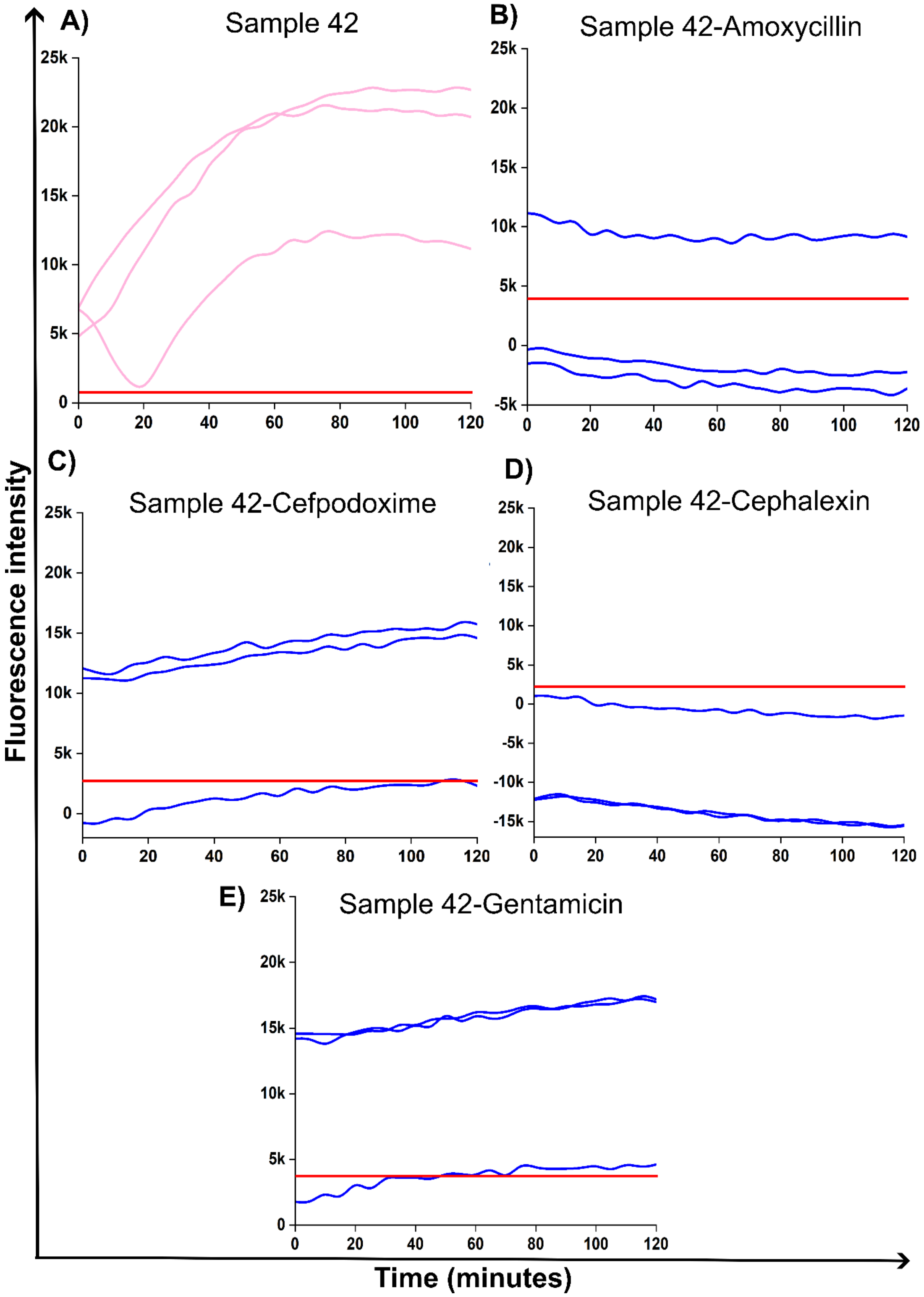


**Figure S34.** **A)** Sample 42 = UTI, **B)** Sample 42 = Amoxicillin sensitivity, **C)** Sample 43 = Cefpodoxime sensitivity, **D)** Sample 43 = Cephalexin sensitivity, and **E)** Gentamicin sensitivity.

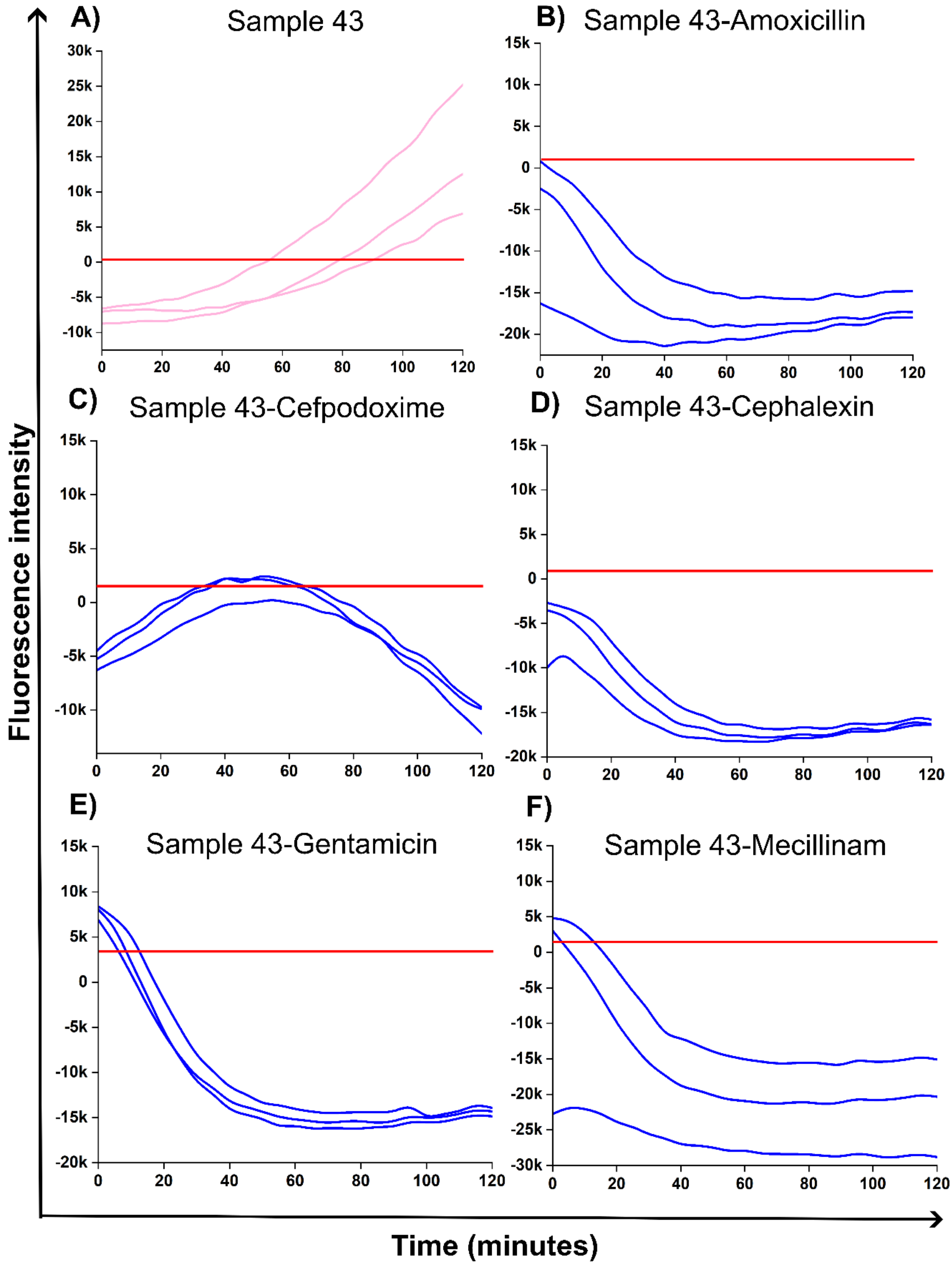


**Figure S35.** **A)** Sample 43 = UTI, **B)** Sample 43 = Amoxicillin sensitivity, **C)** Sample 43 = Cefpodoxime sensitivity, **D)** Sample 43 = Cephalexin sensitivity, **E)** Gentamicin sensitivity, and **F)** Mecillinam sensitivity.

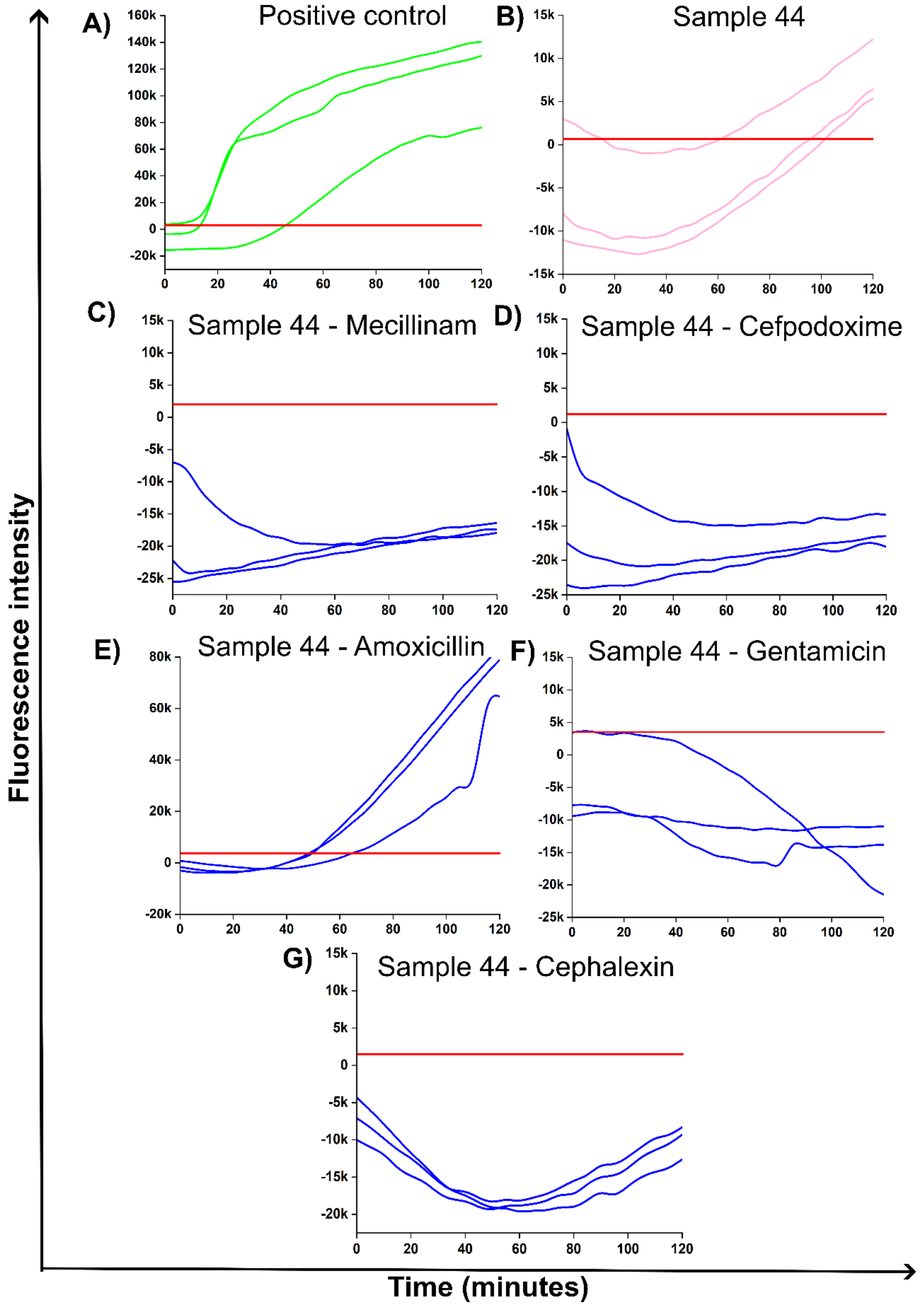


**Figure S36. A)** Positive control, **B)** Sample 44 = UTI, **C)** Sample 44 = Mecillinam sensitivity, **D)** Sample 44 = Cefpodoxime sensitivity, **E)** Sample 44 = Amoxicillin resistance, **F)** Sample 44 = Gentamicin sensitivity, and **G)** Sample 44 = Cephalexin sensitivity.

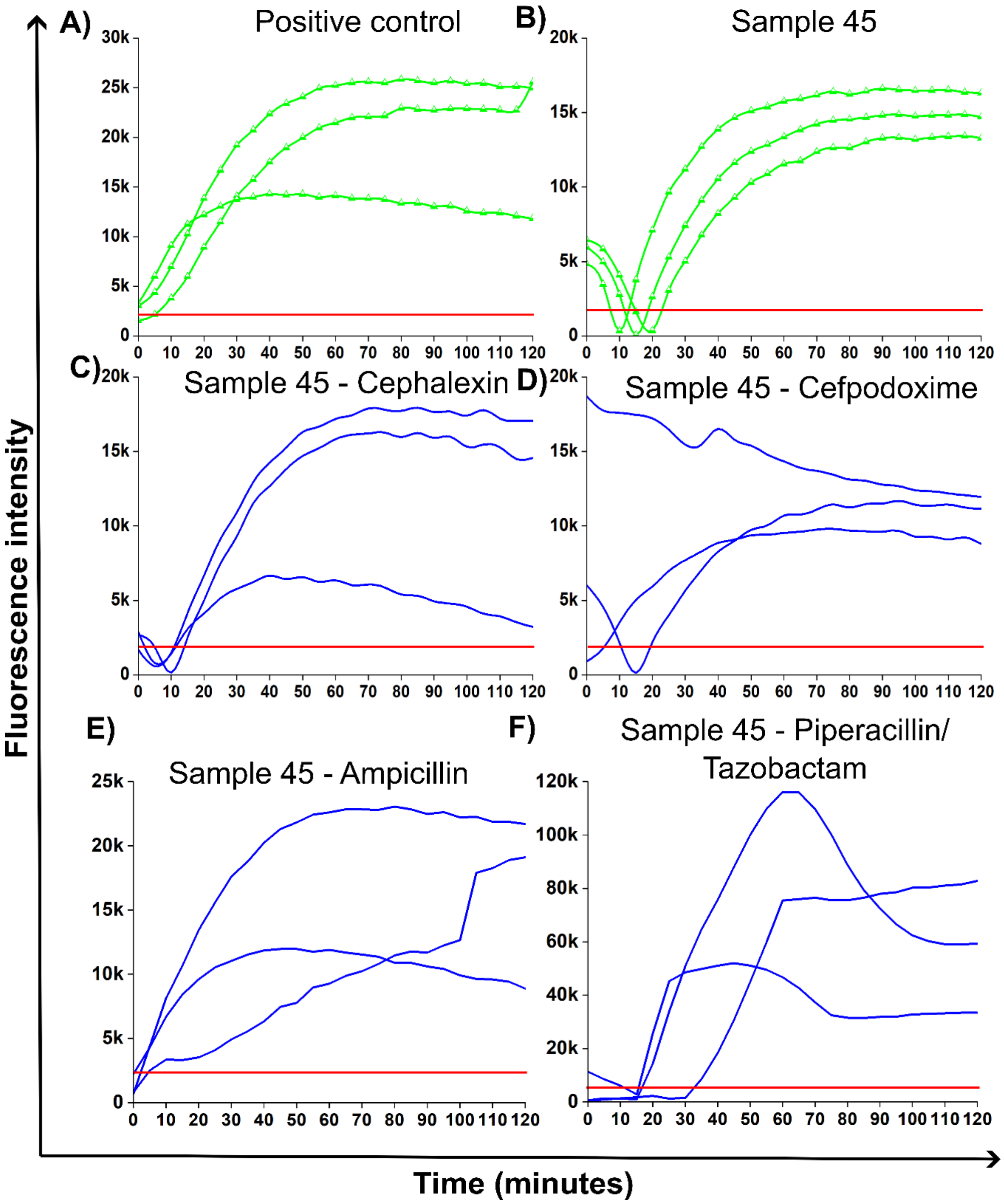

A)
Positive control
B)
Sample 45
C)
Sample 45 - Cephalexin
D)
Sample 45 - Cefpodoxime
E)
Sample 45 - Ampicillin
F)
Sample 45 - Piperacillin/ Tazobactam
Fluorescence intensity
Time (minutes)

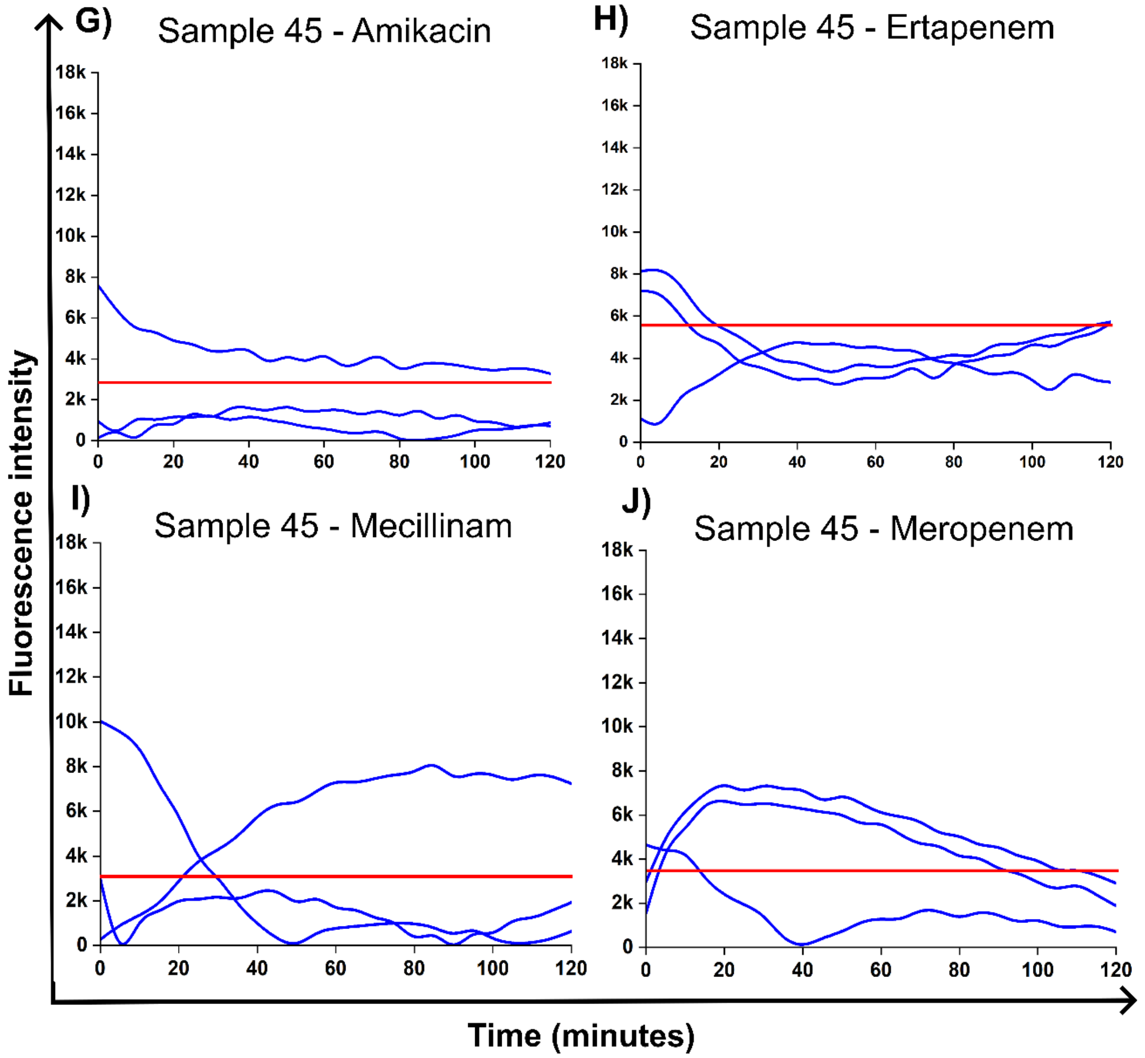


**Figure S37.** **A)** Positive control, **B)** Sample 45 = UTI, **C)** Sample 45 = Cephalexin resistance, **D)** Sample 45 = Cefpodoxime resistance, **E)** Sample 45 = Ampicillin resistance, **F)** Sample 45 = Piperacillin/Tazobactam resistance, and **G)** Sample 45 = Amikacin sensitivity, **H)** Sample 45 = Ertapenem sensitivity, **I)** Sample 45 = Mecillinam sensitivity, and **J)** Sample 45 = Meropenem sensitivity.

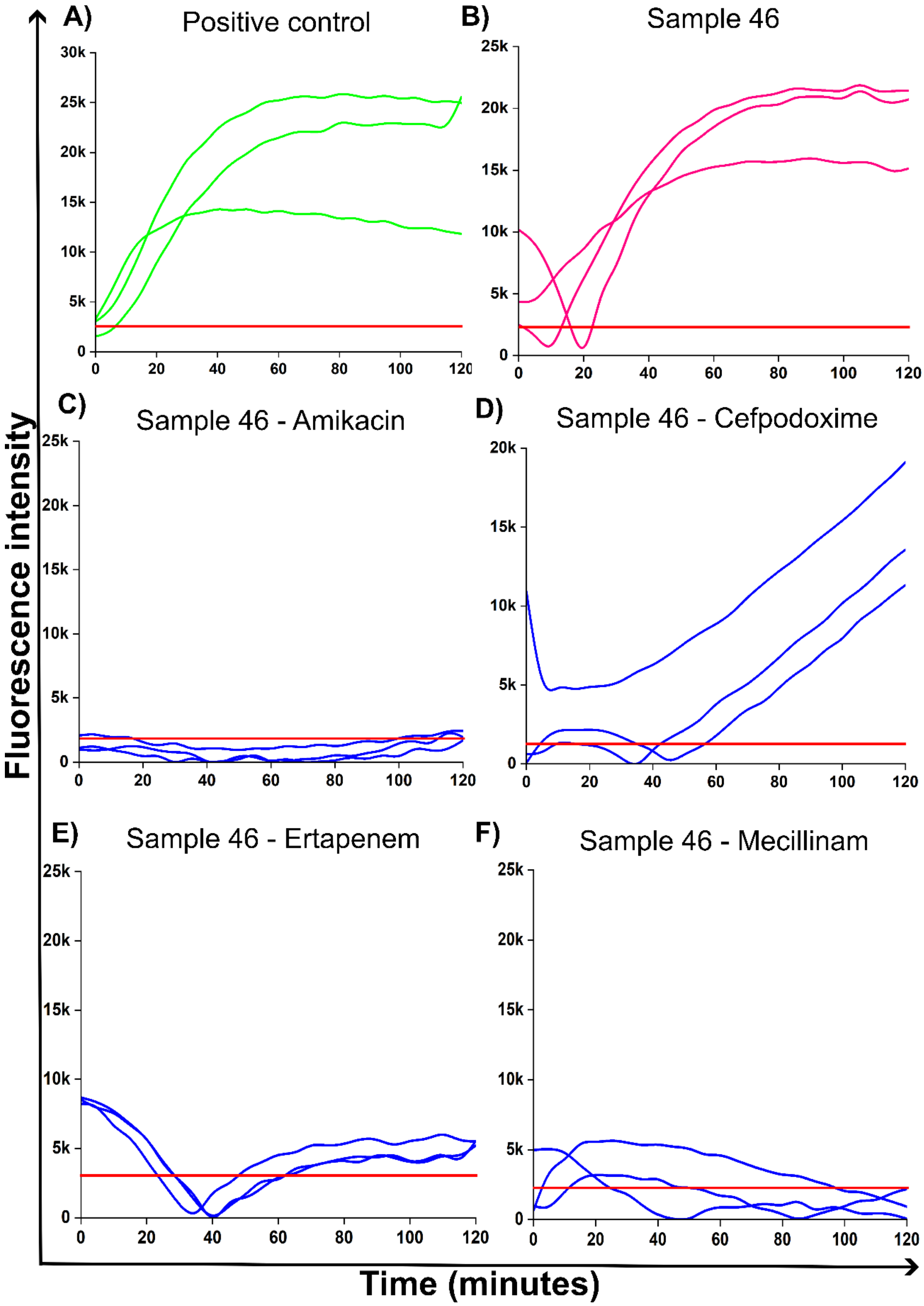

A)
Positive control
30k
25k
20k
15k
10k
5k
0
0
20
40
60
80
100
120
B)
Sample 46
25k
20k
15k
10k
5k
0
0
20
40
60
80
100
120
C)
Sample 46 - Amikacin
D)
Sample 46 - Cefpodoxime
E)
Sample 46 - Ertapenem
F)
Sample 46 - Mecillinam
Fluorescence intensity
Time (minutes)

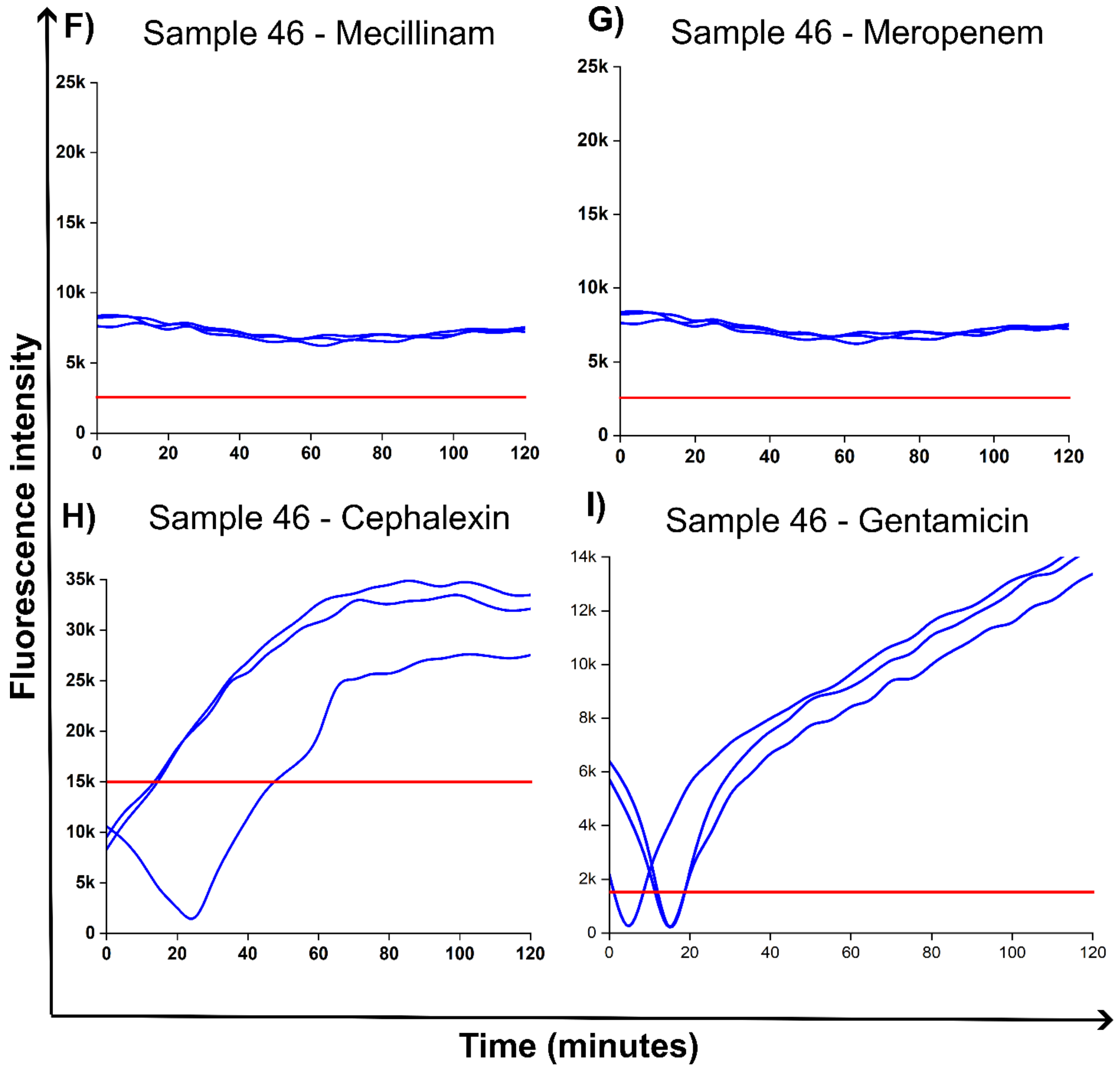


**Figure S38.** **A)** Positive control, **B)** Sample 46 = UTI, **C)** Sample 46 = Amikacin sensitivity, **D)** Sample 46 = Mecillinam resistance, **E)** Sample 46 = Ertapenem sensitivity, **F)** Sample 46 = Mecillinam sensitivity, **G)** Sample 46 = Meropenem sensitivity, **H)** Sample 46 = Cephalexin sensitivity, and **I)** Gentamicin sensitivity.

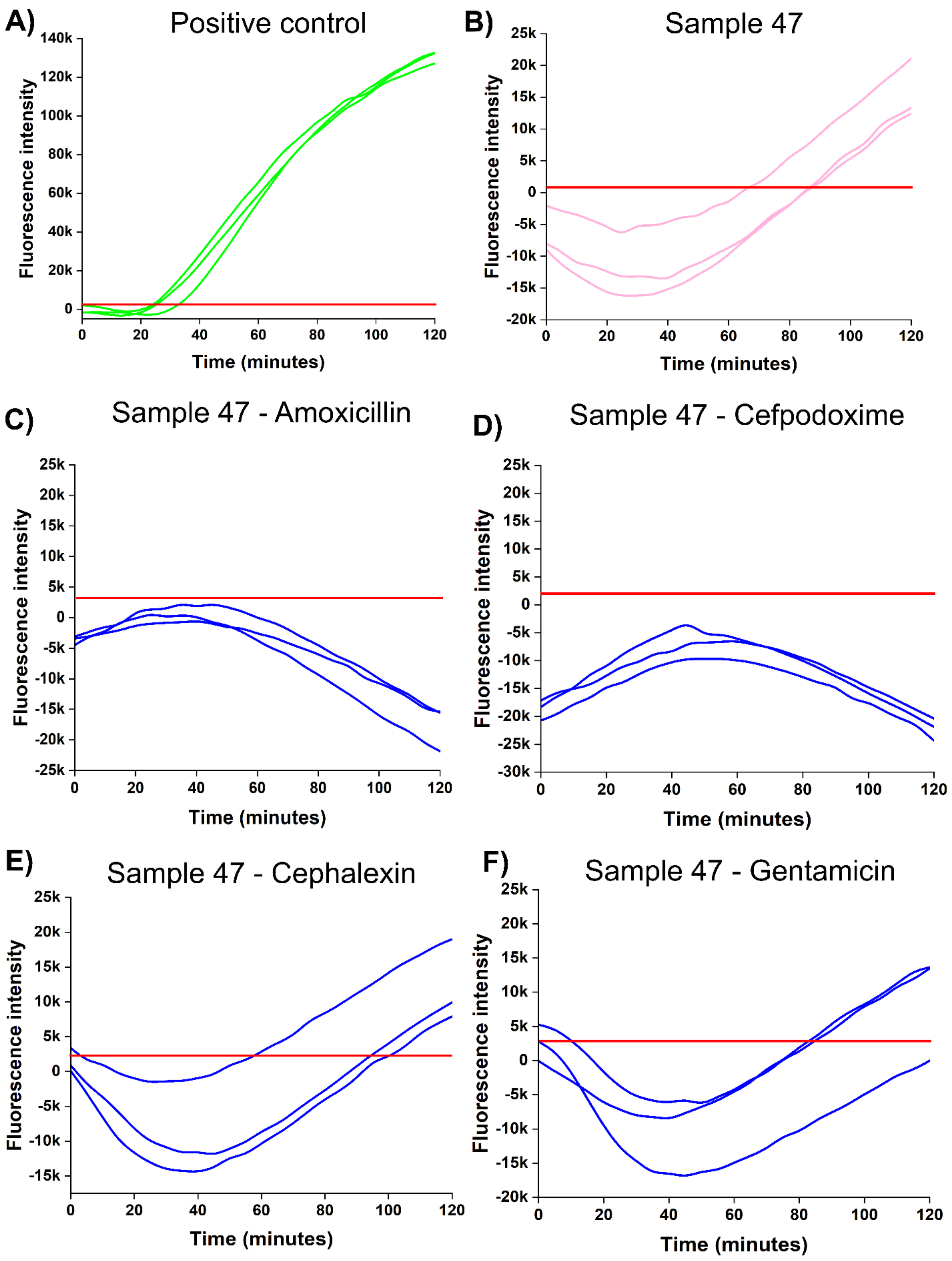


**Figure S39. A)** Positive control, **B)** Sample 47 = UTI, **C)** Sample 47 = Amoxicillin sensitivity, **D)** Sample 47 = Cefpodoxime sensitivity, **E)** Sample 47 = Cephalexin resistance, and **F)** Sample 47 = Gentamicin resistance (False-positive).

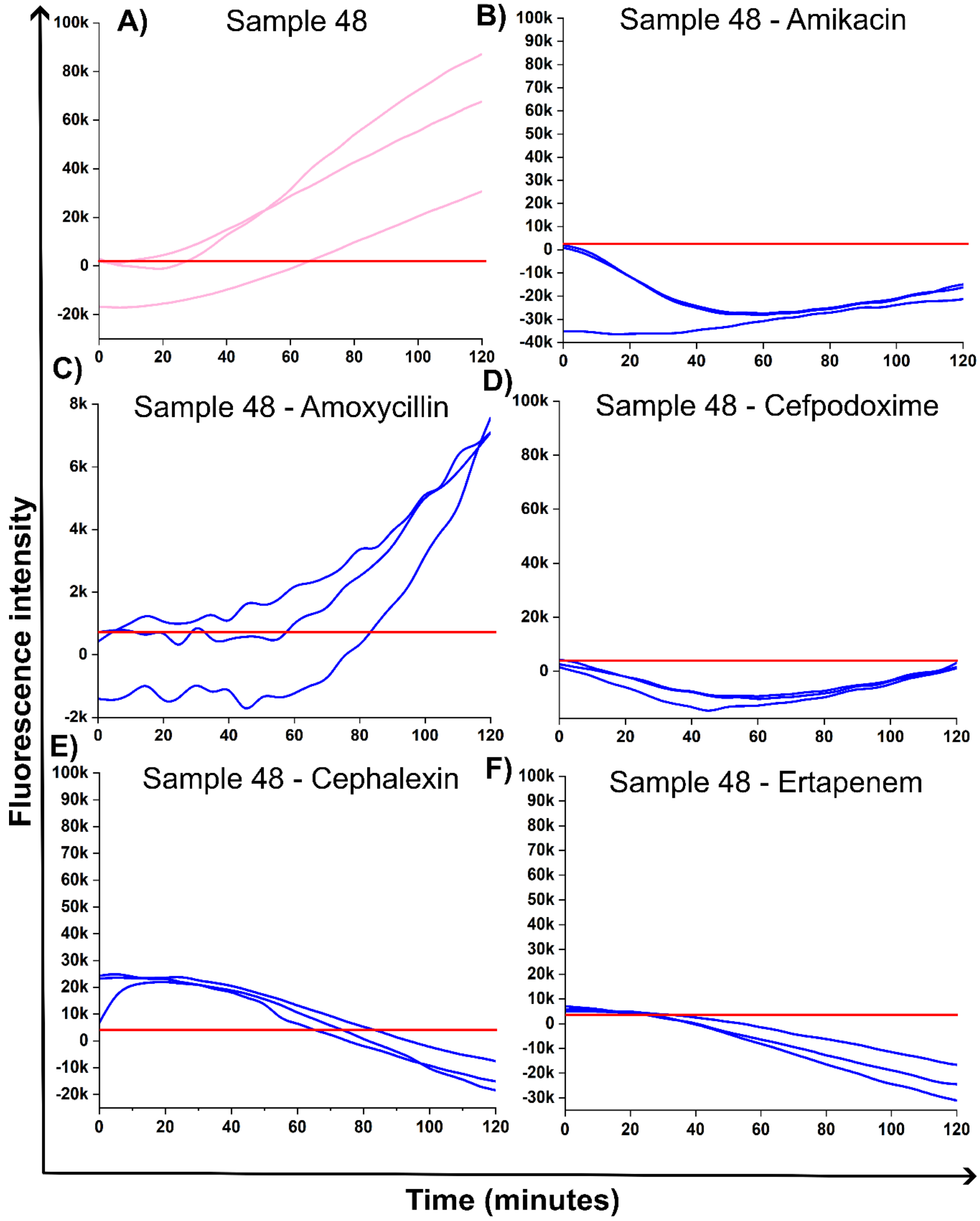
A) Sample 48
B) Sample 48 - Amikacin
C) Sample 48 - Amoxycillin
D) Sample 48 - Cefpodoxime
E) Sample 48 - Cephalexin
F) Sample 48 - Ertapenem
Fluorescence intensity
Time (minutes)
100k
80k
60k
40k
20k
0
-20k
0
20
40
60
80
100
120
90k
70k
50k
30k
10k
-10k
-30k
-40k
8k
6k
4k
2k
-2k

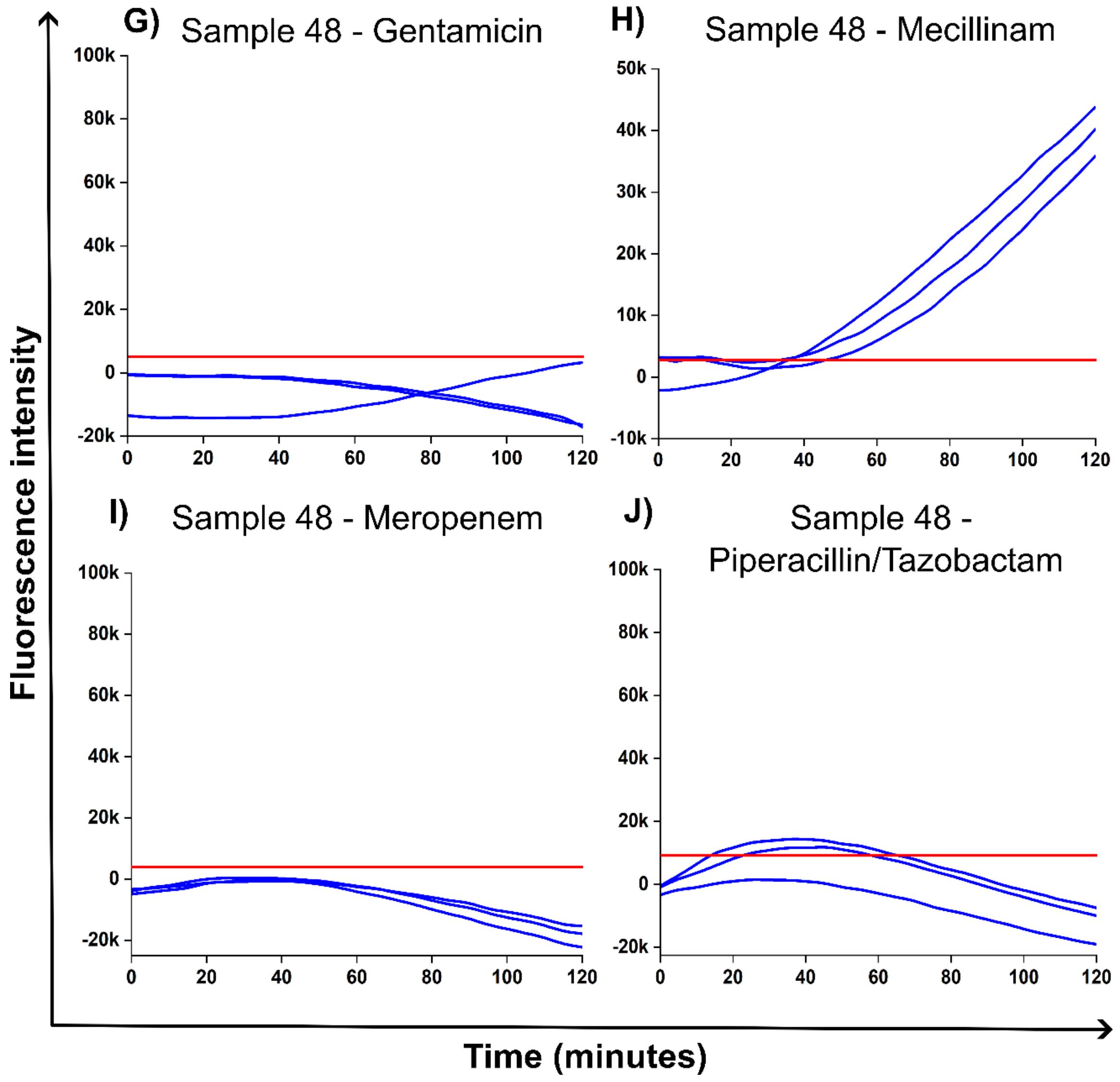


**Figure S40. A)** Sample 48 = UTI, **B)** Sample 48 = Amikacin sensitivity, **C)** Sample 48 = Amoxicillin resistance, **D)** Sample 48 = Cefpodoxime sensitivity, **E)** Sample 48 = Cephalexin sensitivity, **F)** Sample 48 = Ertapenem sensitivity, **G)** Sample 48 = Gentamicin sensitivity, **H)** Sample 48 = Mecillinam resistance, **I)** Sample 48 = Meropenem sensitivity, and **J)** Sample 48 = Piperacillin/Tazobactam.

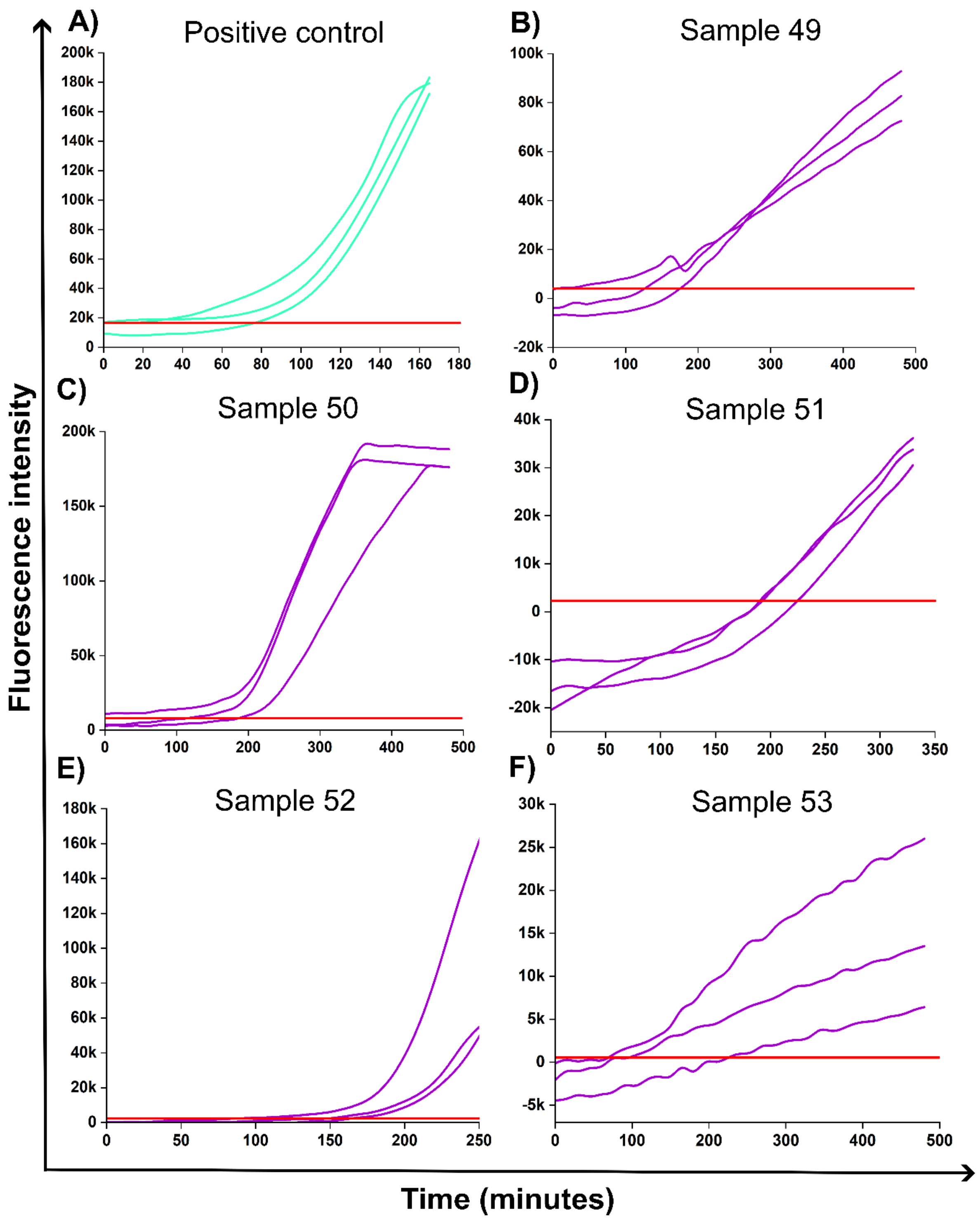


**Figure S41. A)** Positive control. Sample 49 **(B)**, Sample 50 **(C)**, Sample 51 **(D)**, Sample 52 **(E)**, and Sample 53 **(F)** = UTI.

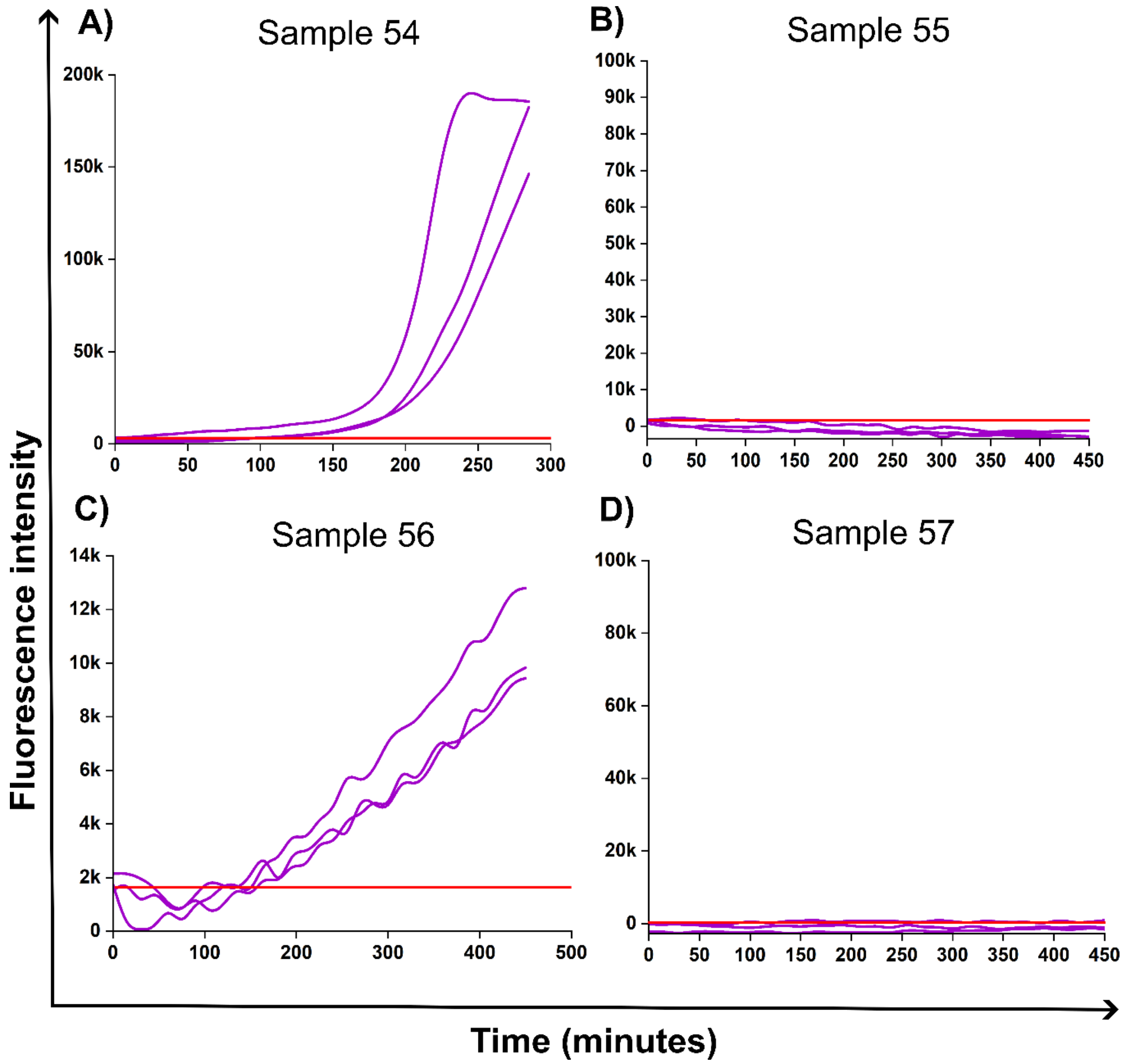


**Figure S42. A)** Positive control. Sample 49 **(B)**, Sample 50 **(C)**, Sample 51 **(D)**, Sample 52 **(E)**, and Sample 53 **(F)** = UTI.

| Clinical samples | UTI-AMR status/ Uropathogenic bacteria | Resistance Or Intermediate resistance (*I) | Sensitivity |
|---|---|---|---|
| **Sample 1** | No UTI | - | - |
| **Sample 2** | *E. coli* | - | - |
| **Sample 3** | No UTI | - | - |
| **Sample 8** | No UTI | - | |
| **Sample 9** | No UTI | - | - |

| | | | |
|---|---|---|---|
| **Sample 10** | No UTI | - | - |
| **Sample 11** | No UTI | - | - |
| **Sample 12** | *P. mirabilis* | - | Amoxicillin<br>Cephalexin<br>Gentamicin<br>Mecillinam<br>Cefpodoxime |
| **Sample 13** | *Klebsiella oxytoca* | Amoxicillin | Cephalexin<br>Gentamicin<br>Mecillinam<br>Cefpodoxime |
| **Sample 14** | *Klebsiella variicola* | - | Cefpodoxime |
| **Sample 15** | Mixed growth | - | - |
| **Sample 16** | Mixed growth | - | - |
| **Sample 17** | Mixed growth | - | - |
| **Sample 18** | Mixed growth | - | - |
| **Sample 19** | Mixed growth | - | - |
| **Sample 20** | *E. coli* | | Amoxicillin<br>Cephalexin<br>Gentamicin<br>Mecillinam<br>Cefpodoxime |
| **Sample 21** | *E. coli* | Amoxicillin | Cephalexin<br>Gentamicin<br>Mecillinam<br>Cefpodoxime |
| **Sample 22** | *E. coli* | Amoxicillin | Cephalexin<br>Gentamicin<br>Mecillinam<br>Cefpodoxime |
| **Sample 23** | *E. coli* | | Amoxicillin<br>Cephalexin<br>Gentamicin<br>Mecillinam<br>Cefpodoxime |

| | | | |
|---|---|---|---|
| **Sample 24** | *E. coli* | | Amoxicillin<br>Cephalexin<br>Gentamicin<br>Mecillinam<br>Cefpodoxime |
| **Sample 25** | *ESBL E. coli* | Amoxicillin<br>Cephalexin<br>Mecillinam<br>Cefpodoxime | Gentamicin |
| **Sample 26** | *S. simulans* | | Amoxicillin<br>Cephalexin<br>Gentamicin<br>Cefpodoxime |
| **Sample 27** | *Enterococcus sp.* | Amoxicillin | Cephalexin |
| **Sample 28** | *Enterococcus faecium* | Amoxicillin<br>Cephalexin | |
| **Sample 33** | *P. mirabilis* | | Gentamicin<br>Cephalexin<br>Cefpodoxime<br>Amoxicillin |
| **Sample 35** | *Step. Group B.* | Amoxicillin | Cephalexin<br>Cefpodoxime<br>Gentamicin<br>Mecillinam |
| **Sample 36** | *K. pneumoniae* | Amoxicillin | Cephalexin<br>Gentamicin<br>Mecillinam<br>Cefpodoxime |
| **Sample 37** | *K. variicola* | Amoxicillin | Cephalexin<br>Gentamicin<br>Cefpodoxime |
| **Sample 38** | *P. aeruginosa* | Piperacillin/Tazobactam | Amikacin<br>Meropenem |
| **Sample 39** | *S. simulans* | | Amoxicillin<br>Cephalexin |

| | | | |
|---|---|---|---|
| | | | Gentamicin<br>Cefpodoxime |
| **Sample 40** | *S. saprophyticus* | Cefpodoxime | Amoxicillin<br>Cephalexin<br>Gentamicin |
| **Sample 41** | *S. epidermidis* | - | - |
| **Sample 42** | *E. coli* | - | Amoxicillin<br>Cephalexin<br>Gentamicin<br>Cefpodoxime |
| **Sample 43** | *E. coli* | - | Amoxicillin<br>Cephalexin<br>Gentamicin<br>Mecillinam<br>Cefpodoxime |
| **Sample 44** | *E. coli* | Amoxicillin | Cephalexin<br>Gentamicin<br>Mecillinam<br>Cefpodoxime |
| **Sample 45** | *ESBL E. coli* | Amoxicillin<br>Cephalexin<br>Piperacillin/Tazobactam<br>Cefpodoxime | Amikacin<br>Ertapenem<br>Mecillinam<br>Meropenem |
| **Sample 46** | *ESBL E. coli* | Cephalexin<br>Gentamicin<br>Cefpodoxime | Amikacin<br>Ertapenem<br>Mecillinam<br>Meropenem |
| **Sample 47** | *E. coli* | - | Amoxicillin<br>Cefpodoxime<br>Gentamicin<br>Cephalexin |
| **Sample 48** | *E. coli* | Amoxicillin<br>Mecillinam | Amikacin<br>Cefpodoxime<br>Cephalexin<br>Ertapenem<br>Gentamicin |

| | | | |
|---|---|---|---|
| | | | Meropenem<br>Piperacillin/Tazobactam |
| **Sample 49** | *E. coli* | - | - |
| **Sample 50** | *ESBL E. coli* | - | - |
| **Sample 51** | *E. coli* | - | - |
| **Sample 52** | *Klebsiella* species | - | - |
| **Sample 53** | *Proteus mirabilis* | - | - |
| **Sample 54** | *Pseudomonas aeruginosa* | - | - |
| **Sample 55** | *E. coli* | - | - |
| **Sample 56** | Yeast | | |
| **Sample 57** | *E. coli* | - | - |

**Table S5.** Clinical gold standard results as confirmed by urine culture

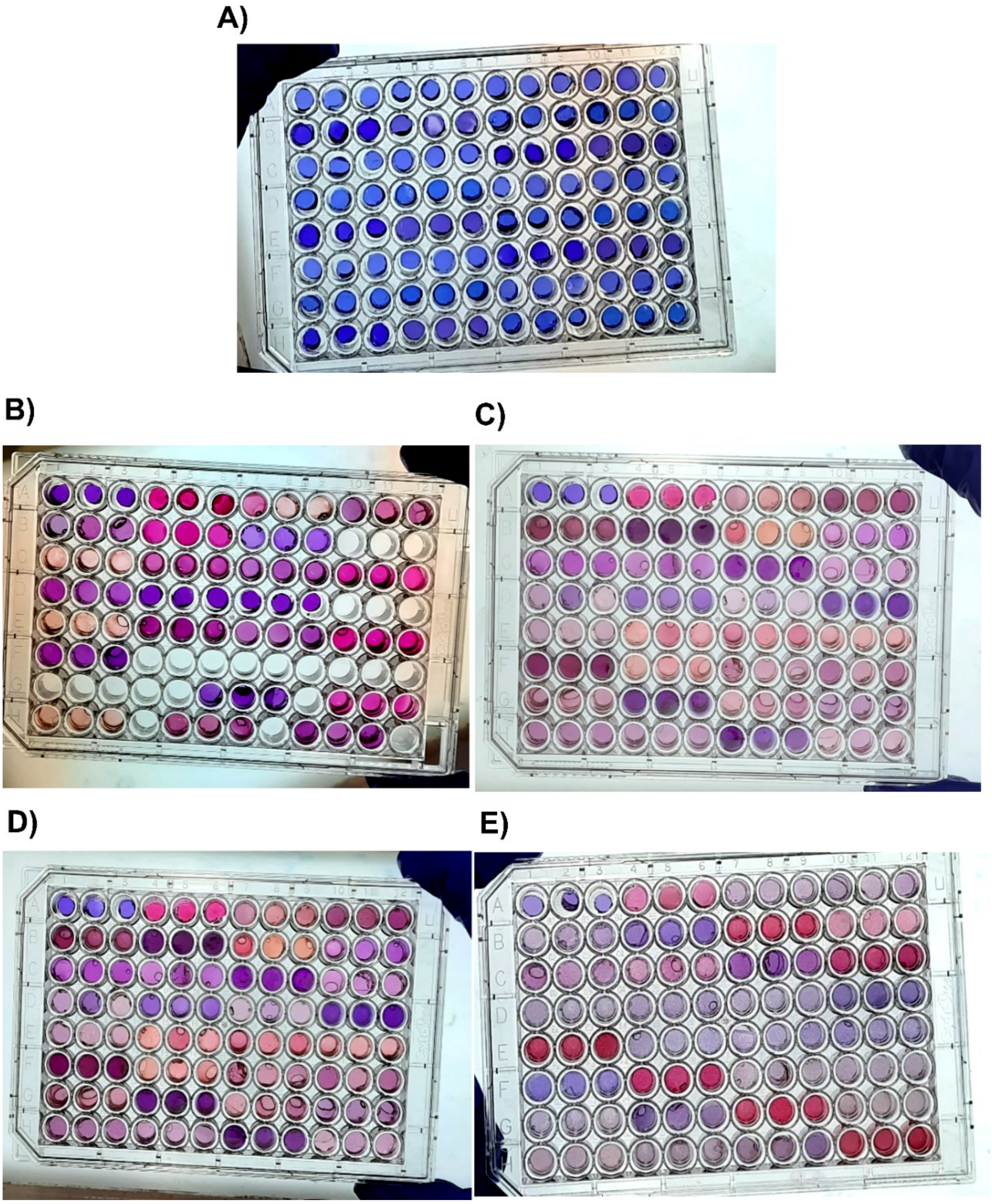


**Figure S43.** Images of 96-well plates containing multifunctional hydrogels before (A) and after (B)-(E) UTI-AMR testing.

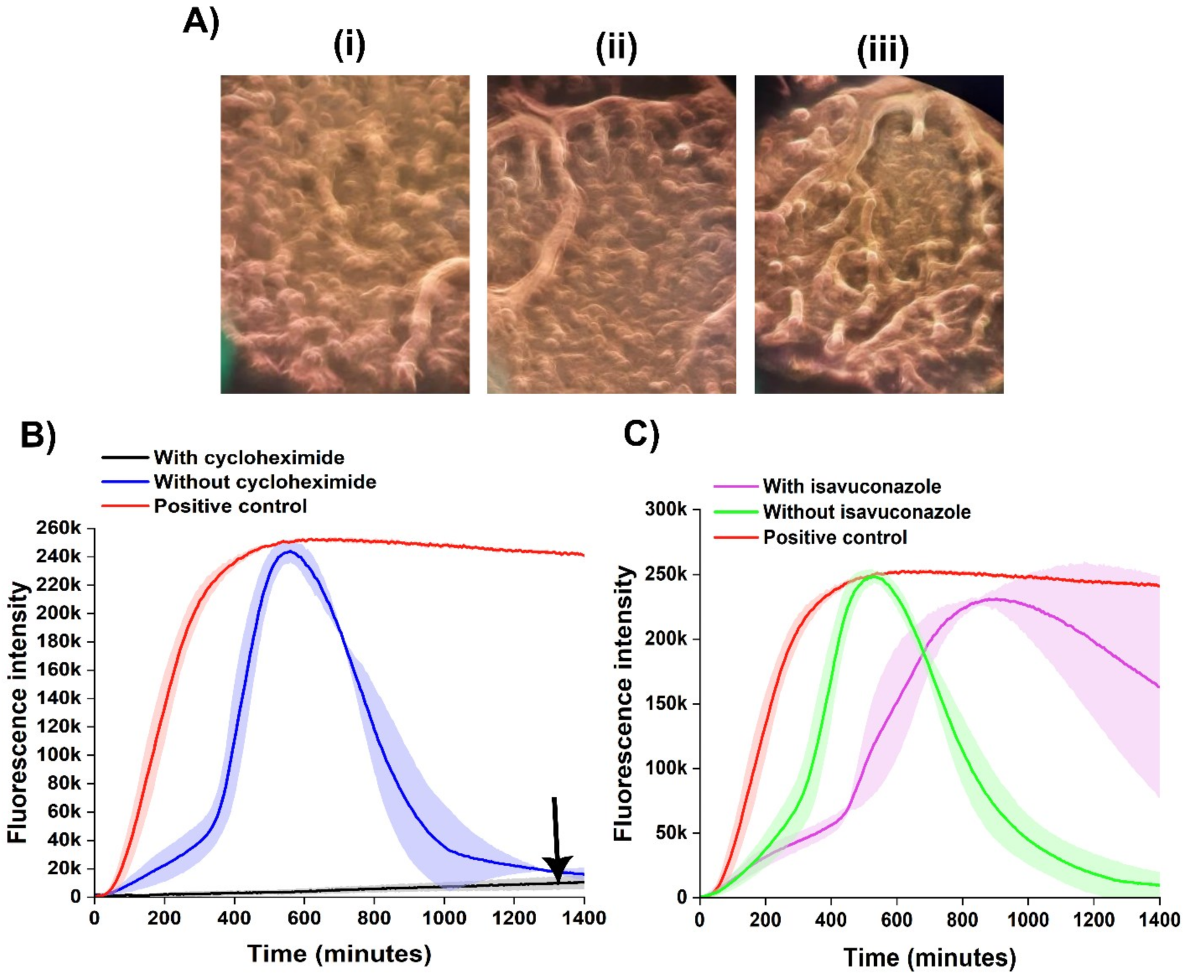


**Figure S44.** Characterization of fungal metabolic kinetics and targeted antimycotic inhibition within NaAMPS hydrogels. **(A)** i-iii. Microscopic observation of progressive fungal vegetative growth and colony networks on the hydrogel surface. **(B)** Real-time flurometric tracking over 1400 minutes comparing uninhibited fungal metabolism (blue line) and positive controls (red line) against matrices containing the proteins synthesis inhibitor cycloheximide (CHX 100 µg/mL, black line). The flat kinetic profile under cycloheximide exposure confirms successful metabolilc suppression. **(C)** Corresponding real-time fluorometric curves demonstrating a lack of fungal inhibition under an initial concentration (100 µg/mL) of the clinical triazole antifungal isavuconazole (purple line), which exhibits a rapid unsuppressed trajectory closely matching the uninhibited positive control (red line) and baseline fungal growth (green line). Shaded regions signify the standard deviation across independent replicates (n=3).

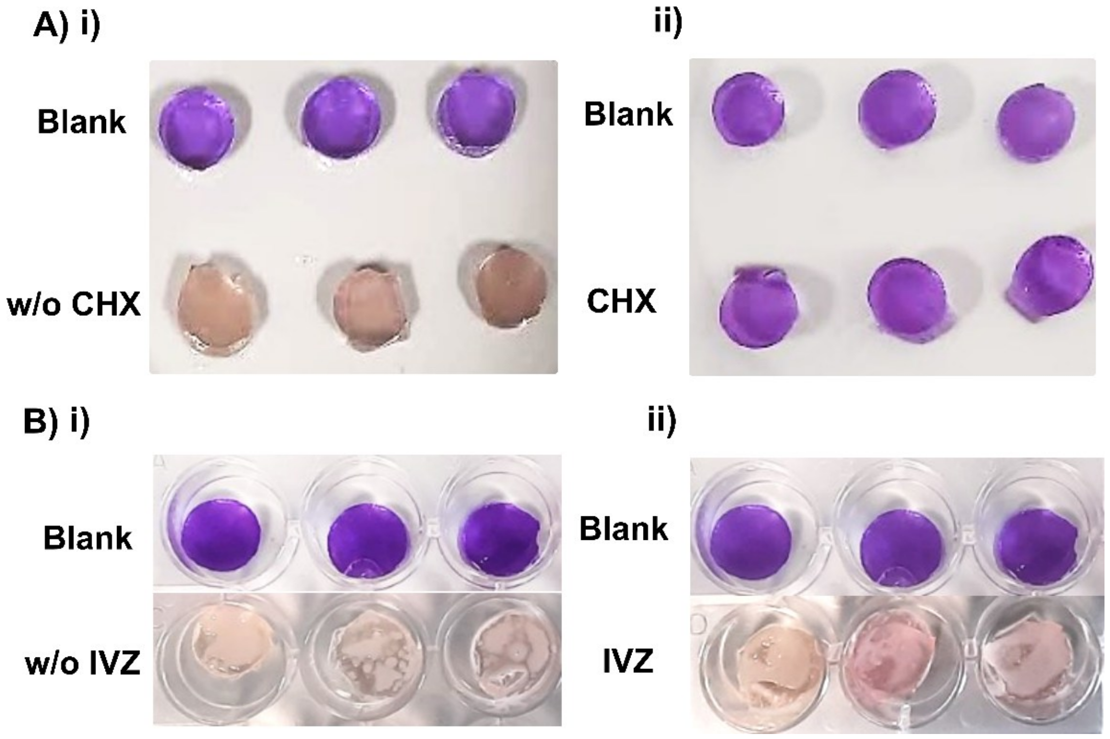


**Figure S45.** Colorimetric validation of fungal background elimination. Photographic profiles of multi-well hydrogel assays assessing metabolic cross-reactivity and verification of fungal growth containment. **(A)** uninoculated blank control hydrogels maintaing the standard purple baseline compared against fungal-inoculated matrices without cycloheximide (w/o CHX, left (**i**)), which transition to a clear, reduced cream-bream colour, and hydrogels containing cycloheximide (CHX 100 µg/mL, right (**ii**)), which remain unreduced and purple. **(B)** Corresponding colorimetric evaluations with isavuconazole (IVZ), showing clear naked-eye visual discrimination between uninhibitated fungal reduction without IVZ (left (**i**)) and with 100 µg/mL IVZ (right (**ii**)).

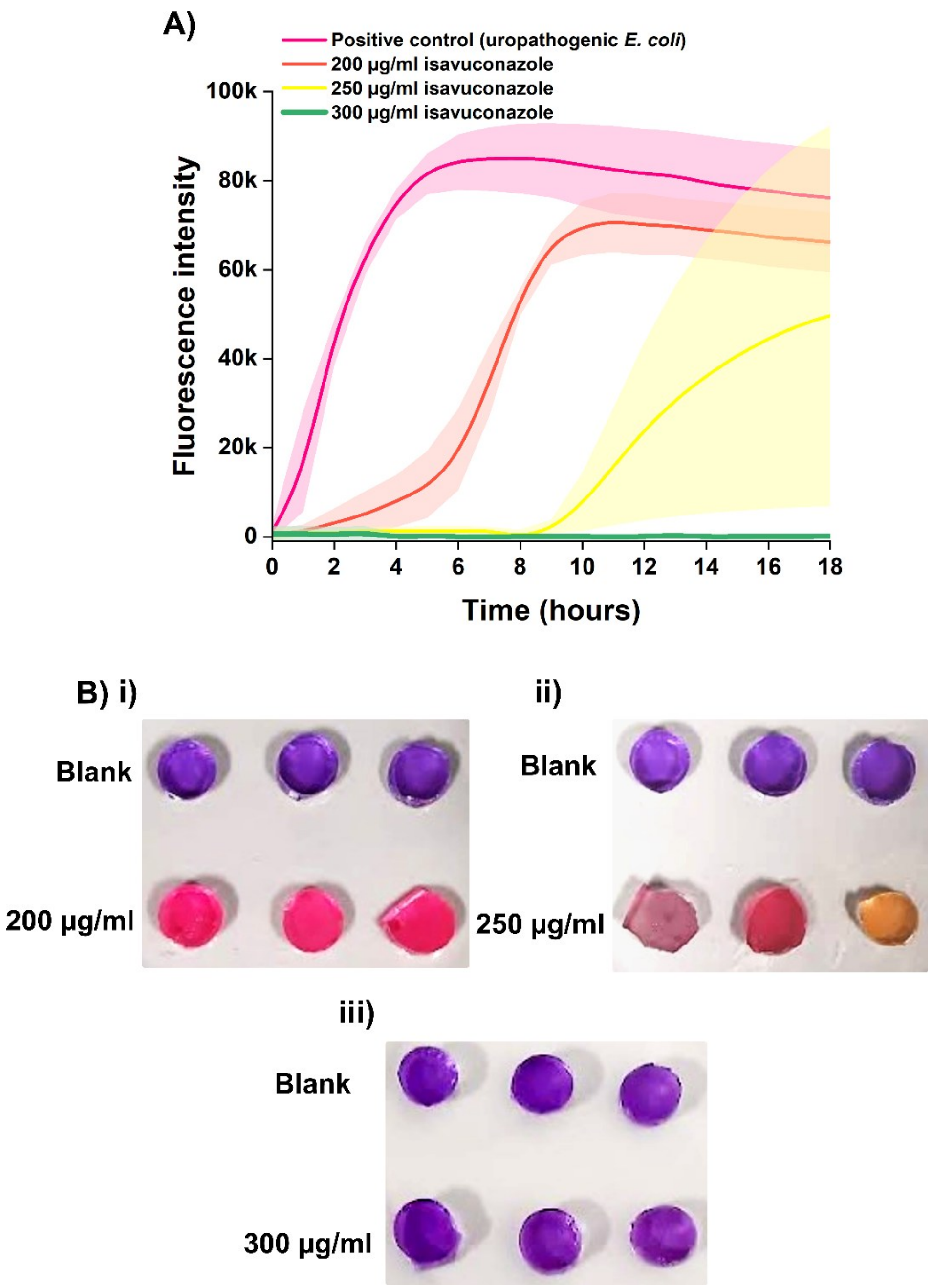


**Figure S46.** IVZ dose-response profiles confirming suppression of fungal interference. **(A)** Real-time fluorometric trajectories of clinical fungal isolates exposed to an elevated concentration gradient of isavuconazole (200, 250, and 300 µg/mL) compared against an uninhibited uropathogenic E. coli positive control (pink line). Shaded regions represent the standard deviation (n=3). The platform captures a clear, concentration-dependent kinetic delay, resulting in a progressive rightward shift of the exponential curves as the antifungal concentration approaches the effective inhibitory threshold. **(B) i-iii.** Endpoint colorimetric hydrogel discs. Sub-therapeutic concentrations (**i**, 200 µg/mL and **ii**, 250 µg/mL) permit localized fungal breakout and subsequent resazurin reduction to a pink-cream-yellow colour, whereas the true therapeutic threshold of 300 µg/mL (**iii**) provides absolute metabolic containment, where the hydrogel network is in its unreduced purple state to eliminate background cross-reactivity.

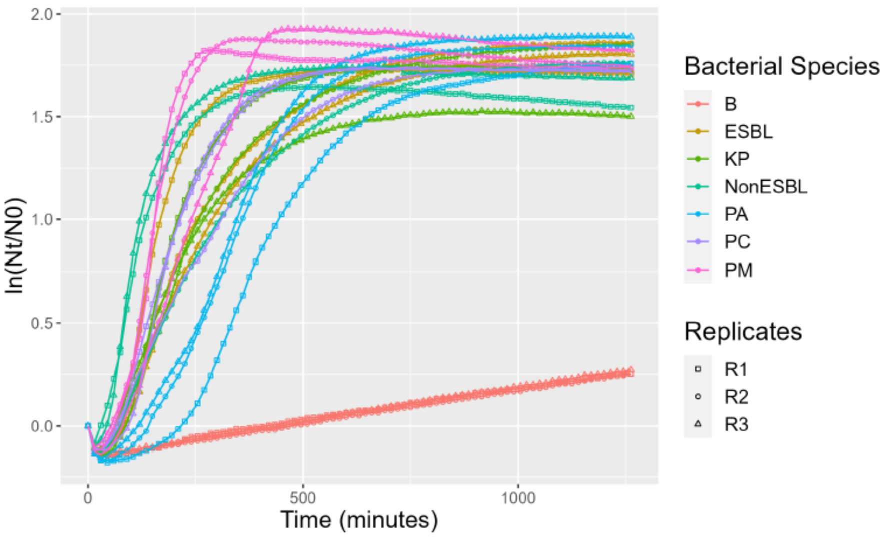


**Figure S47.** Multi-species real-time kinetic growth curve modeling within the hydrogel matrix. Real-time growth profile trajectories tracking the log relative population size ln (Nt/N0) (Y-axis) as a function of Time in minutes (X-axis) across 1250-minute monitoring window. Data series represent distinct clinical uropathogen classes and control conditions across multiple independent replicates (R1, squares; R2, circles; R3, triangles). The distinct, sigmoidal trajectories display high reproductive precision across replicates while captuirng species-specific metabolic variations. Notably, slow-growing or highly distinct metabolic profiles such as Pseudomonas aeruginosa (PA) show delayed metabolic timelines compared to rapid fermenters like Proteus mirabilis (PM), visually highlighting the phenotypic kinetic differences that enable high-accuracy downstream species-level classification.

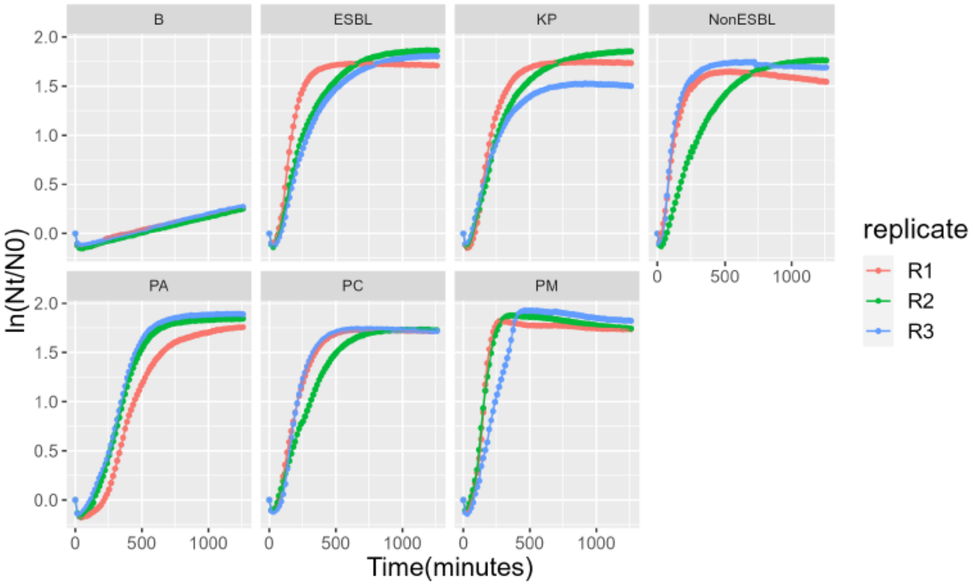


**Figure S48.** Statistical validation and structural fit of the reparameterized Gompertz nonlinear mixed-effects (NLME) model over a 1250-minute timeline. The scatter plot tracks individual standardized modeling residuals over time for a single bacterial species. The narrow, uniform scattering of data points tightly bounding the zero axis across the entire duration of the assay confirms homoscedasticity and demonstrates that the signmoidal framework accurately captures the kinetic profile, validating the high fidelity of the extracted phenotypic growth parameters.

| Model | | R.effect | $A$ | $\mu_m$ | $\lambda$ | AIC | Test | p-value |
|---|---|---|---|---|---|---|---|---|
| Logistic | PCLG1 | $A, \mu_m, \lambda$ | 1.713*** | 0.0068*** | 91.733*** | -652.438 | | |
| | PCLG2 | $A, \mu_m$ | 1.713*** | 0.0067*** | 93.883*** | -656.117 | 1 vs 2 | 0.0215 |
| | PCLG3 | $A, \lambda$ | 1.709*** | 0.0064*** | 95.280*** | -519.603 | 1 vs 3 | <0.0001 |
| | PCLG4 | $\mu_m, \lambda$ | 1.713*** | 0.0068*** | 91.708*** | -662.052 | 1 vs 4 | 0.9957 |
| | PCLG5 | $\mu_m$ | 1.713*** | 0.0067*** | 93.884*** | -656.438 | 4 vs 5 | 0.0082 |
| | PCLG6 | $\lambda$ | 1.710*** | 0.0064*** | 94.949*** | -514.347 | 4 vs 6 | <0.0001 |
| Gompertz | PCGMP1 | $A, \mu_m, \lambda$ | 1.727*** | 0.0068*** | 83.152*** | -928.689 | | |
| | PCGMP2 | $A, \mu_m$ | 1.727*** | 0.0067*** | 85.110*** | -891.016 | 1 vs 2 | <0.0001 |
| | PCGMP3 | $A, \lambda$ | 1.721*** | 0.0065*** | 86.594*** | -619.078 | 1 vs 3 | <0.0001 |
| | PCGMP4 | $\mu_m, \lambda$ | 1.727*** | 0.0068*** | 83.239*** | -934.341 | 1 vs 4 | 0.9506 |
| | PCGMP5 | $\mu_m$ | 1.727*** | 0.0067*** | 85.111*** | -895.016 | 4 vs 5 | <0.0001 |
| | PCGMP6 | $\lambda$ | 1.723*** | 0.0064*** | 86.307*** | -606.342 | 4 vs 6 | <0.0001 |
| Baranyi-Roberts | PCBR1 | $A, \mu_m, \lambda$ | 1.720*** | 0.0147*** | 156.221*** | -765.824 | | |
| | PCBR2 | $A, \mu_m$ | 1.721*** | 0.0146*** | 154.165*** | -759.369 | 1 vs 2 | 0.006 |
| | PCBR3 | $A, \lambda$ | 1.716*** | 0.0134*** | 158.455*** | -602.296 | 1 vs 3 | <0.0001 |
| | PCBR4 | $\mu_m, \lambda$ | 1.720*** | 0.0147*** | 156.191*** | -771.702 | 1 vs 4 | 0.9891 |
| | PCBR5 | $\mu_m$ | 1.721*** | 0.0146*** | 154.165*** | -763.369 | 4 vs 5 | 0.0021 |
| | PCBR6 | $\lambda$ | 1.719*** | 0.0133*** | 157.970*** | -587.415 | 4 vs 6 | <0.0001 |

**Table S6.** Comparative statistical profiling of 18 specific model variations across the Logistic (PCLG1-PCLG6), Gompertz (PCGMP1-PCGMP6), and Baranyi-Roberts (PCBR1-PCBR6) growth classes. The table details the estimated fixed-effect parameters-asymptotic yield plateau (A), maximum specific growth rate ($\mu_m$), and lag phase duration ($\lambda$) alongside their specific random-effect configurations (R.effect) and Akaike Information Criterion (AIC) scores. Nested analysis of variance (ANOVA) likelihood ratio tests assess the statistical significance of step-down random-effect exclusions against saturated baseline models, validating model parsimony and choice based on calculated p-values (***P<0.001).

| Model | df | AIC | dAIC | weight | Mar.$R^2$ | Con. $R^2$ | RMSE |
|---|---|---|---|---|---|---|---|
| PCGMP4 | 7 | -934.3407 | 0 | 1 | 0.9692 | 0.9961 | 0.0355 |
| PCBR4 | 7 | -771.7022 | 162.6386 | $4.824 \times 10^{-36}$ | 0.9628 | 0.9923 | 0.0492 |
| PCLG4 | 7 | -662.0521 | 272.2886 | $7.47 \times 10^{-60}$ | 0.9553 | 0.9878 | 0.0614 |

**Table S7.** Model selection summary and descriptive fidelity metrics across optimal sigmoidal growth formulations. A comparative performance summary validating the selection of the Gompertz mixed-effects model architecture against competing mathematical frameworks. The top-performing configuration from each structural class-reparameterized Gompertz (PCGMP4), Baranyi-Roberts (PCBR4), and Logistic (PCLG4) possessing identical degrees of freedon (fd=7) is evaluated. Selection criteria include the Akaike Information Criterion (AIC), absolute AIC differential (dAIC), Akaike weights (weight), Marginal $R^2$ (variance explained by fixed effects alone), conditional $R^2$ (variance explained by both fixed and random effects), and Root-Mean Square Error (RMSE). The Gompertz framework (PCGMP4) demonstrates definitive descriptive superiority, capturing the raw experimental variations with an Akaike weight of 1.000, maximized explanatory variance (Conditional $R^2$ = 0.9961), and minimized residual error (RMSE = 0.0355).

A)

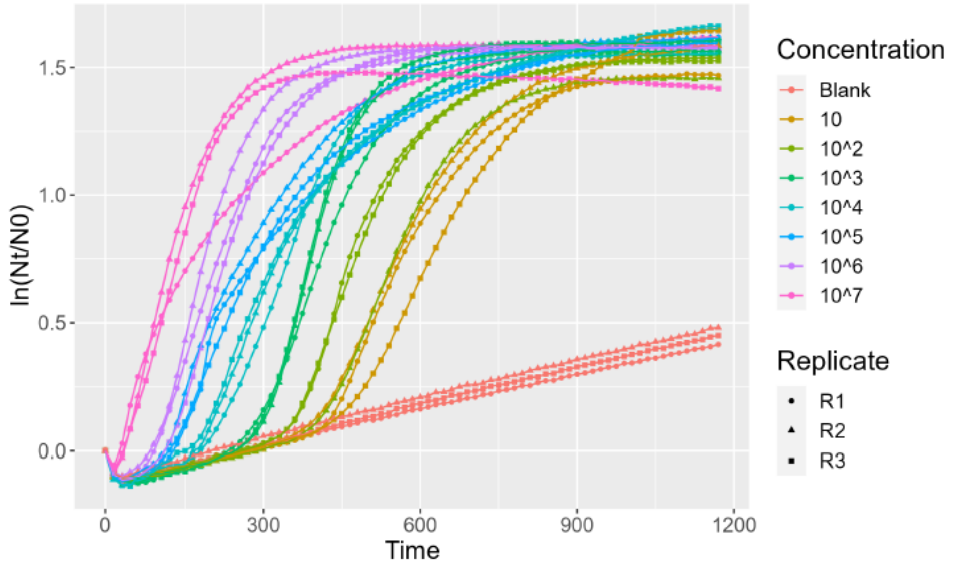


B)

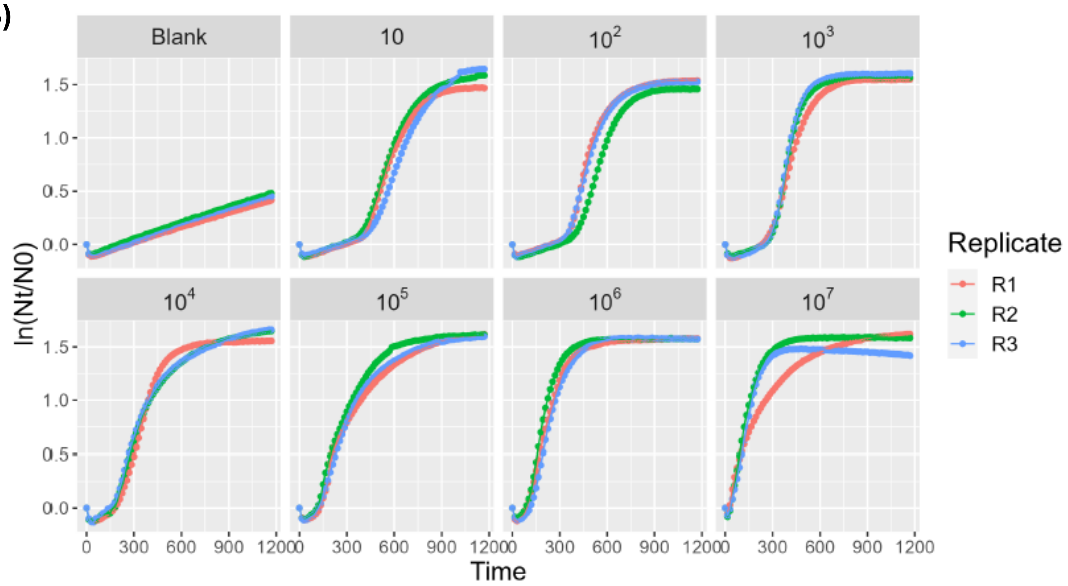


**Figure S49.** Impact of initital bacterial inoculum concentration on real-time kinetic growth trajectories. Comprehensive tracking of log relative population size ln (Nt/N0) over time, demonstrating the mathematical relationship between initial bacterial density and phenotypuc growth parameters. **(A)** Combined growth curves: Overlaid kinetic profiles across all tested concentration ranges ($10^1$ to $10^7$ CFU/mL) alongside an uninoculated hydrogel blank control. The progressive, orderly leftward shift of the sigmoidal curves as concentration increases visually demonstrates the near-perfect negative log-linear correlation with lag phase duration ($\lambda$=-0.991). Conversely, the parallel nature of the exponential slopes highlights that the maximum specidic growth rate ($\mu_m$) remains invariant to the initial concentration load. **(B)** Faceted growth curves: Isolated panel views breaking down each individual initial loading concentration and the control blank across 3 independent technical replicates: R1, R2, R3. The high overlapping symmetry across the individual panels validates the reproducbility of the matrix-bound kinetic modelling framework.

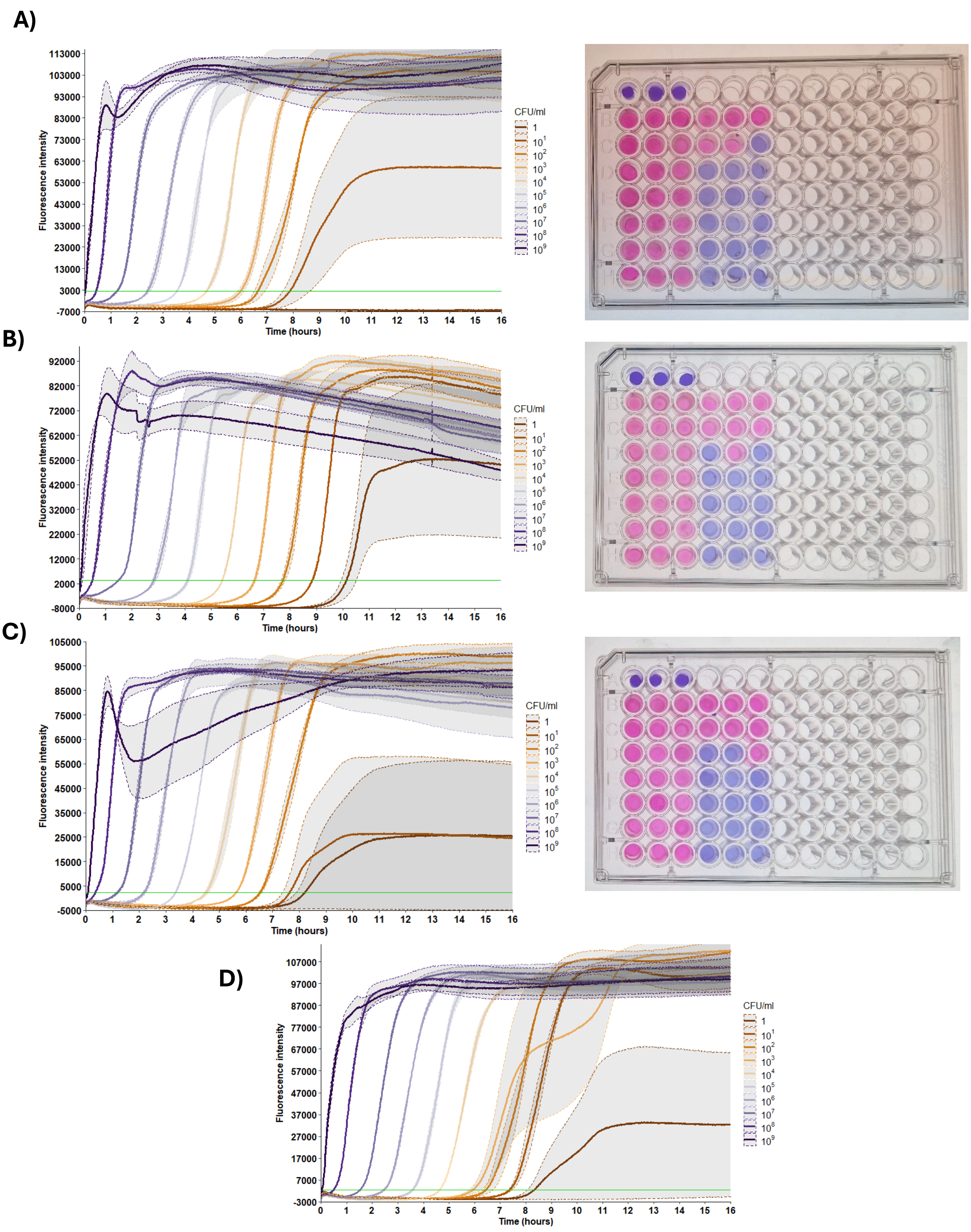


**Figure S50.** Cross-operator validation of multi-species standard dose-response curves in artificial urine. Comprehensive evaluation of platform portability, assay generalizability, and user-invariant analytical performance across an expanded panel of clinical uropathogen isolates. Real-time fluorometric kinetic trajectories (left) and corresponding 96-well colorimetric profiles (right) were indepndently generated using artificial urine spiked with clinical uropathogenic isolates across a 9-log linear dynamic range ($10^{1}$ to $10^{9}$ CFU/mL). **(A)** *E. coli*, **(B)** *P. mirabilis*, **(C)** *ESBL E. coli*, and **(D)** *Klebsiella* species. Shaded regions along the kinetic tracks denote the standard deviation across independent technical replicates.

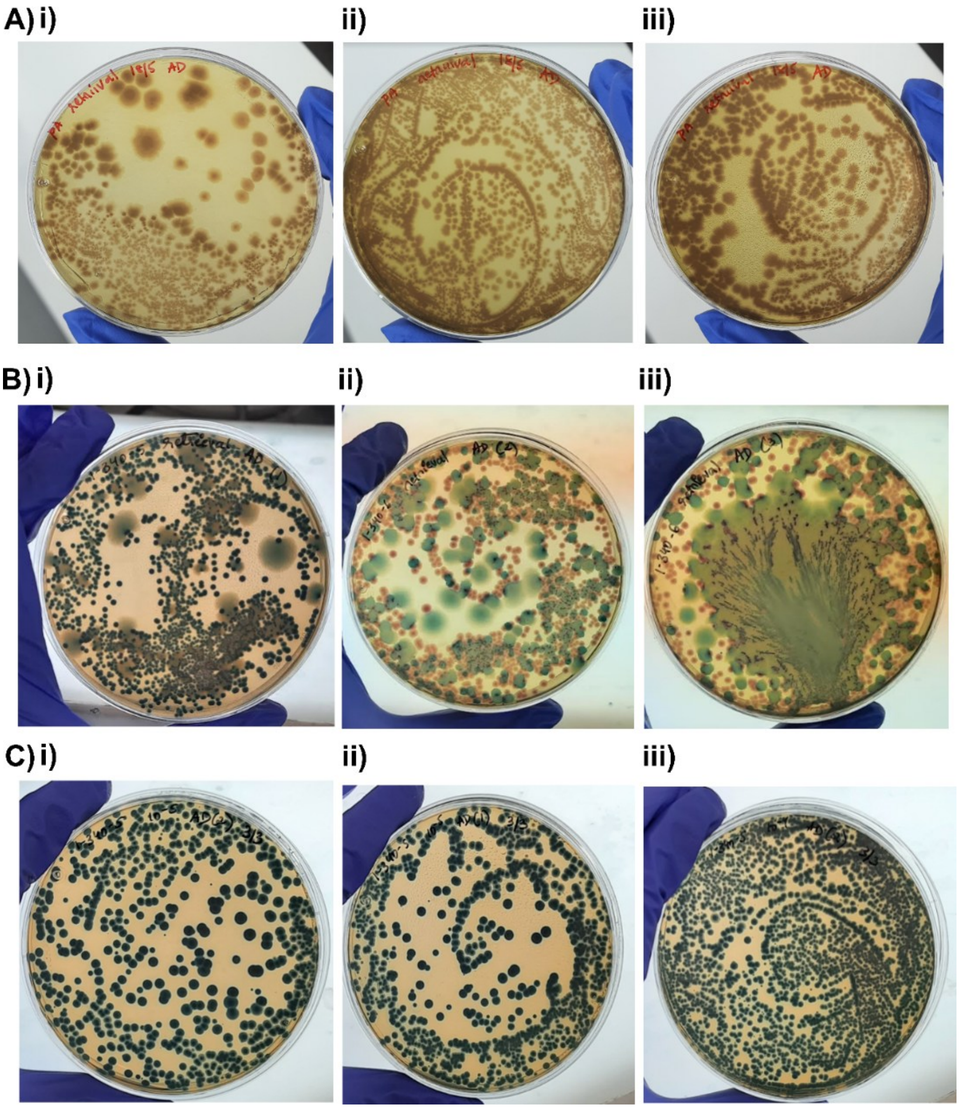


**Figure S52.** Downstream phenotypic testing of clinical samples post-bacterial detection in chromogenic UTI agar: **A)** *Pseudomonas aeruginosa* (sample 54) – brown colonies. Fungal growth was observed in the downstream testing of **B) i)** *Klebsiella* species - small to medium deep blue colonies (Sample 14), fungal species seen as large colony growth with green centre and cloudy opaque edges, **ii)** and **iii)** *E. coli* species – pink colonies (Sample 51), fungal species seen as large colony growth with green centre and cloudy opaque edges in (**ii**) and adundant growth in (**iii**). *Klebsiella* species (Sample 52) – deep blue colonies.

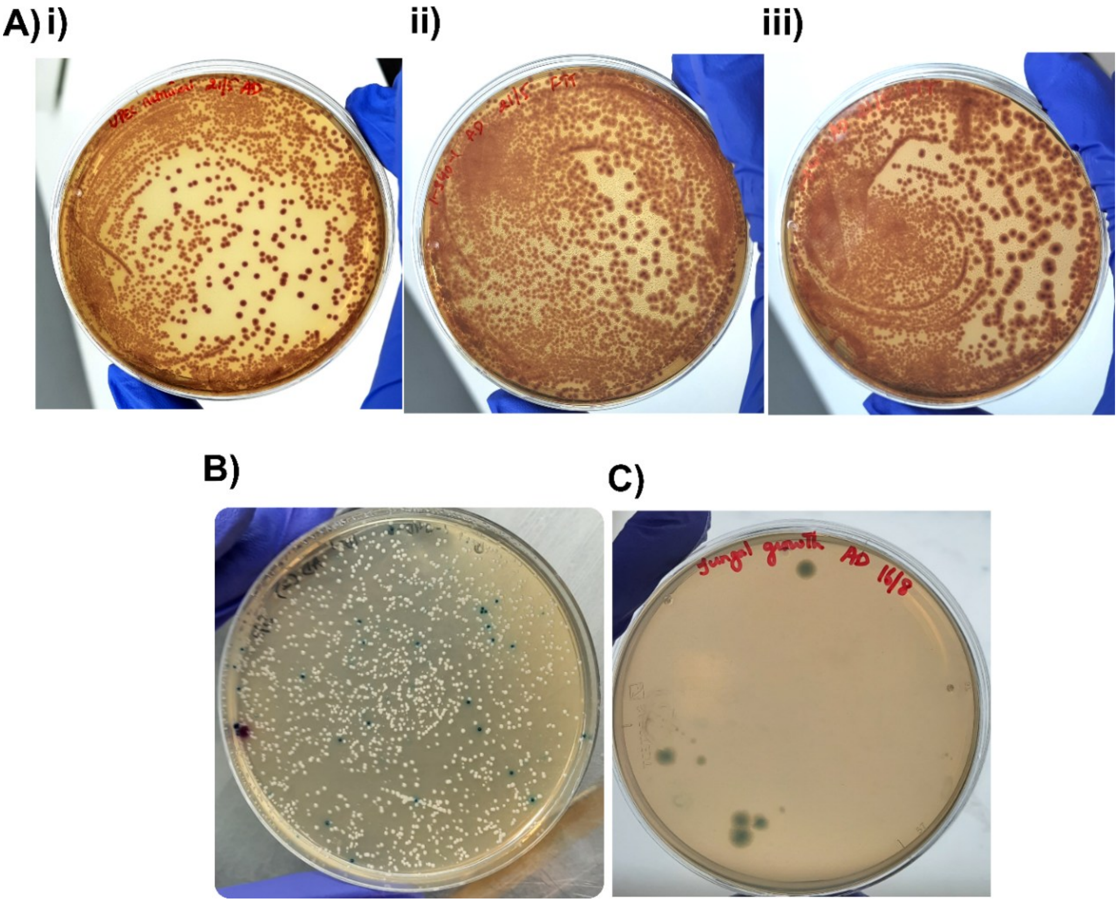


**Figure S53.** Downstream phenotypic testing of clinical samples post-bacterial detection in chromogenic UTI agar: **A)** *E. coli* (sample 49) **(i), (ii),** and **(iii)**. **B)** Mixed bacterial species – *S. aureus* (white) and *Enterococci* (blue-green) (Sample 18) . **C)** Isolation and pure growth of fungal species (from Sample 51) on LB media.
